# Stimulus novelty uncovers coding diversity in survey of visual cortex

**DOI:** 10.1101/2023.02.14.528085

**Authors:** Marina Garrett, Peter Groblewski, Alex Piet, Doug Ollerenshaw, Farzaneh Najafi, Iryna Yavorska, Adam Amster, Corbett Bennett, Michael Buice, Shiella Caldejon, Linzy Casal, Florence D’Orazi, Scott Daniel, Saskia EJ de Vries, Daniel Kapner, Justin Kiggins, Jerome Lecoq, Peter Ledochowitsch, Sahar Manavi, Nicholas Mei, Christopher B. Morrison, Sarah Naylor, Natalia Orlova, Jed Perkins, Nick Ponvert, Clark Roll, Sam Seid, Derric Williams, Allison Williford, Ruweida Ahmed, Daniel Amine, Yazan Billeh, Chris Bowman, Nicholas Cain, Andrew Cho, Tim Dawe, Max Departee, Marie Desoto, David Feng, Sam Gale, Emily Gelfand, Nile Gradis, Conor Grasso, Nicole Hancock, Brian Hu, Ross Hytnen, Xiaoxuan Jia, Tye Johnson, India Kato, Sara Kivikas, Leonard Kuan, Quinn L’Heureux, Sophie Lambert, Arielle Leon, Elizabeth Liang, Fuhui Long, Kyla Mace, Ildefons Magrans de Abril, Chris Mochizuki, Chelsea Nayan, Katherine North, Lydia Ng, Gabriel Koch Ocker, Michael Oliver, Paul Rhoads, Kara Ronellenfitch, Kathryn Schelonka, Josh Sevigny, David Sullivan, Ben Sutton, Jackie Swapp, Thuyanh K Nguyen, Xana Waughman, Joshua Wilkes, Michael Wang, Colin Farrell, Wayne Wakeman, Hongkui Zeng, John Phillips, Stefan Mihalas, Anton Arkhipov, Christof Koch, Shawn R Olsen

**Author notes:** These authors contributed equally to this work (Marina Garrett, Peter Groblewski, Alex Piet, Doug Ollerenshaw, Farzaneh Najafi, Iryna Yavorska). Current address: Georgia Institute of Technology, Atlanta, GA, USA.

## Abstract

Detecting novel stimuli in the environment is critical for learning and survival, yet the neural basis of novelty processing is not understood. To characterize cell type-specific novelty processing, we surveyed the activity of ∼15,000 excitatory and inhibitory neurons in mice performing a visual task with novel and familiar stimuli. Clustering revealed a dozen functional neuron types defined by experience-dependent encoding. Vasoactive-intestinal-peptide (Vip) expressing inhibitory neurons were diverse, encoding novel stimuli, omissions of familiar stimuli, or behavioral features. Distinct Somatostatin (Sst) expressing inhibitory neurons encoded either familiar or novel stimuli. Subsets of excitatory neurons co-clustered with specific Vip or Sst subpopulations, while Sst and Vip inhibitory clusters were non-overlapping. This study establishes that novelty processing is mediated by diverse functional neuron types in the visual cortex.

## Main

The processing and prioritization of novel stimuli is essential to an animal’s survival (*1*). Novelty directs attention, promotes exploration, and triggers learning and memory formation (*2–7*). Neural effects of novelty have been widely documented throughout the brain (*2–4*, *6*). In sensory cortex, novelty boosts population level stimulus responses (*8*, *9*). However single cell measures indicate that novelty effects may be more complex. Both elevated and reduced responses to novel stimuli have been observed (*10–13*), and the dynamics of activity can also be affected (*11*). Further, there are multiple forms of novelty that may involve different cell types (*2–4*, *6*, *7*, *9*, *14–16*). In mice, the effects of both absolutely novel and contextually unexpected stimuli have been reported to differ between cell types including vasoactive intestinal peptide (Vip) and somatostatin (Sst) inhibitory neurons (*9*, *13*, *17–25*).

In modern cell type taxonomies, Vip and Sst inhibitory cells as well as cortical excitatory cells represent distinct “subclasses” that are composed of many finer-grained cell types with distinct gene expression patterns, morphologies, synaptic connectivity, and intrinsic electrical properties (*26–41*). Some cell types are very rare, particularly within the Vip and Sst subclasses, which only comprise ∼5% and ∼2.5% of all neurons, respectively (*29*, *42*, *43*). While some studies have identified functional heterogeneity within Vip and Sst populations (*44–49*), it is unknown whether finer subtypes of Vip, Sst, and excitatory cells are similarly or differentially impacted by novelty and familiarization. The extent of functional heterogeneity has important implications for the computational properties of the local cortical circuitry (*50*, *51*).

Vip and Sst neurons have been shown to powerfully regulate excitatory population activity in different behavioral contexts, as well as during task learning (*52–55*, *55–59*). In the mouse visual cortex, Vip, Sst, and excitatory neurons code not only for visual features, but also for behavioral choices, reward expectation, and locomotor activity (*55*, *60–71*). The extent to which stimulus novelty affects cell type-specific coding for behavior and task-related information remains unknown.

### Open dataset from the Allen Brain Observatory

To broadly characterize novelty processing during behavior in genetically defined cell populations, we used the 2-photon Allen Brain Observatory to perform a survey of activity across nearly 15,000 cells in the mouse visual cortex (Fig. 1A; fig. S1: fig. S2). We imaged cells in transgenic mice expressing GCaMP6 in excitatory, Sst inhibitory, or Vip inhibitory neurons (Fig. 1B; fig. S1B, fig. S2C-E). We collected data from both primary and secondary visual cortical areas across depths spanning layer 2 through upper layer 5 (Fig. 1C; fig. S2). Imaging was performed in mice trained to perform a visually guided task (Fig. 1D; fig. S1D,E; fig. S3). We used longitudinal imaging to track the activity of populations of neurons across multiple sessions with familiar or novel stimuli, alternating between active task performance and passive viewing sessions (Fig. 1C-E; fig. S1F; fig. S4A). Neural activity and animal behavior were monitored throughout each session (Fig. 1F). In this study, we focus on the active behavior sessions to ask how novelty influences encoding of stimulus, behavior, and task information (fig. S4A).

**Figure 1.**
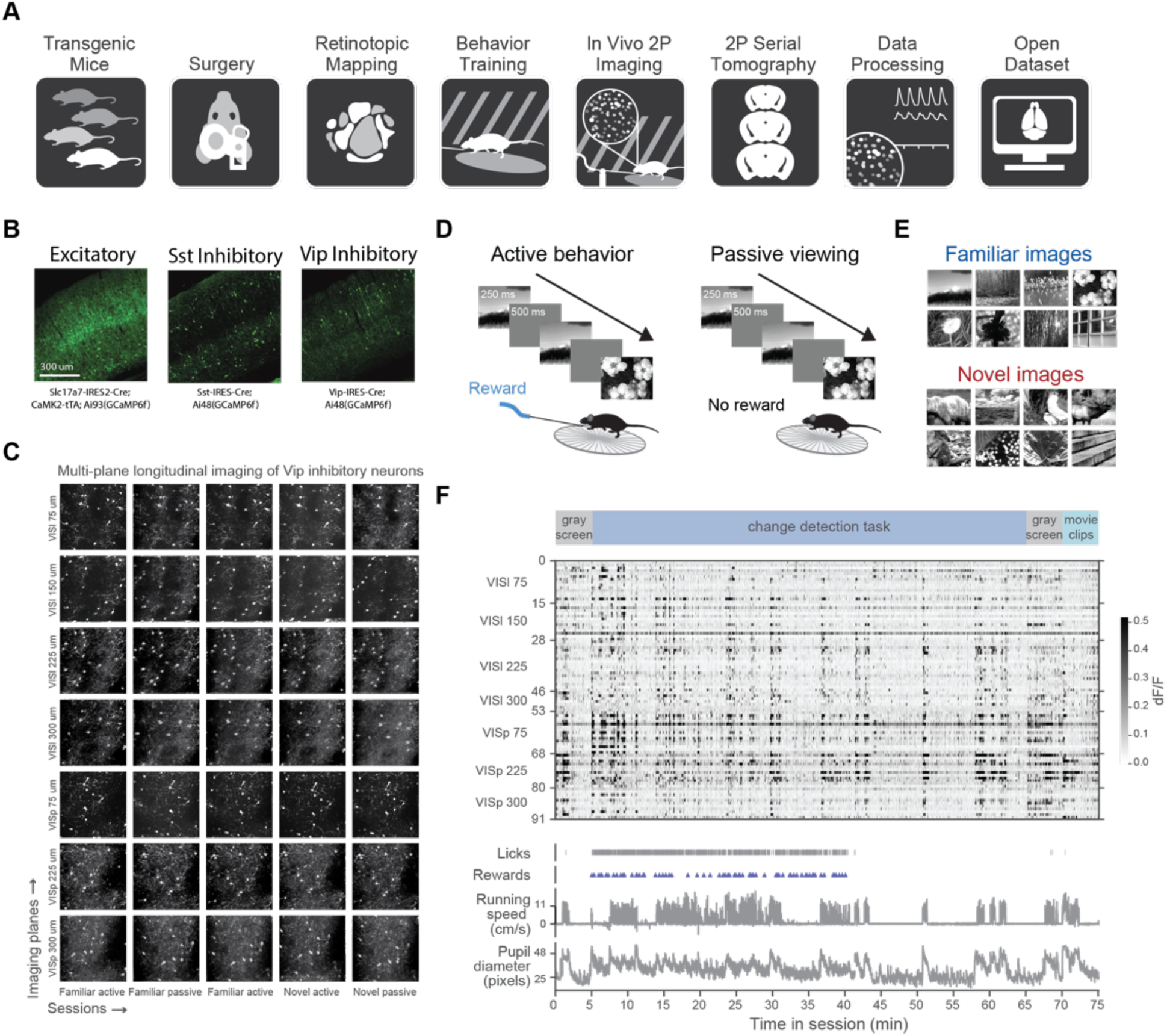
Allen Brain Observatory Visual Behavior 2-photon dataset. **(A)** Data collection pipeline, illustrating steps used to collect and process standardized in vivo physiology during behavior in transgenic mice. **(B)** The genetically encoded calcium indicator GCaMP6fwas expressed in transgenic mice labeling excitatory (Slc17a7-IRES2-Cre;CamK2-tTA;Ai93(GCaMP6f)), Sst inhibitory (Sst-lRES-Cre;Ai148(GCaMP6f)), or Vip inhibitory neurons (Vip-lRES-Cre; Ai148(GCaMP6f)). Images show GCaMP fluorescence in coronal sections from these transgenic lines. **(C)** in vivo 2-photon calcium imaging was performed in the visual cortex of well-trained mice at depths between 75-375 um from the cortical surface. Up to 8 fields of view were recorded in each imaging session, and the same populations of neurons was followed over multiple days of imaging. Images show maximum intensity projections for all recordings from one mouse. **(D)** Mice experienced different session types on different imaging days. Sessions alternated between active task performance and passive viewing sessions, during which mice were satiated and stimuli were displayed in the absence of reward. **(E)** During each imaging session, mice viewed either familiar images (viewed thousands of times during training) or novel images (seen for the first time under the microscope). Each image set was repeated over multiple days, as shown in panel C. **(F)** Top row, each 2-photon imaging session began by recording spontaneous activity in the presence of a gray screen for 5 minutes, followed by 60 minutes of either active task performance or passive viewing, ending in another 5-minute gray screen period and 10 repeats of a 30 second movie clip. Middle panel, calcium fluorescence was measured and baseline normalized to produce dF/F traces. Y-axis shows cumulative number of neurons for each visual area and imaging depth (in um) recorded in this session. Bottom panels, animal behavior, including running speed, pupil diameter, and lick times were recorded concurrently with neural activity, and aligned to the timing of stimulus events. All data is openly available at brain-map.org/explore/circuits/visual-behavior-2p.

Our experimental design includes several additional features to enable the study of neural coding and behavior. The visual task requires that mice detect unexpected image changes (oddballs) that differ from the recent stimulus context but are not novel in an absolute sense (Fig. 2A; fig. S3A) (*15*). We also include unexpected stimulus omissions (Fig. 2B; fig. S3A), which are another form of contextual surprise distinct from stimulus novelty (*14*). Mice learn the task through a multi-stage training curriculum with increasing levels of difficulty (fig. S3; also see (*72*) for detailed analysis of behavior). All data, including 703 physiology and 4,782 training sessions from 107 mice, are publicly available for further analysis (portal.brain-map.org/circuits-behavior/visual-behavior-2p; see Supplemental Text for detailed description of dataset).

**Figure 2.**
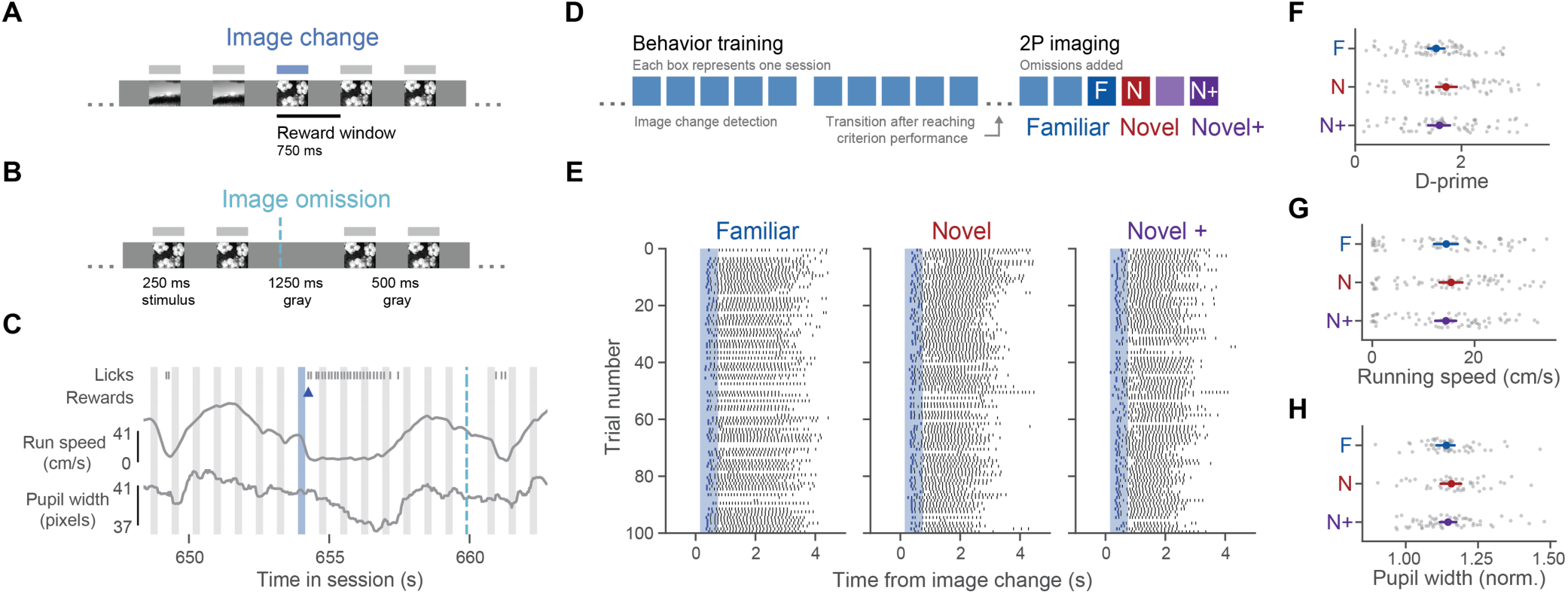
Visual change detection task with familiar and novel images. **(A)** In the go/no-go change detection task, natural scene images were displayed for 250ms, followed by a 500ms gray screen period. Images were repeated a variable number of times until the identity of the image changed. Mice learned to lick within 750ms of an image change to earn a reward. (B) Once mice were well trained, 5% of non-change images were randomly omitted. (C) During each session, animal behavior was measured, including licks, running speed, and pupil diameter, along with the timing of rewards. Gray bars correspond to repeated image presentations, blue bars are image changes, dashed cyan lines indicate the timing of stimulus omissions. (D) Mice learned the task with a set of natural scene images that became highly familiar over weeks (see Supplemental Text for description of training procedure). After reaching performance criteria, mice transitioned to 2-photon imaging sessions, during which they performed the task with familiar images over multiple sessions, followed by several repeated sessions with novel images. Three sessions were selected for subsequent analysis: the first session with novel images (Novel), a subsequent active behavior session with novel images (Novel÷), and a behavior session with familiar images prior to the first novel session (Familiar). Passive viewing sessions that were interleaved between active behavior sessions were excluded in this study. (E) Licking aligned to image change times for the first 100 trials across three sessions for one mouse. (F) Behavior performance across mice (n=66, gray points), measured as d-prime, for each session type (Familiar, Novel, Novel÷). Colored points show average of all mice in each session type +/- 95% confidence interval. (G) Average running speed across mice and sessions. (H) Average pupil diameter across mice and sessions, normalized to the average pupil size in the gray screen period before task performance within each session. Statistics for panels F-G were performed by one-way ANOVA across experience levels followed by Tukey HSD to correct for multiple comparisons (p<0.05).

### Change detection task with familiar and novel images

Mice performed a go/no-go visual change detection task in which they viewed a continuous series of briefly presented images (250 ms duration) interleaved with a gray screen (500 ms) (Fig. 2A) (*72*, *73*). The same image was repeated a variable number of times until the image identity changed (fig. S3A,C,D). Mice learned to lick a reward spout within 750 ms after detecting the image change (fig. S3B, E). Licks outside the reward window delayed the onset of the next trial (fig. S3A). Once mice were well trained, we introduced rare, unexpected stimulus omissions (5% of non-change image presentations) (Fig. 2B). These omissions violated the learned expectation of stimulus timing (fig. S3A). Mice typically did not lick during omissions but were more likely to false alarm to the image presentation following an omission (fig. S3B) (*72*). We measured licking, reward times, running speed, and pupil diameter in each session (Fig. 2C).

Mice learned the task with a set of eight natural scene images that became highly familiar during training (Fig. 2D; Fig 1E; fig. S1f; fig. S2A,B; fig. S3E-J). Each image was shown hundreds of times within each 1-hour session (∼550 presentations per image per session). During GCaMP imaging, mice first performed the task with familiar images over multiple days; we refer to these as “Familiar” image sessions (Fig. 2D; fig. S4A). Subsequently, mice performed the task with a new image set they had never seen before. We refer to the first session with novel images as the “Novel” session. We refer to follow-up sessions with the novel image set as “Novel+” sessions, since the absolute novelty of these images is extinguished upon repeated exposure. The images in the Novel+ behavior sessions differ from Familiar sessions by an order of magnitude fewer exposures (27+/-15 sessions with familiar images prior to Familiar sessions, 2+/-1 session with novel images prior to Novel+ behavior sessions; fig. S4A-C; fig. S3G-I;).

Mice generalized their behavior to novel images, performing the task similarly in the Familiar, Novel, and Novel+ sessions (Fig. 2E,F; fig. S3E,J; fig. S4E-H). This indicates that mice had learned the task rules, and that performance was not disrupted by the introduction of novel stimuli. Other measures of animal behavior were also consistent across familiar and novel sessions, including lick rate, running speed, and pupil width (Fig. 2G,H; fig. S5).

### Impact of novelty on excitatory, Vip, and Sst subclasses

We analyzed responses of each recorded neuronal population across multiple days of change detection task performance with familiar or novel images (12,826 excitatory cells, 468 Sst cells, 1,197 Vip cells; Fig. 3A; fig. S4I-J). For each mouse we identified three sessions for analysis: a session with the highly familiar image set (Familiar), the first session with novel images (Novel), and a subsequent behavior session with the novel image set (Novel+) to follow the familiarization process in the same cells (fig. S4A). These selection criteria resulted in 202 imaging sessions from 134 fields of view in 66 mice (fig. S4I-J). Calcium fluorescence was deconvolved and all analysis was performed on the timing and magnitude of calcium events (Fig. 3A) (*74*). A subset of neurons were detected and matched across all three session types (3,306 Excitatory cells, 200 Sst cells, 411 Vip cells; fig S6; fig. S4J).

**Figure 3.**
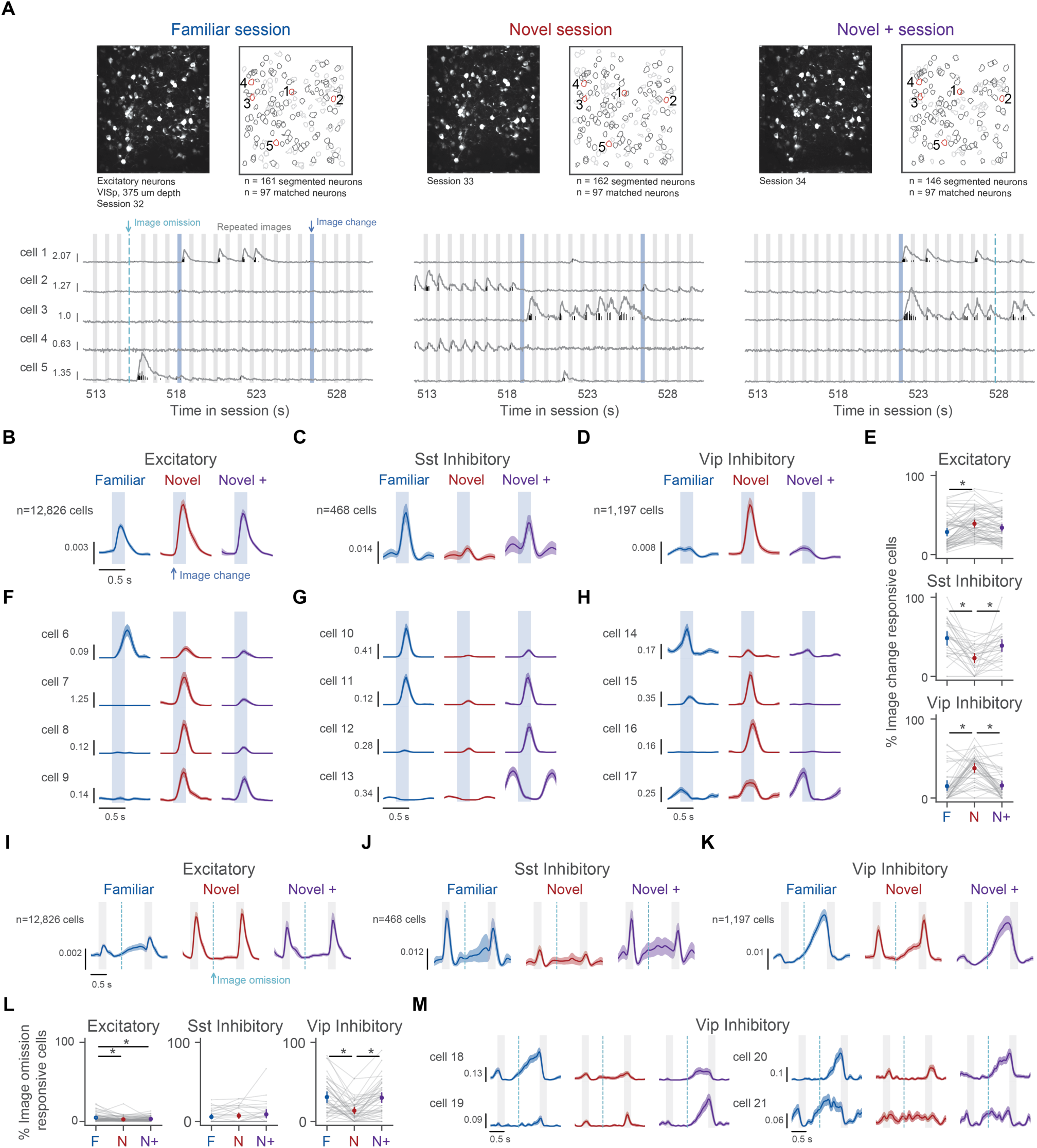
Stimulus novelty alters image change and image omission activity in Excitatory, Sst inhibitory, and Vip inhibitory neurons. **(A)** Populations of neurons were imaged over multiple sessions with familiar and novel images. Top row, panels show imaged neurons as maximum intensity projection (left) and segmented regions of interest (ROls) corresponding to neuronal cell bodies (right). ROls outlined in black were active, segmented, and matched in all 3 sessions. ROls outlined in gray were present in only one or two sessions. ROls outlined in red correspond to the cells whose activity is displayed in bottom row. Bottom row, panels show fluorescence signals measured from 5 example neurons that were matched across sessions. Gray traces are normalized fluorescence values, black lines show the timing and magnitude of deconvolved calcium events. Scale bar shows magnitude of detected calcium events, in units of dF/F. **(B-D)** Population average responses aligned to image changes for all excitatory (n= 34 mice, 12,826 neurons), Sst inhibitory (n= 15 mice, 468 neurons), and Vip inhibitory neurons (n= 17 mice, 1,197 neurons). Y-axis scale shows averaged magnitude of detected calcium events, in units of dF/F. **(E)** Fraction of image change responsive cells for each session (Familiar, Novel, Novel+) for each cell type. Responsiveness is defined as having significantly higher response during image changes compared to the gray screen period at the start of each session. **(F-H)** Average image change evoked responses across sessions for example neurons from Excitatory, Sst inhibitory, and Vip inhibitory cell types. **(1-K)** Population average responses aligned to image omissions for each cell population. Color & scale bar conventions are the same for all panels. **(L)** Fraction of omission responsive cells for each session (Familiar, Novel, Novel+) for each cell type. Responsiveness is defined as having significantly higher response during image omissions compared to the gray screen period at the start of each session. **(M)** Average omission aligned response for 4 example Vip inhibitory neurons.

Novelty differentially impacted excitatory, Sst inhibitory, and Vip inhibitory population activity. Excitatory population responses to image changes were larger in the Novel session compared to both Familiar and Novel+ sessions (Fig. 3B; fig. S7A), and the ability to decode image changes was also enhanced during the Novel session (fig. S7G,H). In contrast, Sst population activity was suppressed in the Novel relative to Familiar and Novel+ sessions (Fig. 3C; fig. S7A). Vip population activity resembled the excitatory population in the Novel session, with strong image evoked responses and enhanced decoding of image changes (Fig. 3D; fig.S7A). However, Vip activity was image-suppressed in both the Familiar and Novel+ sessions. The proportion of neurons responding to image changes in each session reflected these results; a larger fraction of excitatory and Vip neurons were active during Novel sessions, while Sst cells were more responsive during Familiar and Novel + sessions (Fig. 3E; fig. S7E-F).

When examining single cells response profiles, we observed their activity often deviated from the population average (Fig 3F-H). For instance, some excitatory neurons were highly selective for familiar stimuli, whereas others were driven by novel images (Fig. 3F, fig. S7M,N). Among novelty preferring cells, some showed comparable responses during Novel and Novel+ sessions, whereas others only responded during the Novel session. Among the Sst population, some neurons preferred either Familiar or Novel+ sessions, but others responded in both (Fig. 3G; fig. S7M,N). Many Vip cells had enhanced image change responses in the Novel session, but were silent during Familiar and Novel + sessions (Fig. 3H; fig. S7M,N). Others showed ramping activity prior to stimulus onset, or, more rarely, responded to familiar image changes.

For each cell we quantified the strength of the image change response (fig. S7A-F), the difference between change and pre-change responses (fig. S7I-L), the modulation of these responses by novelty (fig. S7M,N), and the dependence of activity on the identity of the image set used during training (fig. S7B-D). The averages of these distributions confirmed the biases across the populations of excitatory, Vip, and Sst cells; on average, excitatory and Vip populations are elevated by novelty but the Sst population is suppressed. However, we observed a broad distribution of values across individual neurons (fig. S7J-N), reflecting substantial heterogeneity across cells even within the same genetic subclass.

### Omission signals in Vip cells emerge with familiarization

Previously we demonstrated that Vip inhibitory neurons have strong ramping activity following the unexpected omission of familiar but not novel images (*18*). However, the time course of familiarization leading to the emergence of this ramping activity was left unresolved. Whether omission ramping occurs in Sst cells is also unknown.

We first validated our previous observation that ramping activity in Vip neurons is high in Familiar sessions but lower in Novel sessions (Fig. 3K; fig. S8A-F). Next, we examined activity in the repeated Novel+ session, as novel stimuli became more familiar, and found that omission ramping activity had re-emerged after just ∼1-2 days of stimulus exposure (Fig. 3K; fig. S8A-F). The fraction of omission responsive Vip cells (Fig. 3L; fig. S8K) and the ability to decode omission signals from population activity (fig. S8G-H) also increased in the Novel+ versus Novel session. These results suggest that the familiarization process whereby Vip neurons develop omission activity requires only just a few 1-hour sessions of stimulus exposure.

In individual Vip cells matched across sessions, omission activity was typically larger in the Familiar and Novel+ sessions compared to Novel session (Fig. 3M; fig S9L,M). However, in contrast to the population average, we observed clear heterogeneity across cells; some cells showed ramping primarily in Familiar sessions, others were specific to Novel+ sessions, and other cells were similarly active in both sessions. The time course of ramping activity also differed; some cells began to ramp prior to the expected stimulus onset time while others only increased activity after the image was omitted (Fig. 3M).

Omission related activity in the excitatory and Sst populations was weak on average (Fig. 3I,J,L; fig. S8), although we observed a small fraction of non-Vip cells with elevated activity following image omissions (Fig. 3L; fig. S8K). Average Sst omission related activity was typically of a similar magnitude across experience levels (fig. S8). Excitatory responses during omissions, while rare, were typically larger in the Familiar session compared to Novel and Novel+ sessions (Fig. 3I,L; fig. S8), suggesting that extensive familiarization is required for omission signals to emerge in these cells.

Several control analyses further demonstrated that omission signals are associated with familiarization, rather than differences in behavior, task engagement, or image features (fig. S9).

### Encoding of stimulus and behavior

In our experiments, stimulus changes and omissions occur concurrently with other task events including animal choices, rewards, and movements, all of which can influence neural activity. To quantify the contributions of these factors and ask whether they are changed with novelty, we used a linear regression model with time-dependent kernels to predict the activity of each cell in each session (Fig. 4A-C; fig. S10A-C) (*63*, *70*, *75*, *76*). Features used to predict neural activity included stimuli (8 images), omissions, task events (hits and misses), and behavioral factors including licking, running speed, and pupil diameter (Fig. 4A; fig. S10A). We computed the unique contribution of each feature by comparing the performance of the full model with reduced models fit without each feature or group of features (Fig. 4C; Fig. S10D). We refer to this fractional reduction in variance explained as the “coding score” (Fig. 4D). Kernels and coding scores for all features across all cells are shown in figs. S11-13.

**Figure 4.**
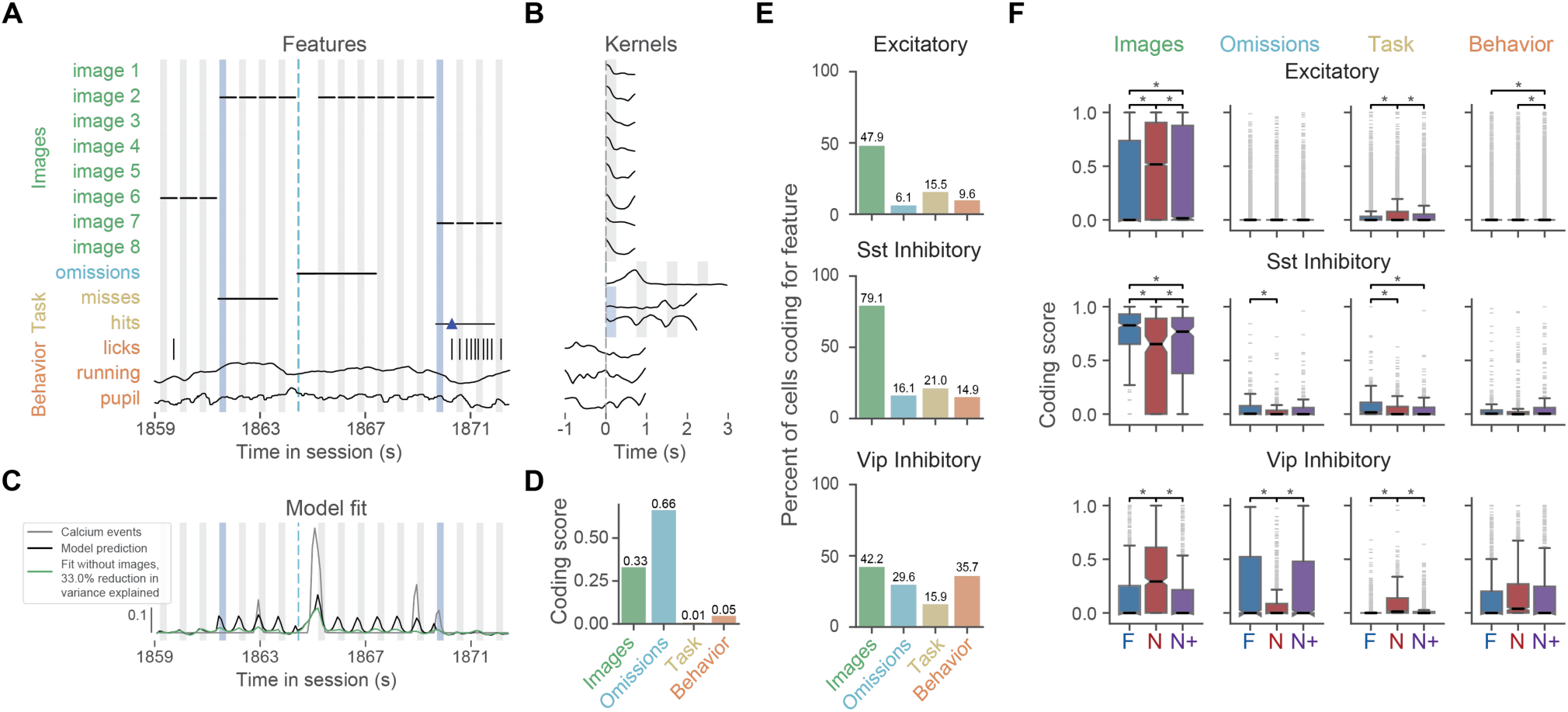
Encoding of stimulus and behavior using kernel regression models. **(A)** Features included in regression model. Gray vertical bars are repeated stimulus presentations, blue vertical bars are image changes, dashed light blue lines are image omissions. Features are shown as rows. Color of feature name on y-axis indicates the feature group it belongs to (images in green, omissions in blue, task in yellow, behavior in orange). For event-aligned kernels (images, omissions and task features), black horizontal lines indicate the time window covered by the kernel for that feature, relative to the event time. Blue triangle indicates time of reward delivery. Vertical black tick marks indicate lick times; kernel window for lick times is +/-1 second around each lick. For continuous features (running and pupil), the full timeseries is shown. Kernels for continuous features cover +/-1 second around each timepoint. (B) Kernel weights for one example cell. Positive weights indicate an increase in cell activity relative to the start of the kernel window, negative weights indicate a decrease. Y-scale of kernel weights are not matched across features. (C) Model fit for example cell. Vertical bars and shading are as in panel A. Gray trace shows detected calcium events, black trace shows model fit for full model with all features included. Green trace shows model fit when image features are removed. The fractional reduction in variance explained after removing a feature or group of features is called the “coding score”. (D) Coding scores for example cell, representing the fractional change in variance explained when each feature group is withheld from the model. (E) Percent of cells encoding each feature group across cell types. A cell is considered as coding for a feature if the coding score for that feature is greater than 0.1. (F) Distribution of coding scores for each feature group (columns) across cell types (rows). Colors indicate session type (Familiar, Novel, Novel÷). Statistics performed by one-way ANOVA followed by Tukey HSD across experience levels (p<0.05).

We grouped features into four categories: images, omissions, task, and behavior information, and examined how the different cell populations coded for these feature groups (Fig. 4A,E; see fig. S10D for ungrouped features). The majority of excitatory and Sst cells primarily encoded images, whereas Vip cells encoded a combination of images, omissions, behavior, and task features (Fig. 4E).

To investigate how novelty influenced neural coding, we split the data by experience level (Familiar, Novel, Novel+) and evaluated coding scores across sessions (Fig. 4F; fig. S14, S15). On average, excitatory and Vip populations had stronger image coding during the Novel session, while Sst cells had reduced image coding. Image coding across the Vip population was not significantly different between the Familiar and Novel+ sessions. In contrast, excitatory and Sst image coding in Novel+ sessions did not fully return to the same level as in Familiar sessions. These results suggest a faster timescale of familiarization in the Vip compared to excitatory and Sst populations.

As expected based on population averages, omission coding in the Vip population was reduced in the Novel session but re-emerged in the subsequent Novel+ session (Fig. 4F; fig. S14). Omission coding in excitatory and Sst cells was weak overall (Fig 4F, fig. S14), although a small subset of these cells coded omissions (fig. S11B) and there was a slight enhancement of omission coding by Sst cells in the Familiar session (Fig. 3F; fig. S14).

Recent studies have characterized behavioral and task related coding in the visual cortex (*63*, *65*, *70*), but it is unknown whether these properties are influenced by stimulus novelty. We found that Vip and Excitatory populations have elevated coding for task features (hits and misses) in the Novel session, while Sst neurons have higher task coding during Familiar sessions (Fig. 4F; fig. S14, S15). This suggests that task information is differentially encoded across cell populations depending on the level of experience with task relevant stimuli. In contrast, coding for behavior features (licking, running, pupil) was not changed in the Novel versus Familiar sessions in any of the cell populations (Fig. 4F; fig. S14).

These results were consistently observed when using ungrouped model features (fig. S14A), when limiting analysis to the subset of cells that were tracked across all 3 experience levels (fig. S14A,B), and when stricter cell selection criteria were used (fig. S14B).

Although we found that novelty related encoding differed significantly between excitatory, Sst, and Vip cells on average, the distributions of coding scores were often wide (Fig. 4F), and examination of single cells indicated substantial heterogeneity within each genetic subclass (fig. S11-13). To evaluate this heterogeneity for underlying structure, we next applied unsupervised clustering to the single cell coding scores.

### Experience-dependent coding clusters

To evaluate how single cells altered their coding properties with novelty and familiarization, and ask whether these properties were shared across subsets of neurons, we used unsupervised clustering on the cross-session normalized coding scores of all cells matched across days (Fig. 5A-C; fig. S16). We identified 12 clusters with distinct patterns of experience-dependent coding (Fig. 5C; fig. S17A-C). These clusters had high within cluster correlations (fig. S16C) and unique response properties that were consistent with their patterns of experience-dependent coding (Fig. 5D,E; fig. S17D-F). We further validated the clusters by comparing cluster distributions to those generated using shuffled data (fig. S18) and familiar-only sessions (fig. S19), demonstrating that the presence of novel stimuli resulted in more diverse response types and more unique clusters.

**Figure 5.**
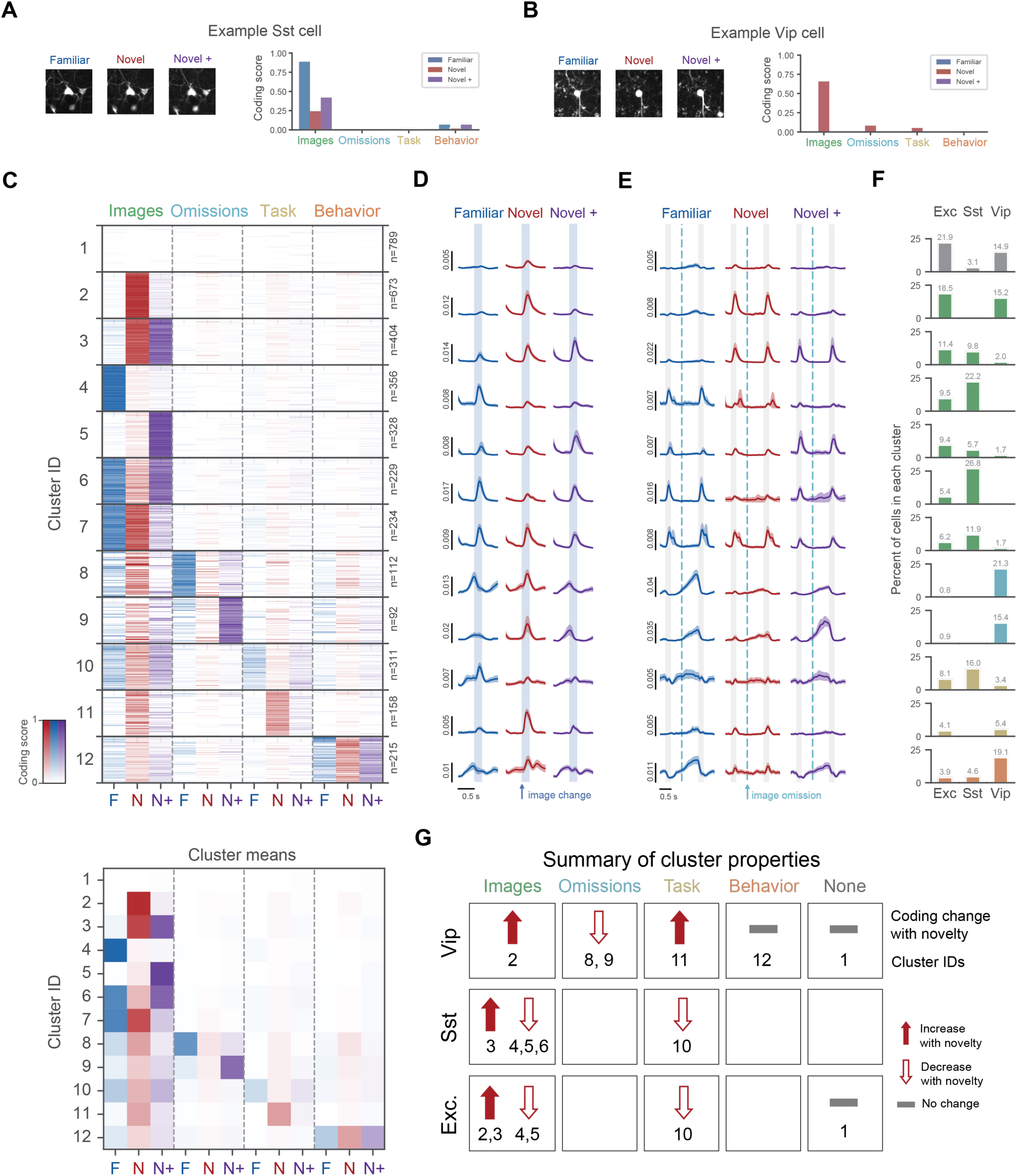
Experience dependent coding diversity of excitatory and inhibitory populations. **(A)** Coding scores across experience levels for example Sst cell. **(B)** Coding scores across experience levels for example Vip cell. **(C)** Top, clustered coding scores for all matched cells. Clustering was performed on all cell types. Each row of the heatmap is one cell, columns are features across experience levels. Number of cells per cluster is shown on right side of plot. Color intensity represents strength of coding score. Bottom, average coding scores for each cluster. **(D)** Average image change evoked response for each cluster, separated by experience level (Familiar, Novel, Novel+). (E) Average omission evoked response for each cluster across experience levels. **(F)** Distribution of cells across clusters for each cell type. Colors indicate the preferred feature encoding of each cluster (images in green, omissions in blue, task in yellow, behavior in orange, non-coding in gray). (G) Summary of cluster properties for the 6 largest clusters within each cell type. Rows are cell types, columns indicate the preferred feature coding of the clusters within each box. Upward arrows indicate an increase in codin1g0strength with novelty, downward arrows indicate a decrease in coding strength with novelty. Gray horizontal line indicates no change with novelty. Numbers under arrows are cluster IDs.

Most clusters showed some degree of image coding during at least one of the sessions (Familiar, Novel, or Novel+; Fig. 5C; fig. S17). The exception was a non-coding cluster that did not encode any features used in our regression model (cluster 1). Six of the clusters selectively encoded images but did not strongly encode omissions, behavior, or task features (clusters 2-7). These clusters were differentiated from one another based on whether they encoded images in the Familiar, Novel, or Novel+ session, or a combination of these (e.g., cluster 2 encoded images only in the Novel session, cluster 8 encoded images in the Familiar and Novel + sessions).

Two clusters encoded image omissions, and were distinguished by whether they encoded omissions in the Familiar or Novel+ session (Fig. 5C; fig. S17; clusters 8 and 9). Task encoding was also present in two clusters; one with elevated task coding in the Novel session and another preferring the Familiar session (clusters 11 and 10). Finally, behavioral features were represented by a single cluster whose coding properties were not modulated by novelty (cluster 12).

We computed the average image change and image omission aligned responses of cells belonging to each cluster across experience levels (Fig. 5D,E; fig. S17D-F), confirming each cluster’s pattern of experience-dependent image encoding. This further revealed distinct activity dynamics across the clusters. For instance, both omission coding (clusters 8 and 9) and behavior coding (cluster 12) clusters had elevated activity in the gray screen period between stimulus presentations, but they differed in their temporal dynamics (Fig. 5D,E; fig. S17D-F). Notably, the clusters selectively encoding familiar images (cluster 4) showed elevated baseline activity prior to image onset in the Familiar session compared to sessions with novel images (Fig. 5D-E; fig. S17D-F).

### Coding diversity within genetic subclasses

Next we asked how each cell type contributed to the clusters we identified. Cells within the excitatory, Vip, and Sst populations were not evenly distributed across clusters, demonstrating diversity both within and across the genetically defined subclasses (Fig. 5F; clusters split by subclass further described in figs. S20-22). Sst and Vip neurons often co-clustered with excitatory cells, indicating shared coding properties between excitatory and inhibitory subpopulations. In contrast, the two inhibitory subclasses rarely shared coding properties with each other and instead fell into distinct, non-overlapping sets of clusters (Fig. 5F).

Most image coding clusters were shared between excitatory and Sst cells (clusters 3-7), but the distributions showed different biases. Sst cells primarily participated in clusters encoding familiar images (clusters 4 and 6), while the largest clusters within the excitatory population had enhanced encoding of novel images (clusters 2 and 3), consistent with population averages (Figs. 3,4). However, we also found a subset of excitatory cells that were highly responsive to familiar images (cluster 4) and a subset of Sst cells responsive to novel images (cluster 3)—these represent smaller, specialized subpopulations that are easily obscured by population level measures.

Clusters shared by excitatory and Vip neurons included one image coding cluster and one task coding cluster, both selective for the first novel session (clusters 2 and 11). Omission coding clusters were strongly biased to Vip neurons (clusters 8 and 9), however a small fraction of excitatory cells also encoded omissions (<2% excitatory cells in clusters 8 and 9).

A few clusters included all three cell populations, to varying degrees. The subset of cells with elevated task coding in the Familiar session (cluster 10) primarily consisted of Sst neurons, with smaller fractions of excitatory and Vip cells. Behavior coding (cluster 12) was most prevalent among Vip cells but was also present among excitatory and Sst neurons.

These results demonstrate that: 1) excitatory, Sst, and Vip inhibitory neurons can be further subdivided into multiple functional subpopulations with distinct patterns of experience-dependent coding, 2) there are shared patterns of coding across excitatory and inhibitory subpopulations, and 3) Sst and Vip inhibitory subpopulations are typically mutually exclusive. The preferred feature coding and direction of novelty induced coding changes for the 6 largest clusters within each genetically defined subclass is summarized in Figure 5G.

## Discussion

By tracking large populations of genetically labeled cells over repeated behavior sessions with familiar and novel images, we revealed unappreciated diversity in the influence of novelty in the visual cortex. We identified distinct clusters of cells whose encoding properties were elevated, suppressed, or unaltered by novelty. This novelty modulation occurred in combination with the encoding of stimuli, omissions, task, or behavior features. Like previous studies (*77*, *78*), we find that the neurons we characterized are not generic “novelty detectors” that respond to any novel stimulus; rather, they combine feature encoding with experience in specific ways. Such multiplexing could support integration of temporal familiarity with sensory and task representations to guide decision-making and learning (*65*, *79*, *80*).

In all genetic subclasses measured in this study—excitatory, Vip, Sst—we observed substantial within-subclass functional heterogeneity, demonstrating that these subclasses cannot be treated as functionally homogenous populations. This finding might be expected given the known molecular diversity of these neurons. Single cell transcriptomics indicates there exists over a dozen transcriptomic-types (t-types) of both Vip and Sst cells (*26*, *27*, *33*, *36*, *37*, *81*). T-types often correspond to subpopulations with distinct morphologies, connectivity profiles, and electrophysiological properties, which can shape their role in circuit function (*28*, *30*, *34*, *35*, *37*, *39*). Accordingly, the novelty-based response types we identified could correspond to distinct t-types. Future work can test this idea using 2-photon imaging followed by retrospective spatial transcriptomics (*82*, *83*).

The heterogenous functional properties we observe could also reflect distinct synaptic dynamics and plasticity mechanisms (*32*, *48*, *84–87*). Indeed, Aitken et al. (*85*) demonstrated that a neural network model with local, unsupervised synaptic learning rules can recapitulate some of the within-subclass functional diversity we observe experimentally. In this model, within-subclass heterogeneity is not built-in but rather emerges through activity-dependent modulation of synaptic weights. There is experimental evidence for this possibility as well; for instance, a subset of Vip neurons with highly dynamic spines (*87*) could enable rapid changes in coding and dynamics following novelty exposure.

Neuromodulation is likely to be a key mediator for generating the diversity of novelty-related coding changes we observe (*3*, *88–90*). Novelty strongly engages multiple neuromodulatory systems (*3*, *6*, *91–96*), and neuromodulators can regulate cell excitability and synaptic plasticity (*97*). Different t-types express distinct patterns of neuromodulator receptors (*88*) which could provide a rich substrate for within-subclass heterogeneity of novelty effects.

On a population level, the finding that novel stimuli produce elevated responses in both Vip and excitatory cells but reduced responses in Sst cells is consistent with the canonical Vip-Sst-excitatory disinhibitory circuit in the cortex (*17*, *43*, *52*, *57*, *98*). Our result that Vip and Sst clusters are largely mutually exclusive is also consistent with the known mutual inhibition between (*53–56*, *99–101*). However, the diversity of clusters we observe within these subclasses suggests that multiple Vip-Sst subnetworks operate in parallel to mediate novelty processing and familiarization, along with other circuit motifs. Subnetworks formed via interactions of neurons belonging to different experience dependent coding clusters could serve distinct roles, including the initial detection of novelty, gating plasticity, forming stimulus-reward associations, or stabilizing learned sensory representations (*51*, *52*, *84*, *97*, *98*, *102–105*).

Importantly, this study establishes novelty and familiarization as key drivers of cellular functional diversity in the visual cortex. Our results provide a rich substrate for future work linking diversity in novelty coding to cell type identity, synaptic connectivity, and models of circuit dynamics.

## Acknowledgments

We thank the Allen Institute founder, Paul G. Allen, for his vision, encouragement, and support. We thank the Allen Institute Laboratory Animal Services team for assistance with tissue collection, the Imaging team for assistance with imaging histology sections, the Transgenic Colony Management team for generating and maintaining the mouse lines, and the Animal Care Team for taking care of our mice. We thank Karel Svoboda for valuable comments on the manuscript.

## Author contributions

Conceptualization: MG, PG, AP, DO, FN, IY, CB, JK, SM, CR, SG, BH, JP, SM, AA, CK, SRO

Data curation: MG, PG, AP, DO, FN, IY, AA, MB, SC, SD, SdV, DK, JK, JL, PL, NM, CBM, NO, JP, NP, CR, SS, DW, YB, NG, BH, IK, SK, KM, CN, MO, PR, KS, JS, TKN, XW, WW

Formal analysis: MG, PG, AP, DO, FN, IY, SD, DK, JK, JL, PL, SM, NO, JP, NP, CR, SS, DW

Funding acquisition: JP, CK

Investigation: SC, NO, CR, SS, RA, CB, NC, AC, MD, MD, CG, NH, TJ, IK, SK, SL, EL, KM, CN, KN, PR, KR, JPS, JS, TKN, XW

Methodology: MG, PG, AP, DO, FN, IY, CB, MB, SC, SdV, JL, PL, SM, NO, JP, CR, SS, DW, DA, RH, LK, QL, AL, FL, DS, BS, SM, AA, SRO

Project administration: MG, PG, SC, LC, FD, JL, SN, AW, KR, DS, WW, SRO

Resources: DF, RH, XJ, AL, LN, GKO, MO, CF, WW, HZ, JP

Software: MG, PG, AP, DO, FN, IY, AA, MB, SD, SdV, DK, JK, JL, PL, NM, CBM, NO, JP, DW, NC, DF, NG, RH, XJ, LK, A, FL, CM, LN, GKO, MO, KS, MMW, CF, WW

Supervision: MG, PG, SC, JL, NO, AW, CB, TD, DF, LN, KR, DS, MMW, CF, WW, JP, SM, AA, CK, SRO

Validation: MG, PG, AP, DO, FN, IY, AA, MB, SC, SD, SdV, DK, JK, JL, PL, SM, NM, CBM, NO, JP, NP, CR, SS, DW, SG, NG, BH, LK, FL, IM, CM, MO, KS, CF, WW

Visualization: MG, PG, AP, DO, FN, IY, CR, SRO

Writing – original draft: MG, PG, AP, DO, FN, IY, CK, SRO

Writing – review & editing: MG, PG, AP, DO, FN, IY, SdV, PL, SG, BH, XJ, SM, AA, CK, SRO

## Data and materials availability

All data generated and analyzed as part of this study are available for download in Neurodata Without Borders (NWB) format using the AllenSDK python toolkit: https://allensdk.readthedocs.io/en/latest/visual_behavior_optical_physiology.html and the DANDI archive: https://doi.org/10.48324/dandi.000711/0.231121.1730

## Code availability

Custom code written to support data processing and analysis in the study are available at: https://github.com/AllenInstitute/AllenSDK https://github.com/AllenInstitute/ophys_etl_pipelines https://github.com/AllenInstitute/ophys_nway_matching https://github.com/AllenInstitute/visual_behavior_analysis https://github.com/AllenInstitute/visual_behavior_glm https://github.com/AllenInstitute/brain_observatory_utilities

## Supplementary Figures

**Figure S1.**
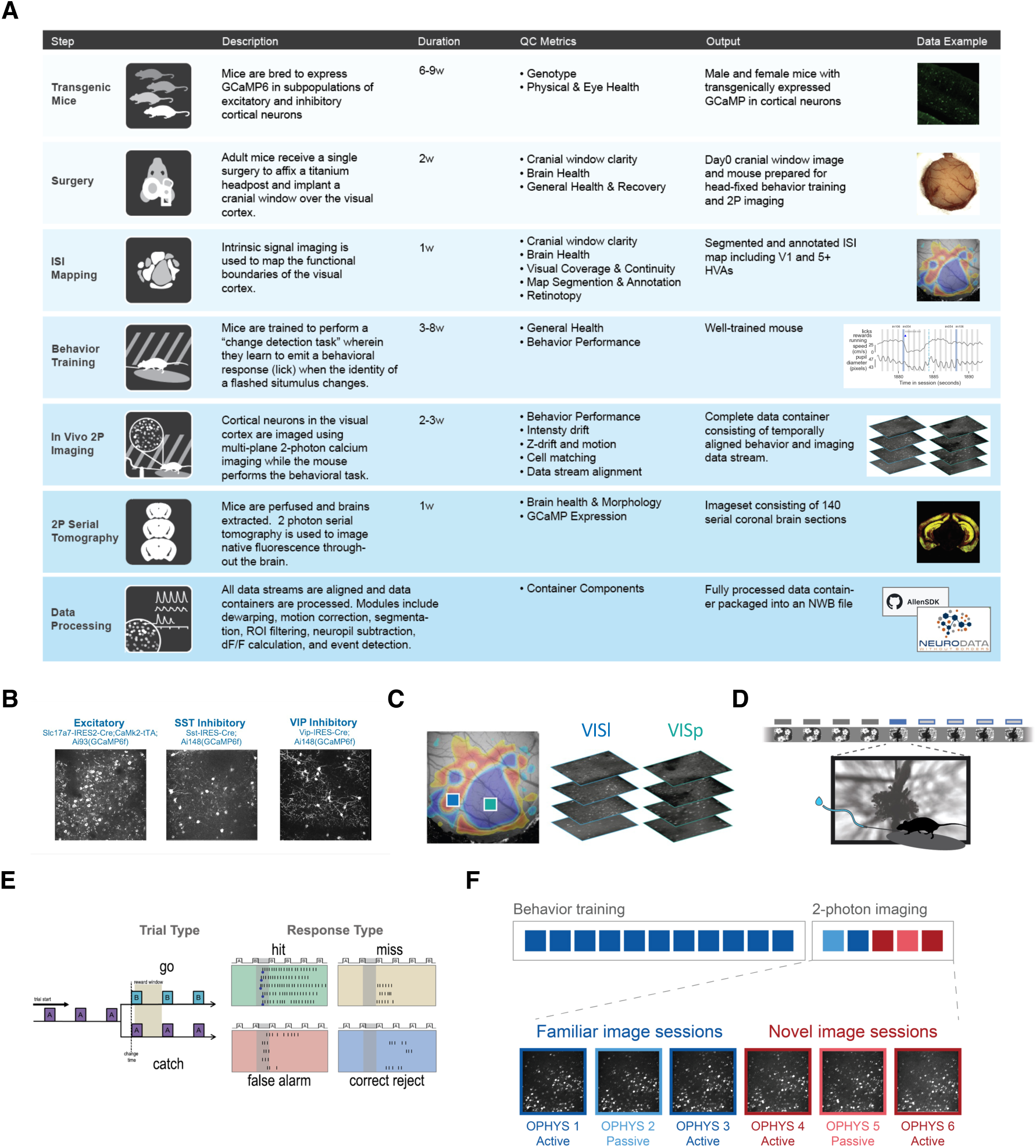
Allen Brain Observatory 2P Pipeline. (**A**) Experimental workflow, timeline, and Quality Control (QC) procedures. (**B**) Transgenic lines expressing GCaMP6 in genetically defined cell types. Images shown are maximum intensity projections of calcium fluorescence movies measured with 2-photon microscopy for each transgenic line. (**C**) Intrinsic signal imaging was performed to map retinotopic coordinates and area boundaries of mouse visual cortical areas. Recordings were targeted to the center of visual space within each area. Imaging depths spanned 75 – 375 um form the cortical surface, and up to 8 imaging planes were recorded in each session. (**D**) Schematic of go/no-go image change detection task. (**E**) Trial outcomes for go (change) and catch (non-change) trials based on mouse licking behavior. (**F**) Session types during 2-photon imaging consisted of either active behavior or passive viewing sessions, with familiar or novel stimuli. 2-photon imaging was performed in well-trained mice.

**Figure S2.**
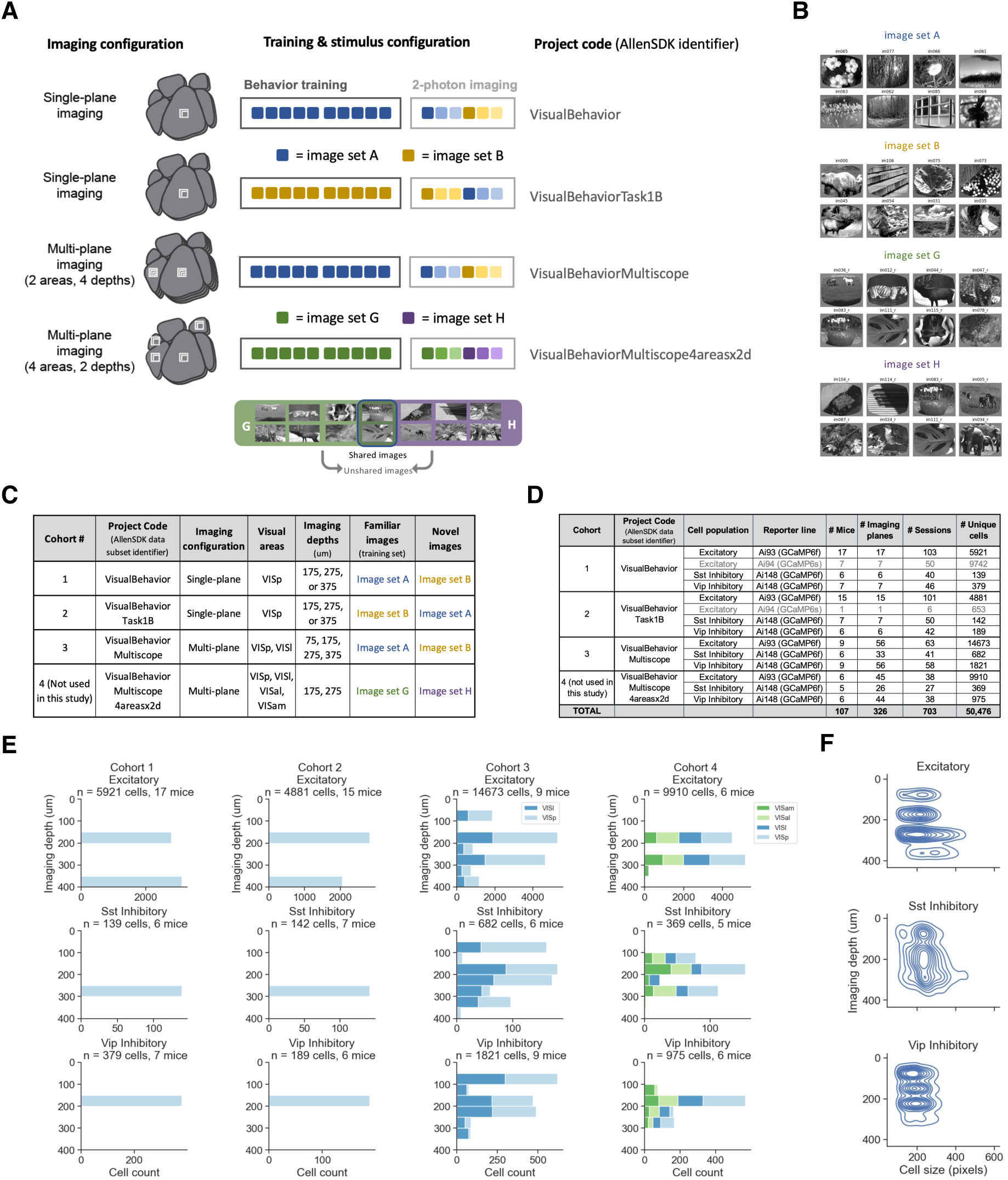
Allen Brain Observatory Visual Behavior 2P Dataset. (**A**) The Visual Behavior 2P dataset includes 4 cohorts of mice with different imaging configurations, areas and depths recorded, and image sets used during training. Some mice were trained on image set A and tested on image set B, while others had the order of image sets reversed. One group of mice had a unique design with 2 familiar images present in the novel image sessions; this dataset was not included in the current study but is available at brain-map.org. (**B**) Each image set consisted of 8 natural scene images. (**C**) Table describing the experimental conditions for the 4 cohorts of mice shown in A. (**D**) Numbers of mice, sessions, and cells recorded for each transgenic line, in each cohort. (**E**) Distribution of recorded cells by imaging depth for each cohort. Cohorts 1 and 2 used single-plane imaging and only one depth was targeted per mouse. Cohorts 3 and 4 used multi-plane imaging. Cells were targeted within each transgenic line based on known expression patterns of each cell population. (**F**) Density plot showing size of segmented cells across depths for each cell population.

**Figure S3.**
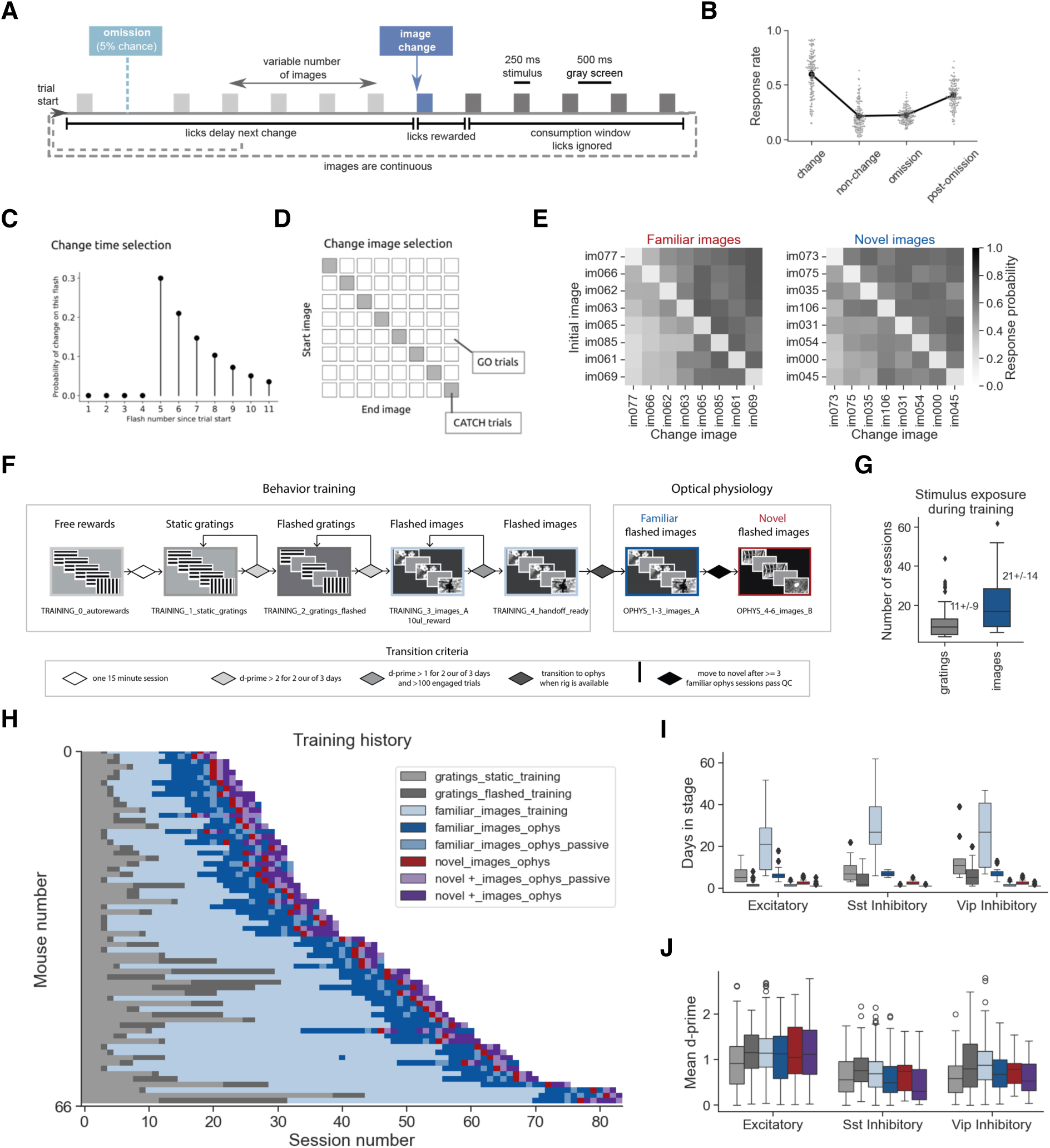
Go/no-go visual change detection task and training procedure. (**A**) Schematic of image change detection task, illustrating the stimulus cadence and dependence of task flow on mouse behavior. Stimuli are presented for 250 ms with a 500ms gray screen interval. Images are repeated a variable number of times (see panel C) before an image change occurs. Mice must lick within 750 ms after the image change to earn a water reward. Licks in a 3 second grace period after change onset are ignored. Licks outside the reward window and grace period cause the trial to reset, delaying the onset of the next change. During 2-photon imaging sessions in well trained mice, 5% of non-change image presentations are randomly omitted. Images are never omitted during training. (**B**) Response rate for changes, non-changes, omissions, and post-omission images in well trained mice, in sessions during which 2-photon imaging was performed. (**C**) Change time distribution, illustrating the range of image presentations (flashes, x-axis) over which a change time can be drawn after the start of a trial, with y-axis showing probability of image change occurring on each image presentation. (**D**) Transition probability matrix illustrating 64 possible transitions between each of the 8 images shown in each session. Transitions from a given image to itself are catch trials, shown on the diagonal. (**E**) Response probability across image transitions for Familiar and Novel image sessions in well trained mice. (**F**) Schematic of behavioral training procedure. Mice are automatically progressed through a series of training stages, beginning with static gratings and ending with flashed natural scene images, before being transitioned to 2-photon imaging. During the 2-photon imaging stage of the experiment, mice perform the task with either familiar or novel images. In some sessions, mice passively view the same stimuli in open loop mode with the lick spout retracted after being given their daily water. (**G**) Number of sessions with image stimuli and gratings during behavioral training. (**H**) Training history for all mice included in the study. Colors indicate training stage, as in panel F, with the addition of repeated sessions with novel images in purple, and passive viewing sessions indicated in a unique shade of blue or purple as shown in legend**. (I)** Number of sessions spent in each behavior stage across transgenic mouse lines (legend shared with panel H). (**J**) Behavioral performance, quantified as the mean d-prime value over the course of a session (including both engaged and disengaged periods) across training stages for each transgenic mouse line (legend shared with panel H).

**Figure S4.**
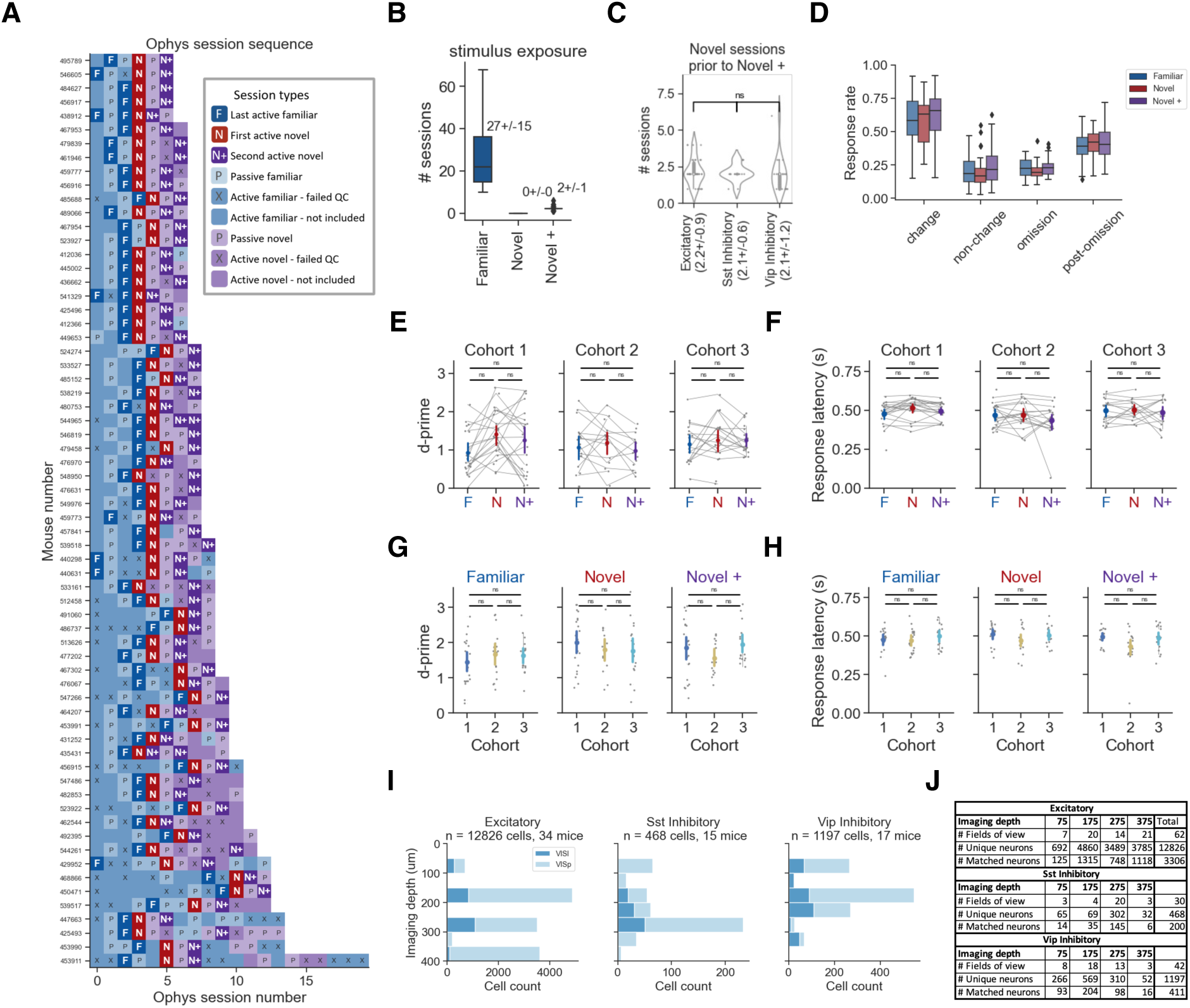
Selection criteria and task performance for Familiar, Novel, and Novel+ sessions. (**A**) Session sequence during 2-photon imaging for mice included in this study, highlighting sessions that were selected for analysis (darker colors, see legend). Selected sessions include the last active behavior session with familiar images (Familiar, dark blue), the first session with novel images (Novel, red), and a subsequent active behavior session with novel images (Novel+, dark purple). Passive viewing sessions (indicated with letter P) were interleaved among active behavior sessions but were not used in the primary analysis for this study (but see fig. S9C). 2-photon imaging sessions that did not pass quality control (QC) are indicated with an X. (**B**) Number of sessions with familiar or novel images prior to the 2-photon imaging sessions included in this study for each session type (Familiar, Novel, Novel+). (**C**) Number of sessions with novel images experienced prior to the Novel+ session for mice expressing GCaMP6 in different cell types. (**D**) Response rate for relevant task conditions (image changes, repeated non-changes, image omissions, and post-omission image presentations) separated by experience level (Familiar, Novel, Novel+). Response rate is measured as the fraction of trials for each condition where the mouse emitted a lick in a 750ms window after the event. (**E**) Behavior performance, measured as the mean discriminability index, d-prime, across sessions for each experience level (Familiar, Novel, Novel+) for each cohort of mice (see fig. S2A-D for description of cohorts). Gray points and connecting lines represent individual sessions from each mouse, colored points represent mean +/- 95% confidence intervals across sessions. No significant differences were found across experience levels in any of the cohorts (p = 0.05, one-way ANOVA followed by Tukey HSD). (**F**) Behavioral response latency following image changes across experience levels for each cohort. No significant differences found across experience levels (statistics performed as described for panel E). (**G**) Behavior performance, measured as the maximum d-prime for each session, across cohorts of mice (colored points) for each experience level (columns) (see fig. S2A-D for description of cohorts). Colored points represent mean +/- 95% confidence interval across mice. No significant differences were found across cohorts (p = 0.05, one-way ANOVA followed by Tukey HSD). (**H**) Behavioral response latency following image changes across cohorts (colored points) for each experience level (columns). Statistics were performed as in panel G. (**I**) Number of cells recorded across imaging depths (y-axis) and areas (colors) for the set of mice and sessions included in this study, based on the criteria described in panel A. (**J**) Numbers of imaging planes, unique neurons, and matched neurons per cell type and imaging depth (in um).

**Figure S5.**
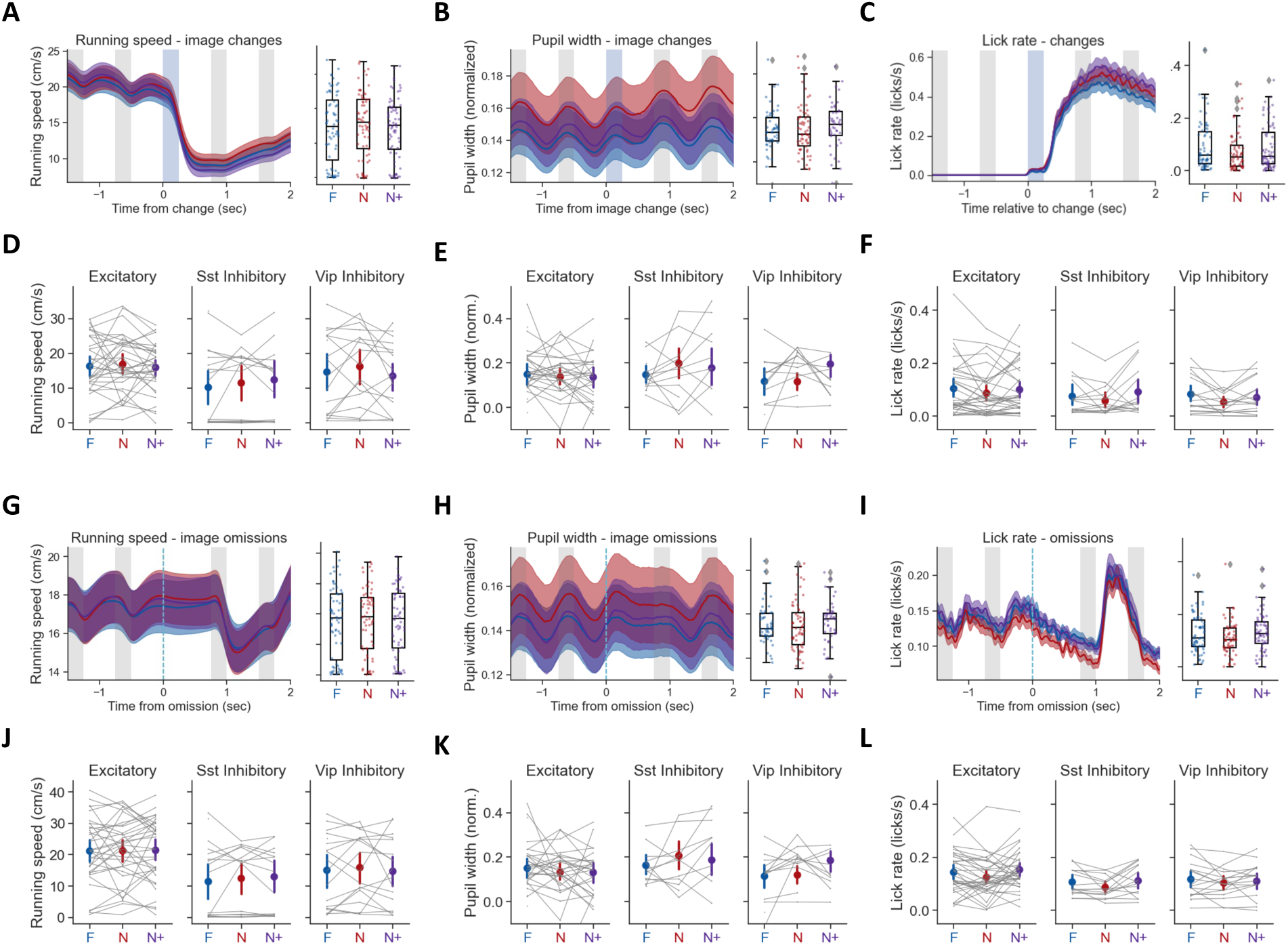
Average pupil, running and licking across experience levels. (**A**) Image change aligned running speed across experience levels (Familiar, Novel, Novel +). Left panel, time course of image change locked running speed averaged across sessions. Right panel, quantification of mean running speed across sessions for each experience level. Running speed was averaged over a 500ms window following image changes for each trial, then averaged across all trials within each session. Each data point is one session. (**B**) Mean image change aligned pupil width averaged across sessions for each experience level. Pupil width during behavior performance is normalized to the average pupil width during the 5-minute gray screen period prior to the start of the task to produce a fractional value relative to the no stimulus period. Left panel, time course normalized pupil width. Right panel, quantification of pupil width across sessions for each experience level. Normalized pupil width was averaged over a 500ms window following image changes for each trial, then averaged across all trials within each session. Each data point is one session. (**C**) Mean image change aligned lick rate averaged across sessions for each experience level. (**D**) Average running speed during image changes across sessions (gray points) and experience levels (colored points) split by transgenic mouse line (expressing GCaMP6f in excitatory, Sst, of Vip populations). Gray lines connect data points for each mouse. (**E**) Average normalized pupil width during image changes across sessions (gray points) and experience levels (colored points) split by transgenic mouse line (excitatory, Sst, Vip). Gray lines connect data points for each mouse. (**F**) Average lick rate during image changes across sessions (gray points) and experience levels (colored points) split by transgenic mouse line (excitatory, Sst, Vip). Gray lines connect data points for each mouse. (**G-I**) Same as panels A-C but aligned to image omissions instead of image changes. Behavior was quantified in a 750ms window following each image omission in a given session, then averaged across the session. (**J-L**) Same as panels D-F but for image omissions. Behavior was quantified in a 750ms window following each image omission in a given session, then averaged across the session, and plotted across experience levels for each transgenic line. For all panels, significance of differences across experience levels was computed by a one-way ANOVA followed by Tukey HSD (p=0.05). No significant differences were detected. All error bars indicate +/- 95% confidence intervals.

**Figure S6.**
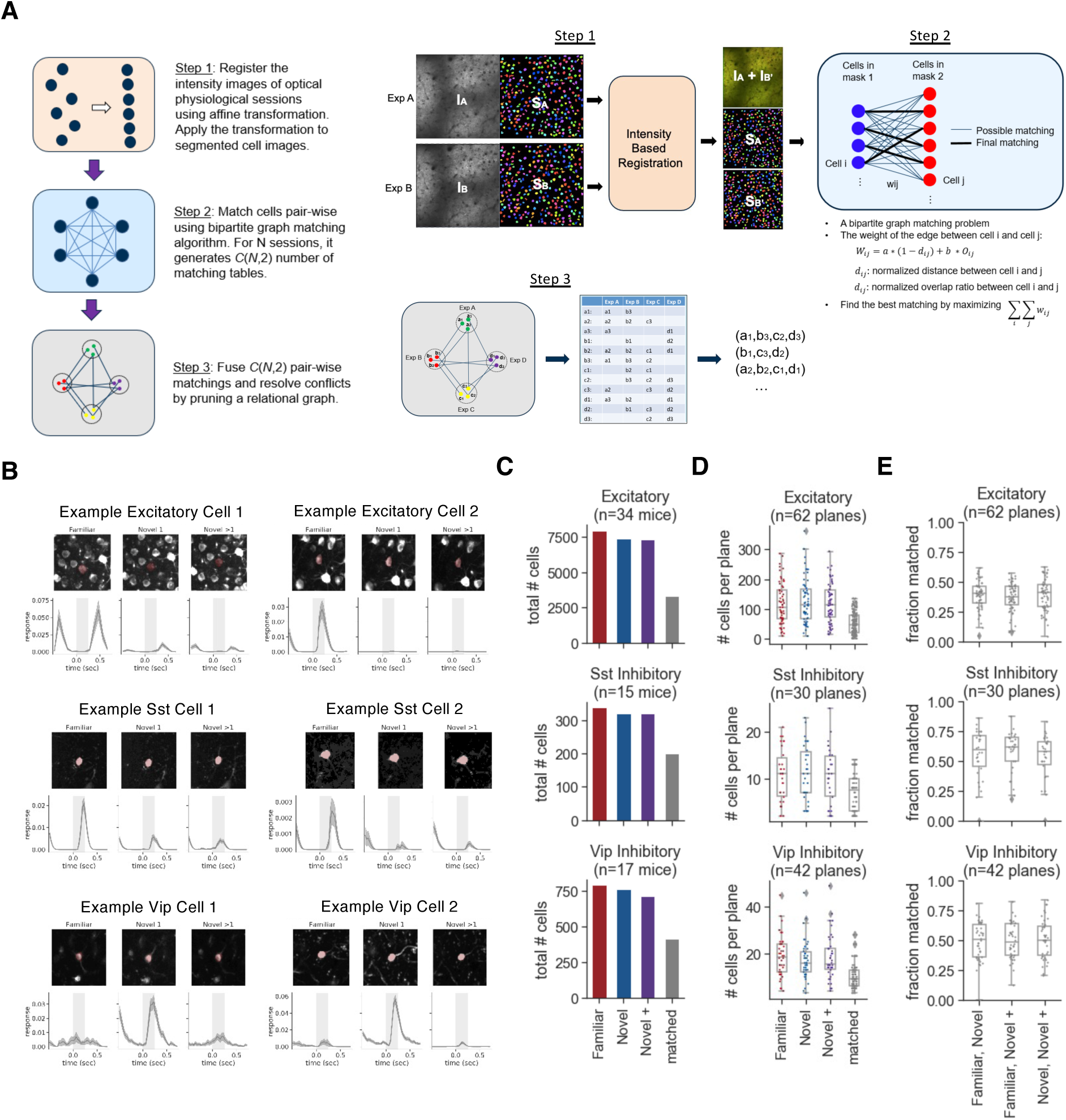
Description of cross-session registration and cell matching. (**A**) Schematic of cell matching procedure. Average projection images from each pair of 2-photon imaging sessions are aligned to each other and the registration transform is applied to the segmented ROI masks (step 1). A bi-partite graph matching algorithm is used to identify the best ROI match between the two sessions using intersection over union (IOU) of the ROI masks, and the inter-centroid distance with a maximum allowable distance of 10 pixels (step 2). Each pair of sessions was registered and matched, and cell specimen IDs were unified across all pairwise matches (step 3). (**B**) Example images of matched cells and average image responses in each of the 3 session types included in this study. Two example cells (columns) are shown for each cell class (rows: Excitatory, Sst, Vip). (**C**) Total number of cells detected in each session (colors, Familiar, Novel, Novel +) and number matched across all 3 sessions (gray). (**D**) Number of cells detected in each session and matched across all sessions for each 2-photon field of view (each point is one imaging plane). (**E**) Fraction of cells matched for each pair of sessions for each mice expressing GCaMP6f in each cell type (excitatory, Sst inhibitory, Vip inhibitory). Individual points are unique imaging planes.

**Figure S7.**
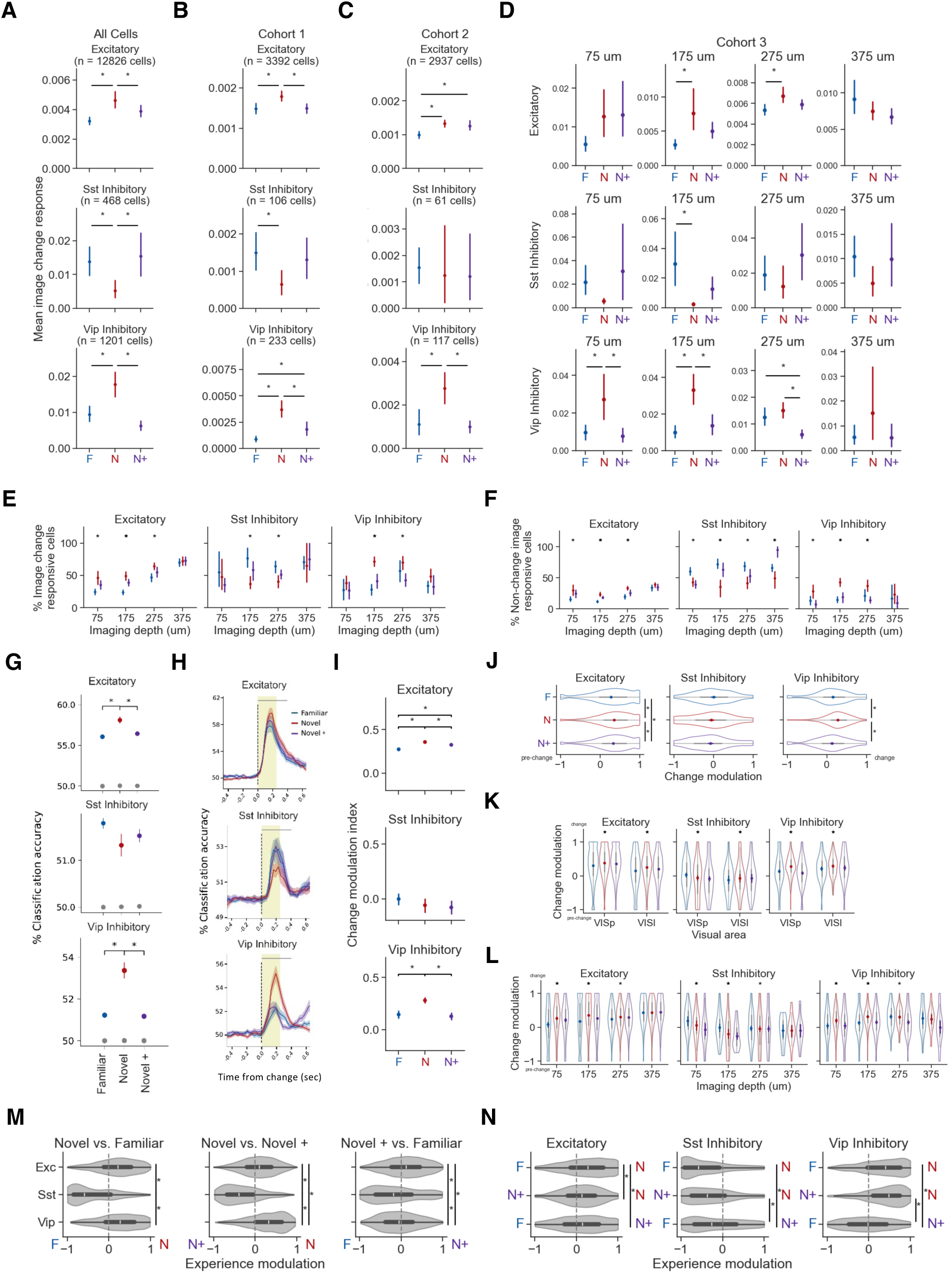
Quantification of image change response across experience levels. Statistics for all panels were performed by one-way ANOVA across experience levels followed by Tukey HSD (p=0.05). Data points represent mean +/- 95% confidence interval across neurons (except for panel G which averages across sessions). (**A-D**) Average image change evoked activity across experience levels (Familiar in blue, Novel in red, Novel+ in purple) for each cell type (top row is excitatory, middle row is Sst, bottom row is Vip) for all cells (panel A),or separately for groups of mice trained with different image sets (cohorts 1 and 2, panels B and C), or mice imaged with multi-plane 2-photon imaging spit by imaging depth (panel D). (**E**) Percent of cells with significant responses to image changes across experience levels for each imaging depth (including both single and multi-plane sessions – see figure S2 for description of cohorts & imaging configurations). Responsiveness is defined has having a significantly larger response during image changes compared to a shuffled distribution of activity from the 5-minute gray screen period prior to each session (see Fig. 1F for illustration of session structure) for at least 10% of all image changes. (**F**) Fraction of responsive cells for non-change image presentations. (**G**) Population decoding accuracy for classification of image changes from pre-change (i.e. repeated) image presentations across cell types (rows) and experience levels (colors). Colored points represent the mean across sessions +/- 95% confidence intervals for each experience level. Values for shuffled data are in gray. (**H**) Time-resolved decoding accuracy, reflecting the percentage of correct identifications over time for change vs. non-change conditions across experience levels (colors) and cell types (rows). Classification was performed separately on each time point and averaged in a 250ms window after stimulus onset to produce the values in (panel **G**). (**I**) Average value of change modulation index across experience and cell types, computed as each cell’s mean response to image changes (500 ms window after change onset) minus the mean response to pre-change images (500 ms window after onset of pre-change image), over the sum of the two. Positive values of the index indicate larger responses to image changes compared to non-change images. Negative values indicate larger responses to repeated pre-change images. (**J**) Distribution of change modulation index across cell types (columns) and experience levels (colors), demonstrating heterogeneity in response properties within each cell type. (**K**) Distribution of change modulation index across cell types (columns) and experience levels (colors), split by imaged visual area (on x-axis: VISp, primary visual cortex; VISl, lateromedial visual area). (**L**) Distribution of change modulation index split by imaging depth for each cell type. (**M**) Distribution of experience modulation index for each pair of experience levels (i.e. sessions), computed as each cell’s mean response to image changes in “session 1” minus the mean response to image changes in “session 2”, over the sum. For each panel, “session 1” is listed first in the title and “session 2” is listed second. Stronger responses during session 1 compared to session 2 are indicated by positive values on the x-axis. Rows are cell types (excitatory, Sst inhibitory, Vip inhibitory). Statistics are performed across cell types. Note that all cell types have relatively wide distributions, indicating that despite biases in the average, there are a diversity of response types. (**N**) Same data as panel M, showing strength of experience modulation for each pair of experience levels, grouped by cell type. Statistics are performed across experience level pairs within each cell type. Excitatory cells have stronger responses to image changes during sessions with less overall experience (i.e. Novel sessions compared to Familiar and Novel+, Novel+ session compared to Familiar session). Sst cells have stronger responses to sessions with more overall experience (Familiar sessions compared to Novel and Novel+, Novel+ compared to Novel). Vip cells have stronger responses to Novel image changes compared to Familiar and Novel+, however the response to Familiar and Novel+ image changes is the same on average, indicating that novelty effects saturate within just a few days across the Vip population.

**Figure S8.**
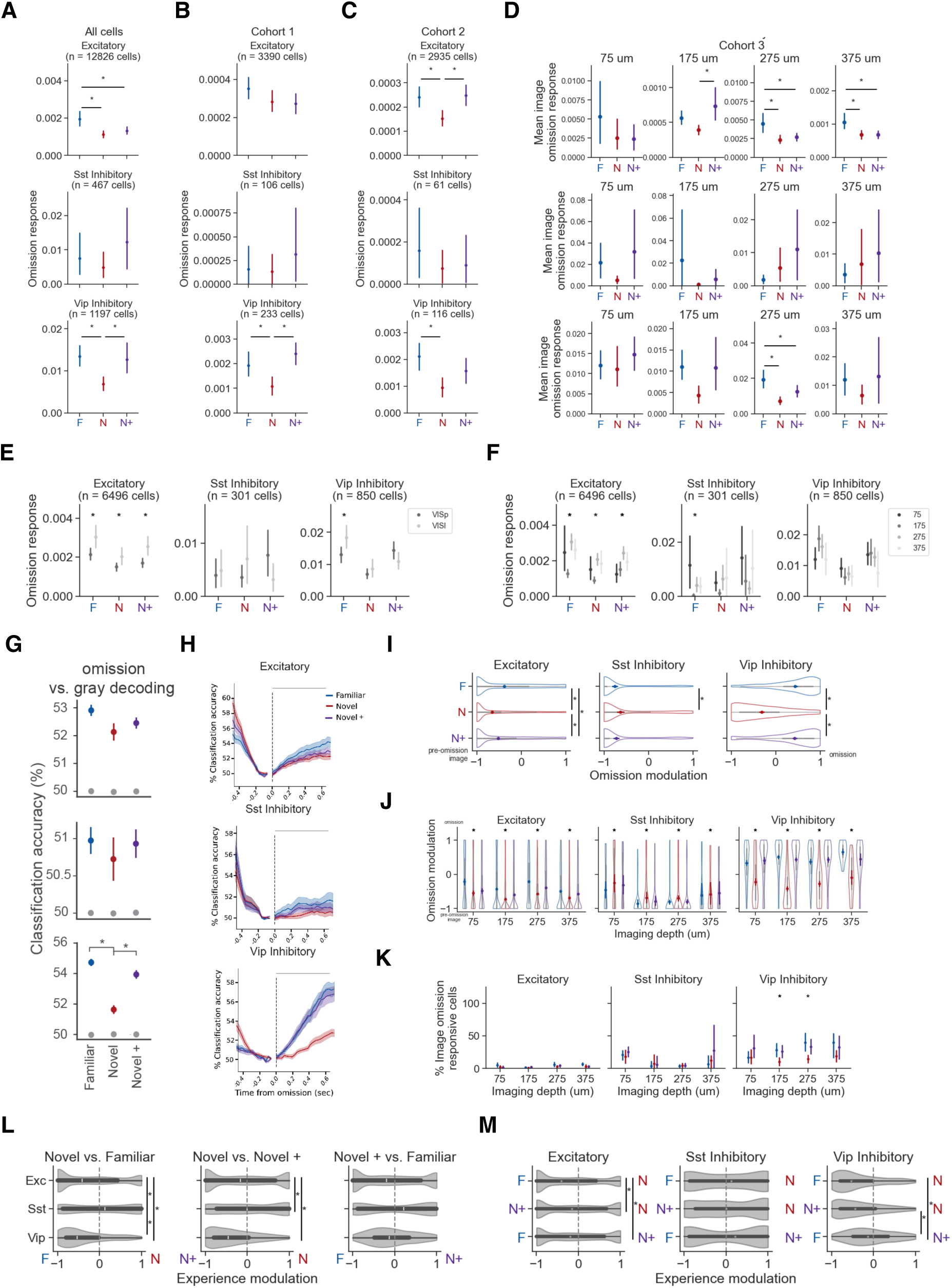
Quantification of image omission related activity across experience levels. Statistics for all panels were performed by one-way ANOVA across experience levels followed by Tukey HSD (p=0.05). Data points represent mean +/- 95% confidence interval across neurons (except for panel G which averages across sessions). (**A-D**) Average image omission evoked activity across experience levels (Familiar in blue, Novel in red, Novel+ in purple) for each cell type (top row is excitatory, middle row is Sst, bottom row is Vip) for all cells (panel A),or separately for groups of mice trained with different image sets (cohorts 1 and 2, panels B and C), or mice imaged with multi-plane 2-photon imaging spit by imaging depth (panel D). (**E**) Average omission response across experience levels and cell types split by visual area (VISp, primary visual cortex; VISl, lateromedial visual area), demonstrating stronger omission responses in higher area VISl compared to VISp for excitatory cells and Vip cells during Familiar sessions. Results are shown for multi-plane imaging sessions (cohort 3). (**F**) Average omission response across experience levels and cell types split by imaging depth for multi-plane imaging sessions. (**G**) Population decoding accuracy for classification of activity following unexpectedly omitted images relative to the gray screen period just prior to omission onset across cell types (top row is excitatory, middle row is Sst, bottom row is Vip). Classification was performed on each session then averaged across sessions. Colored points represent the mean across sessions +/- 95% confidence intervals for each experience level. Values for shuffled data are in gray. (**H**) Time-resolved decoding accuracy, reflecting the percentage of correct identifications over time for image omission vs. pre-omission gray screen across experience levels (colors) and cell types (rows). Classification was performed separately on each time point and averaged in a 750ms window after stimulus onset to produce the values in (panel G). (**I**) Distribution of omission modulation index across cell types and experience levels, computed as each cell’s mean response during image omissions (750 ms window after omission onset) minus the mean response during the gray screen period prior to omissions (500ms window prior to omission onset), over the sum of the two. Positive values of the index indicate stronger activity during omissions compared to expected gray screen periods. Negative values indicate larger responses during the gray screen preceding the omission compared to after the image was omitted. (**J**) Distribution of omission modulation index across cell types and experience levels, split by imaging depth (x-axis). (**K**) Percent of cells with significantly larger responses during image omissions (750ms window after omission) compared to a shuffled distribution of activity from the 5-minute gray screen period at the start of each imaging session (see Fig. 1F for illustration of session structure). Responsiveness is defined as having at least 10% of image omissions with a significant response (p<0.05 via paired t-test over 100 iterations of shuffle for gray screen period). (**L**) Distribution of experience modulation index for each pair of experience levels (i.e. sessions), computed as each cell’s mean response during omissions in “session 1” minus the mean response during omissions in “session 2”, over the sum. For each panel, “session 1” is listed first in the title and “session 2” is listed second. Stronger responses during session 1 compared to session 2 are indicated by positive values on the x-axis. Rows are cell types (excitatory, Sst inhibitory, Vip inhibitory). Statistics are performed across cell types. (**M**) Same data as panel L, showing strength of experience modulation for each pair of experience levels, grouped by cell type. Statistics are performed across experience level pairs within each cell type. Vip cells have larger omission responses in Familiar and Novel+ sessions compared to Novel sessions. Excitatory and Sst cells’ omission related activity is not experience dependent on average, although a small subset of neurons at the extremes show a bias of omission activity for a given session type, with no dependence on experience level.

**Figure S9.**
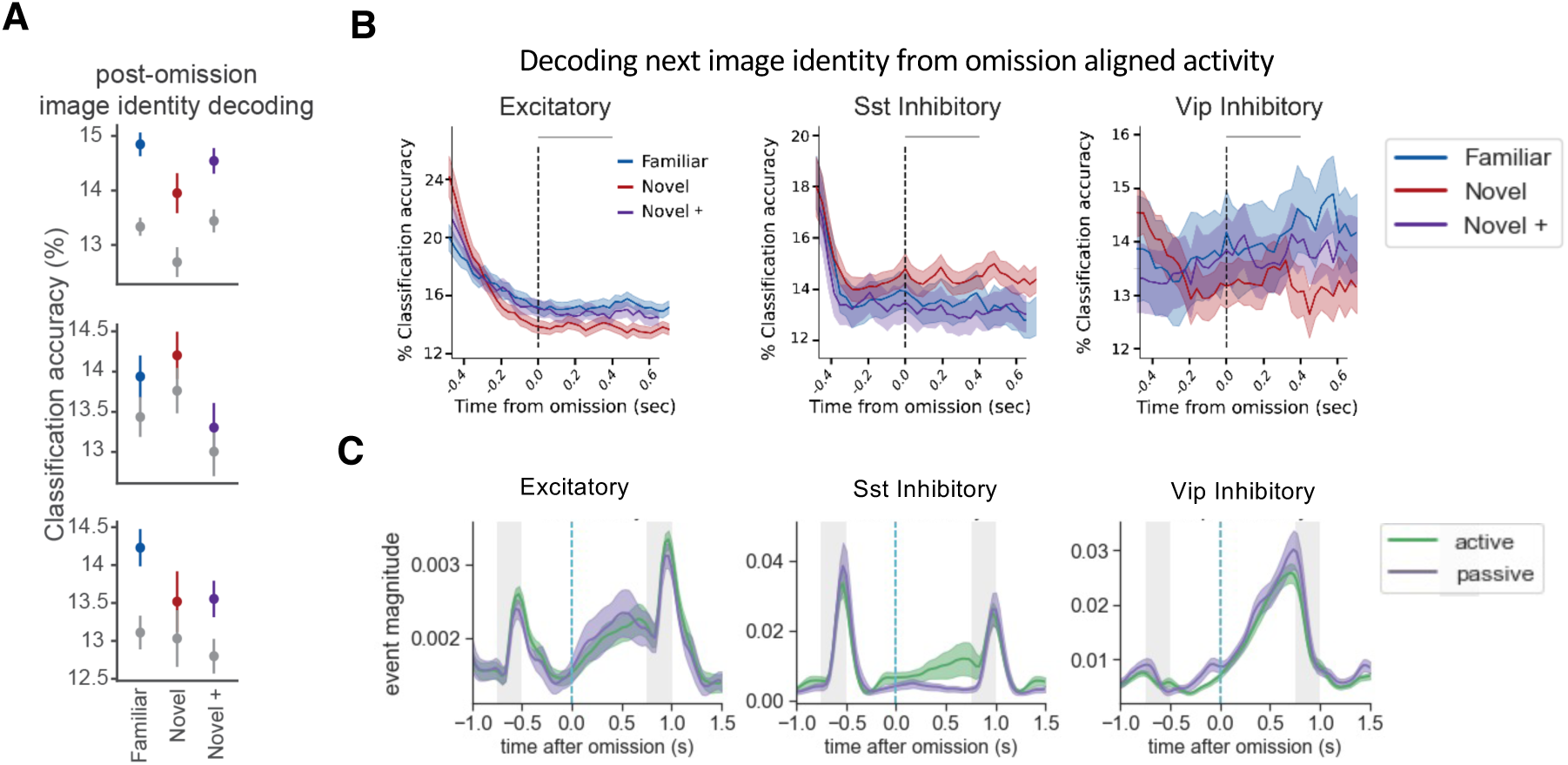
Omission related activity does not depend on next image identity or behavior state. (**A**) Decoding of post-omission image identity from population activity during omissions for excitatory (top row), Sst (middle row), and Vip (bottom row) sessions. Data points compare the classifier’s ability to decode the identity of omitted image. Classification was performed on each timepoint, as in panel B, then averaged over the 750ms window following omission onset for each session, then averaged across sessions to produce the colored data points in panel A. Gray points represent shuffle. Data points represent mean +/- 95% confidence interval. No significant differences were detected across sessions (one way ANOVA followed by Tukey HSD, p=0.05). (**B**) Time resolved decoding of the next image’s identity based on activity aligned with omissions. (**C**) Omission-aligned population responses (+/- SEM) across cell types, comparing sessions during active behavior performance (green), and passive viewing sessions (purple) in which the mouse was sated and viewed the stimulus in open loop, demonstrating that task engagement does not impact population level omission responses of Vip or excitatory neurons,

**Figure S10.**
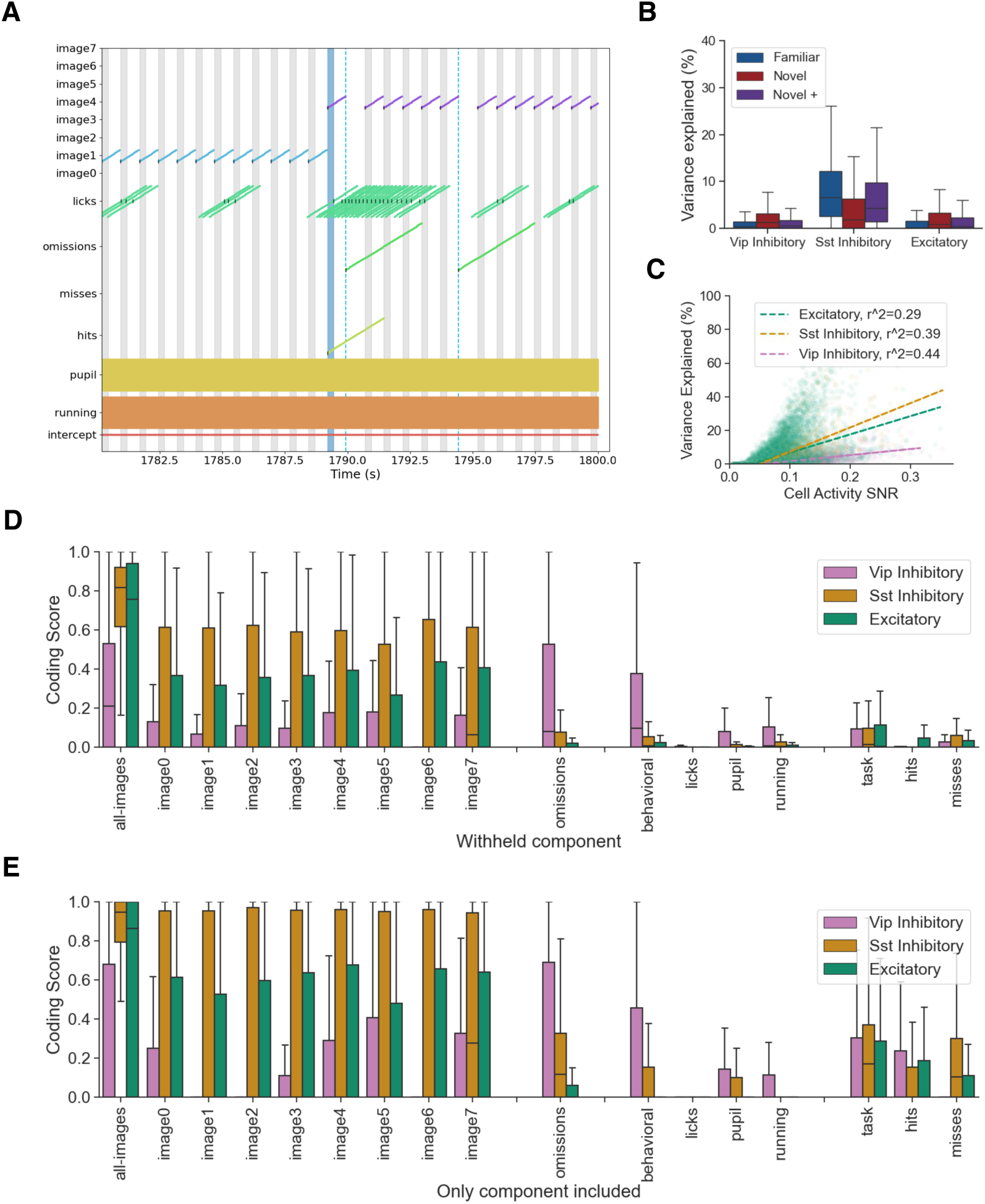
Encoding model methods and validation. (**A**) Illustration of kernel length and timing for each feature. Kernels for each of the 8 images in each image set start at image onset (gray bands mark image presentations) and extend for 750ms. Kernels for licking are aligned to each detected lick (black tick marks) and extend forward and backward in time 1s. Kernels for omissions are aligned to the time point when each image would have been presented (dashed blue line) and extend for 3s. Kernels for hits and misses are aligned to the onset of the change image (blue band marks change image presentation) and extend for 2.25s. Kernels for running speed and pupil diameter are continuously convolved with the data timeseries and extend forward and backward in time 1s. (**B**) Distribution of explained variance for each cell type split by experience level. Boxplots show the quartiles of each cell type and the whiskers mark +/- 1.5x Interquartile Range (Q3-Q1). Outliers are not shown for clarity. (**C**) Explained variance of the full model for each cell against the signal-to-noise ratio (SNR) of the neural activity trace. Here, SNR is the mean of each cell’s dF/F signal divided by its standard deviation, computed over the full 1-hour session. Lines show the best linear fit, and correlation values are shown in inset. (**D**) Distribution of coding scores for each model feature, split by cell type. Boxplots show the quartiles of each cell type and the whiskers mark +/- 1.5x Interquartile Range (Q3-Q1). Outliers are not shown for clarity. (**E**) Distribution of coding scores for each cell type computed by refitting reduced models with only a single feature.

**Figure S11.**
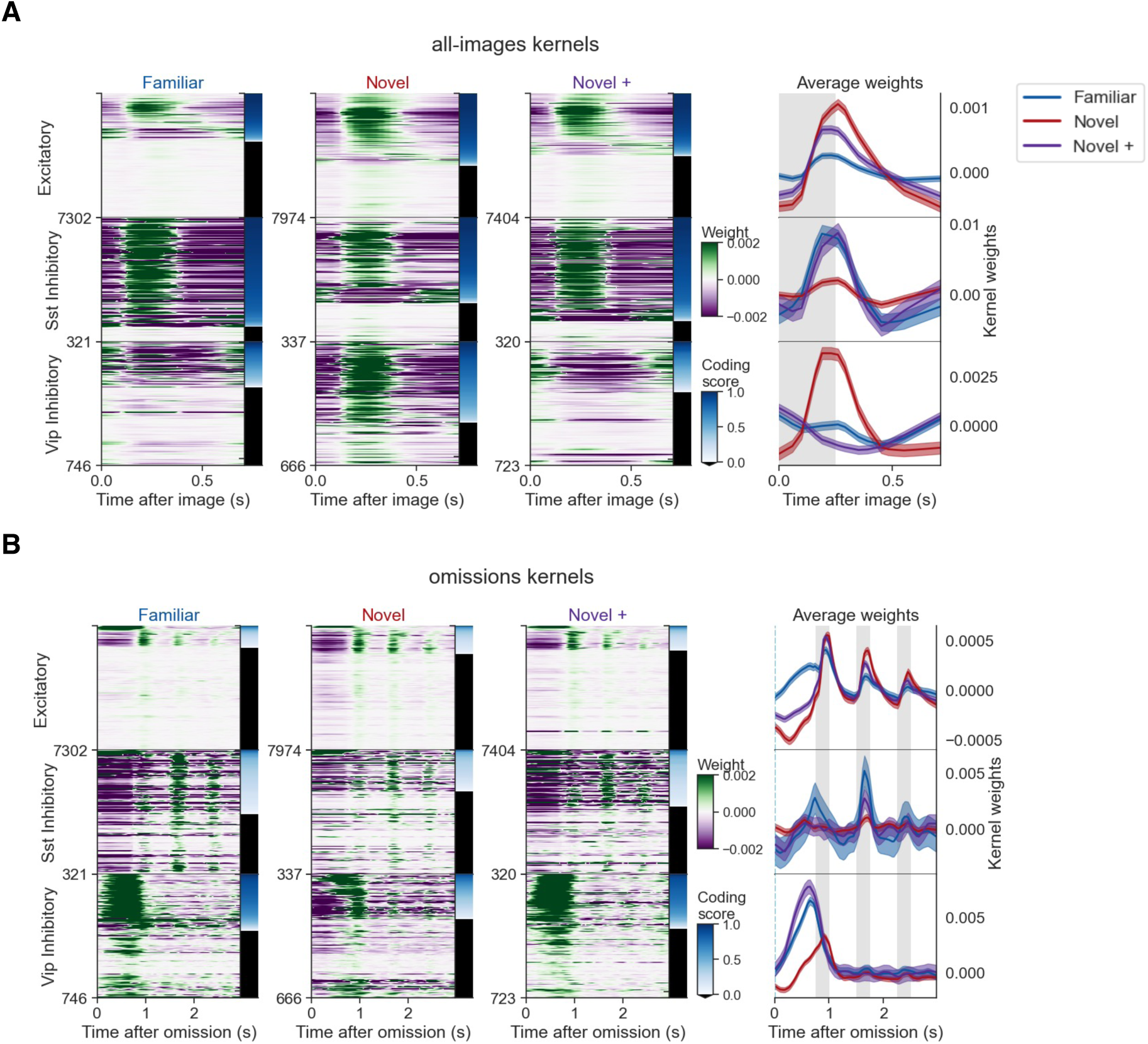
Image and omission kernels for all cells. (**A**) Kernel weights for image presentations experience levels (Familiar, Novel, Novel+). Left three columns show heatmaps of learned image kernels for all cells. Cells are sorted by cell type, and then by image coding score within each cell type. Rows are not matched across experience levels. Right column shows average image kernel weights for each cell type (rows) colored by experience level. (**B**) Omission kernels across experience levels. Left three columns show heatmaps of learned omission kernels for all cells. Cells are sorted by cell type, and then by omission coding score. Right column shows average omission kernel weights for each cell type, colored by experience level.

**Figure S12.**
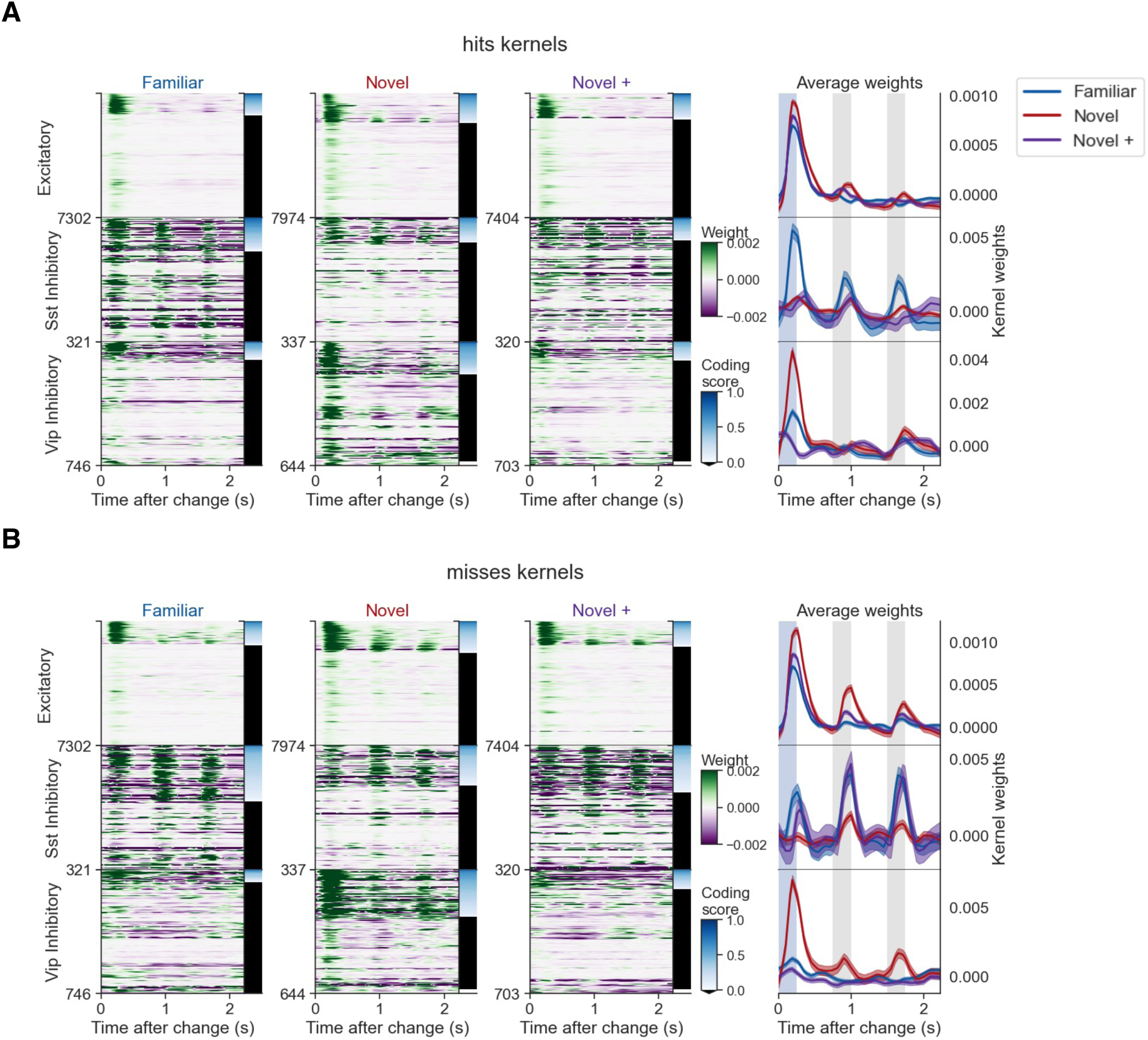
Task kernels for all cells. (**A**) Kernel weights for hits across experience levels (Familiar, Novel, Novel+). Hits are image changes with a correct licking response, earning a water reward. Left three columns show heatmaps of learned hit kernels for all cells. Cells are sorted by cell type, and then by strength of coding score within each cell type. Rows are not matched across experience levels. Right column shows average hit kernel weights for each cell type (rows) colored by experience level. (**B**) Kernel weights for misses across experience levels. Misses are image changes where the mouse did not lick and thus no reward was delivered. Left three columns show heatmaps of learned miss kernels for all cells. Cells are sorted by cell type, and then by strength of miss coding score. Right column shows average miss kernel weights for each cell type, colored by experience level.

**Figure S13.**
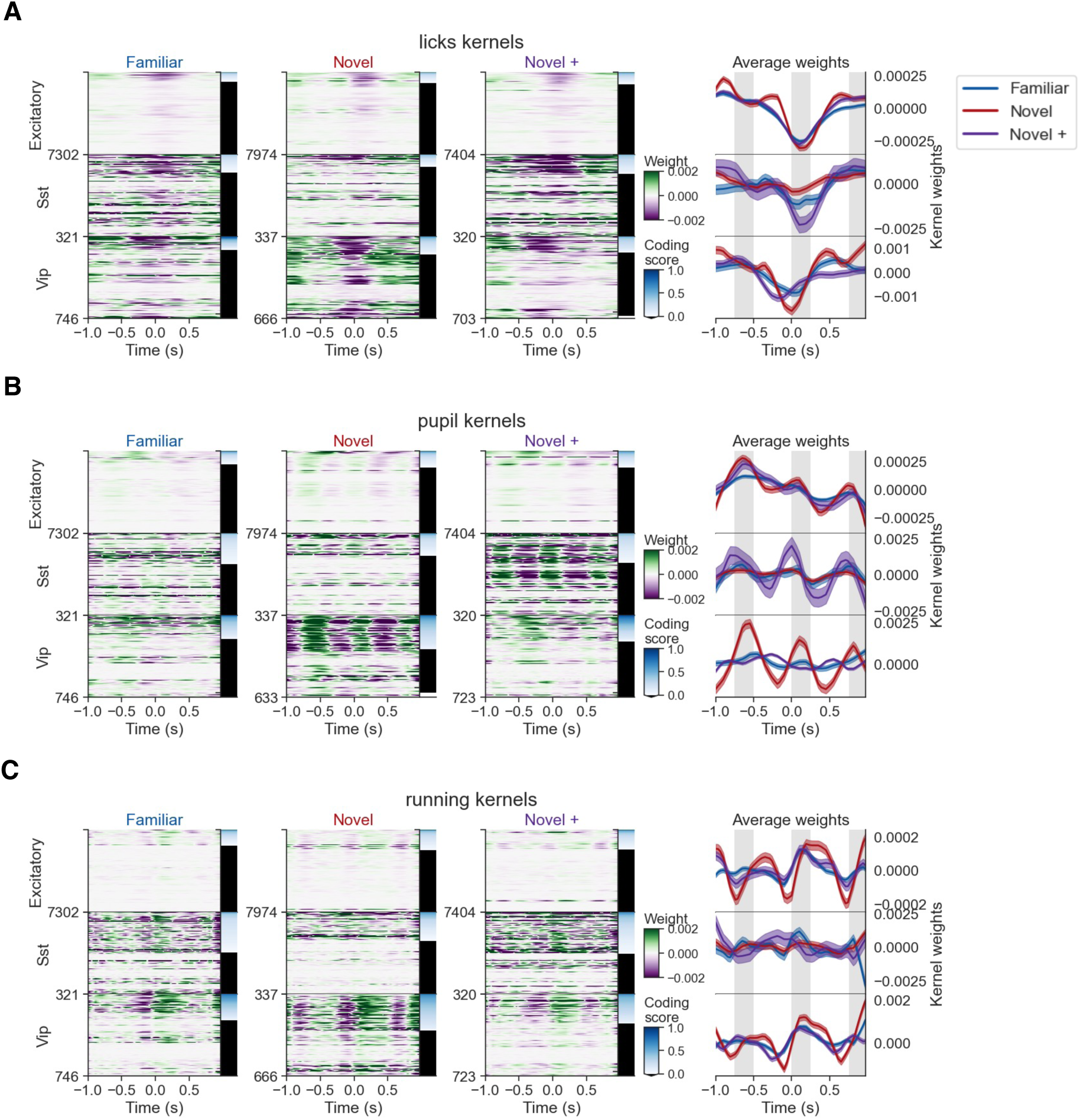
Behavior kernels for all cells. (**A**) Kernel weights for licks across experience levels (Familiar, Novel, Novel+). Lick kernels spanned +/-1 second around each lick onset. Left three columns show heatmaps of learned hit kernels for all cells. Cells are sorted by cell type, and then by strength of lick coding score within each cell type. Rows are not matched across experience levels. Right column shows average lick kernel weights for each cell type (rows) colored by experience level. (**B**) Kernel weights for pupil diameter across experience levels. Pupil kernels were continuous across the session, taking a +/- 1 second window around each time point. Left three columns show heatmaps of learned pupil kernels for all cells. Cells are sorted by cell type, and then by strength of pupil coding score. Right column shows average pupil kernel weights for each cell type, colored by experience level. (**C**) Kernel weights for running speed across experience levels. Running kernels were continuous across the session, taking a +/- 1 second window around each time point. Left three columns show heatmaps of learned running kernels for all cells. Cells are sorted by cell type, and then by strength of coding score. Right column shows average running kernel weights for each cell type, colored by experience level.

**Figure S14.**
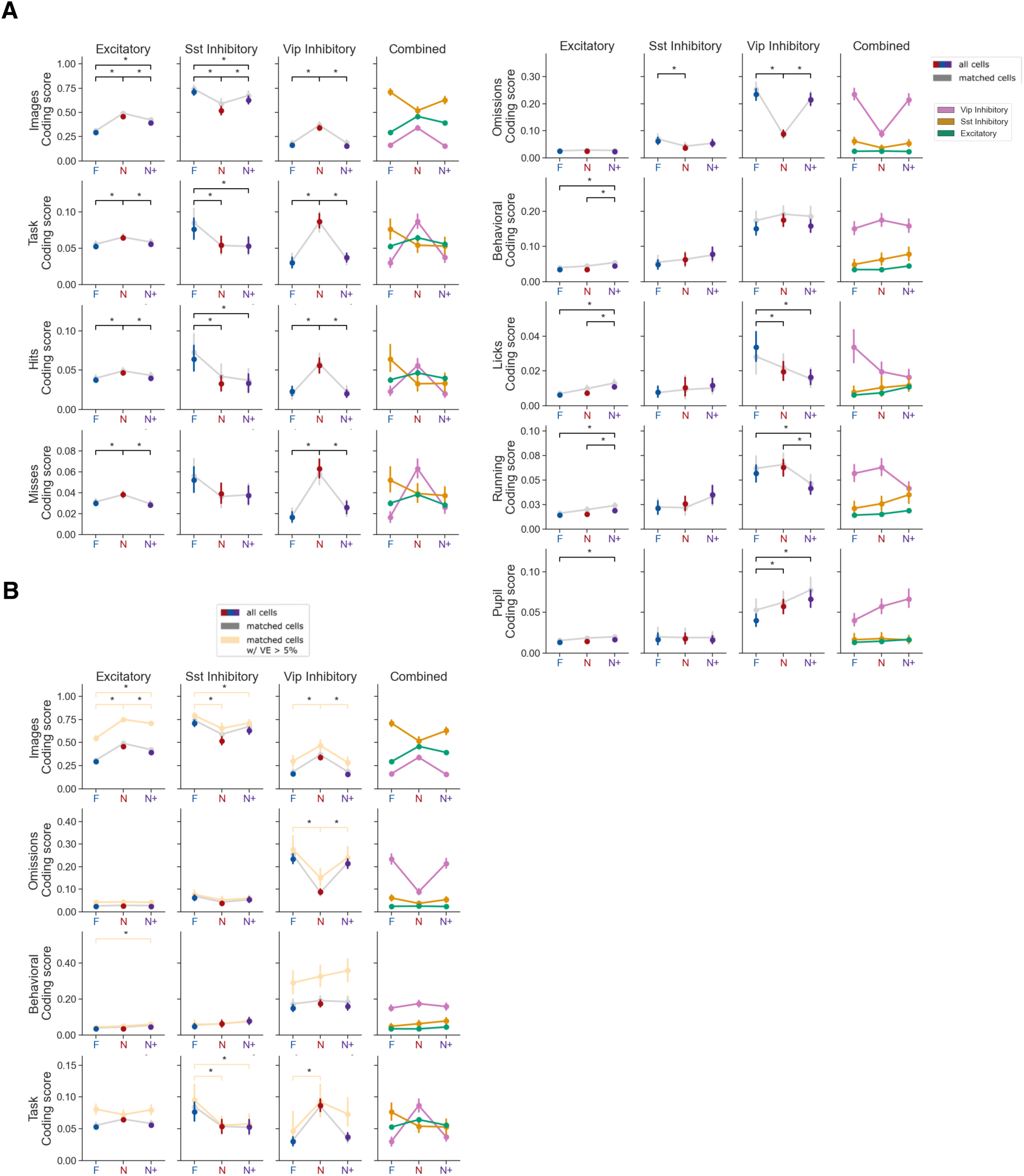
Coding scores across experience levels and cell types. (**A**) Average coding scores for each experience level (colored points) across cell types (left columns) for main feature groups (images, omissions, task, behavioral) and individual features within groups (hits, misses, licks, running, pupil). Colored points show mean +/- 95% confidence intervals. Gray points connected by gray lines show values for cells matched across all experience levels. Differences between experience levels are marked as significant after a one-way ANOVA followed by Tukey HSD (p = 0.05). Statistics were computed on all cells. Right column shows direct comparison across cell types (mean of all cells within each cell type). (**B**) Average coding scores across experience levels (colored points) and cell types (left columns) for main feature groups (images, omissions, task, behavior) for all cells (red, blue, purple points), matched cells (gray lines and points), and matched cells with model explained variance of at least 5% in at least one session (light yellow points). Statistics were performed across matched cells passing the 5% explained variance threshold. Right column shows direct comparison across cell types.

**Figure S15.**
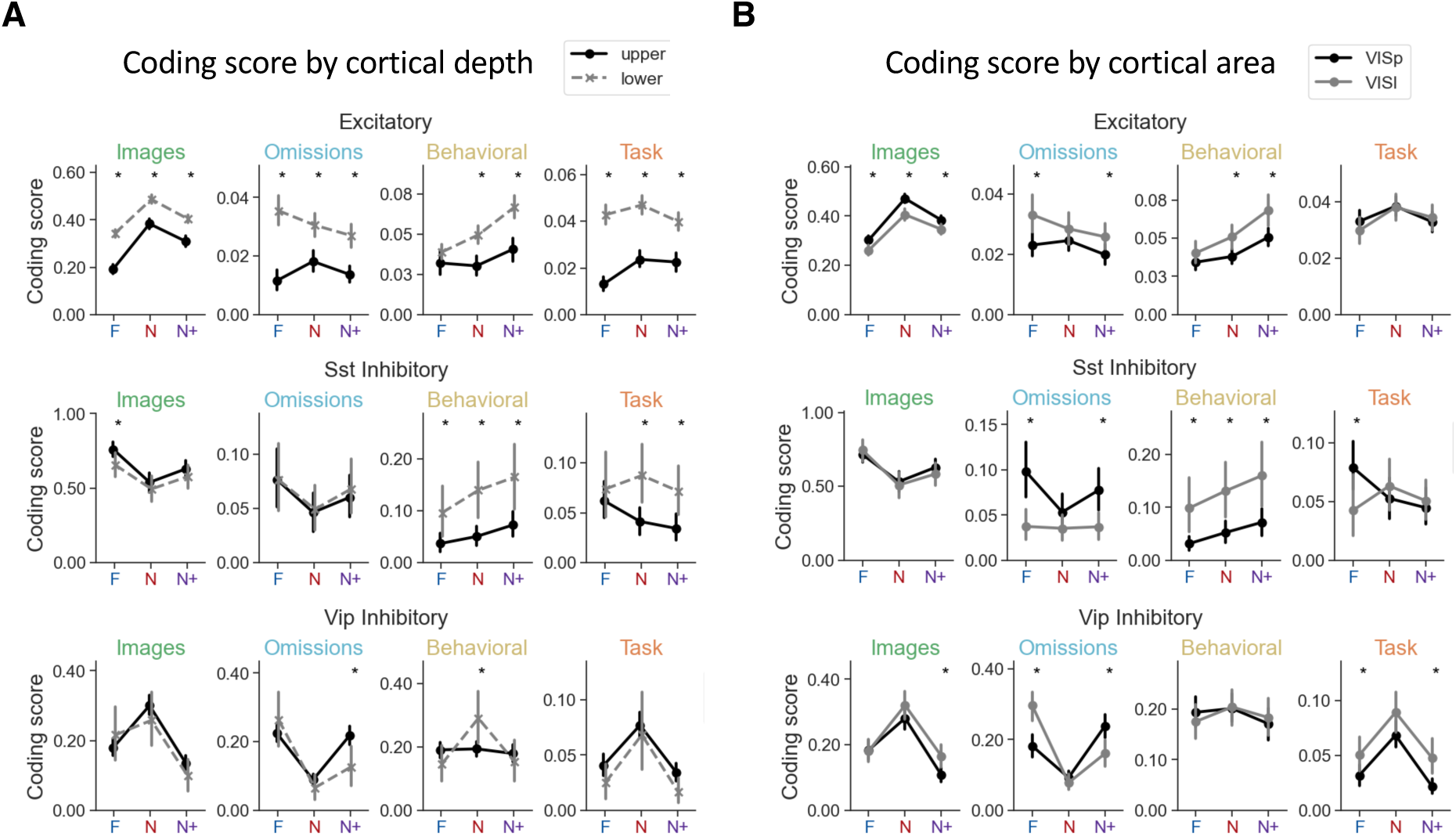
Coding scores by cortical area and depth. (**A**) Average coding scores for each cell type (rows) and feature group (columns) across experience levels, split by cortical depth (mean +/- 95% confidence intervals). Cells were divided into upper and lower layers of the cortex using a dividing line of 250um below the cortical surface. Only data from mice with multi-plane imaging (Cohort 3) are included. (**B**) Average coding scores for each cell class (rows) and feature group (columns) across experience levels, split by cortical area (mean +/- 95% CI confidence intervals). For all panels, only data from mice with multi-plane imaging (Cohort 3) are included. Differences between cortical areas or depths for a given experience level are marked as significant with the result of a two-sided independent sample t-test (p < 0.05).

**Figure S16.**
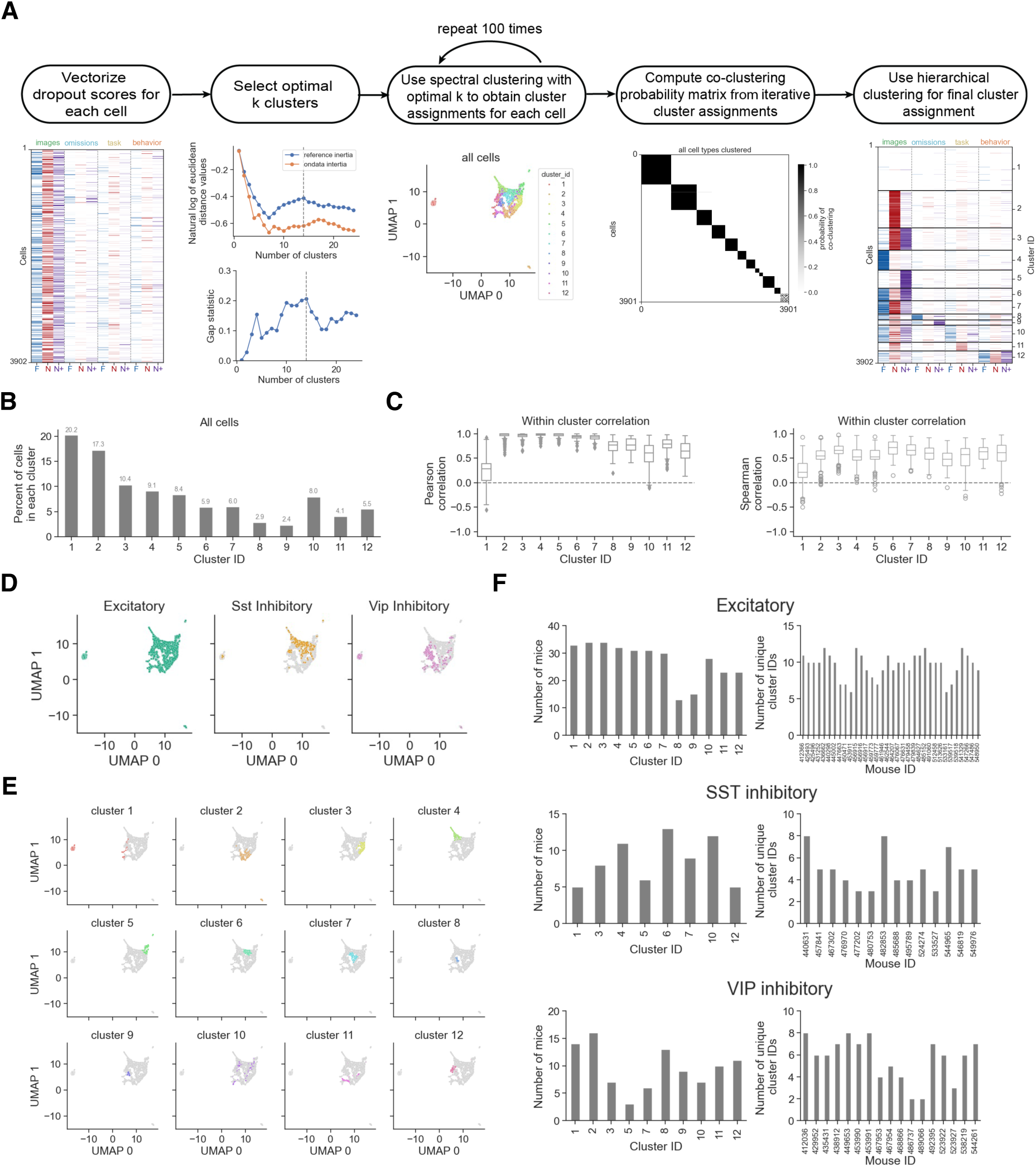
Clustering methods. (**A**) Schematic of steps used to cluster vectorized coding scores for cells matched across sessions. All cell types (excitatory, Sst inhibitory, Vip inhibitory) were included in clustering. Gap statistic was used to determine number optimal of clusters. Final clusters were determined based on hierarchical clustering on the co-clustering probability matrix to identify cells that reliably fell into the same cluster across iterations of the clustering algorithm. Clusters are ordered by their preferred coding feature (images, omissions, task, behavior). (**B**) Cluster sizes, shown as fraction of cells falling into each cluster. (**C**) Within cluster correlation, computed as the Pearson (left) or Spearman (right) correlation of each neuron’s vectorized coding scores with the cluster average, demonstrating that clusters are self-consistent. (**D**) Data visualized after dimensionality reduction using Uniform Manifold Approximation and Projection (UMAP) of all matched cell coding scores, colored by cell class to demonstrate separability of Vip and Sst cell coding properties. (**E**) UMAP of all matched cell coding scores, separated and colored by cluster ID. (**F**) Left, number of mice that contributed to each functional cluster demonstrating that each cluster was comprised of cells from multiple mice. Right, number of clusters identified per mouse.

**Figure S17.**
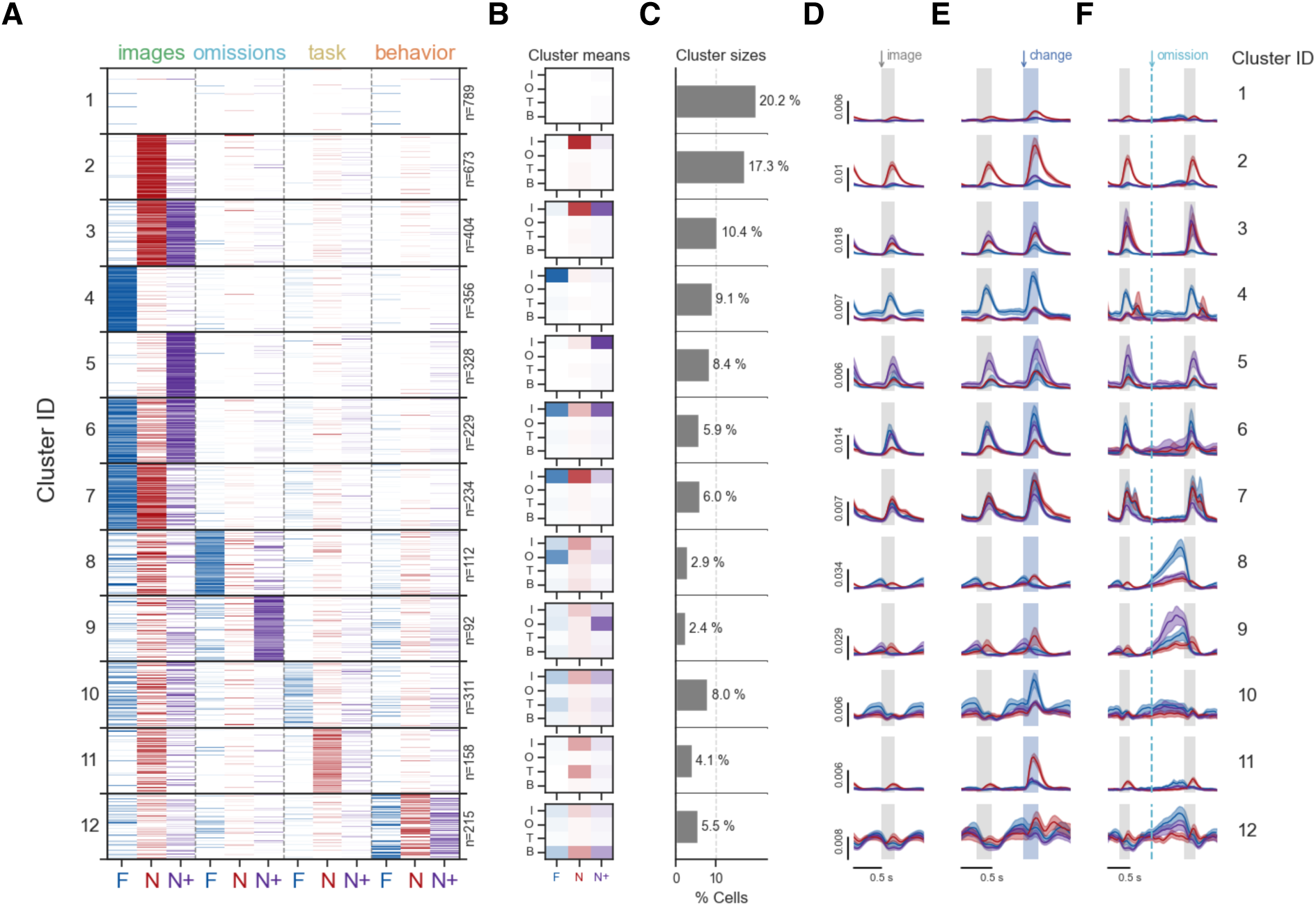
Cluster properties for all matched cells. (**A**) Clustered coding scores for all matched cells from all cell types (n = 3,298 excitatory cells,194 Sst cells, 409 Vip cells; total of 3,901 neurons from 131 fields o view). Clusters are sorted by preferred feature coding (image, omission, task, behavior). Color intensity represents strength of coding score, as in Fig. 5C. (**B**) Coding scores averaged across all cells within each cluster. Rows are feature groups (image, omission, task, behavior) and columns are experience levels (Familiar, Novel, Novel+). (**C**) Percent of cells belonging to each cluster, relative to the total number of matched cells across all cell types (n=3,901). Cluster sizes split by cell type can be found in Fig 5F. (**D**) Average response to non-change image presentations for each cluster, colored by experience level (red-Familiar, blue-Novel, purple-Novel+). (**E**) Average response to image changes across experience levels for each cluster. Gray bar is the pre-change stimulus, blue bar is the change stimulus. (**F**) Average response to image omissions across experience levels for each cluster. Dotted lines mark the time of omission. All cell types are included in the population average for each cluster. See figs. S20-S22 for coding scores and population responses split by cell type.

**Figure S18.**
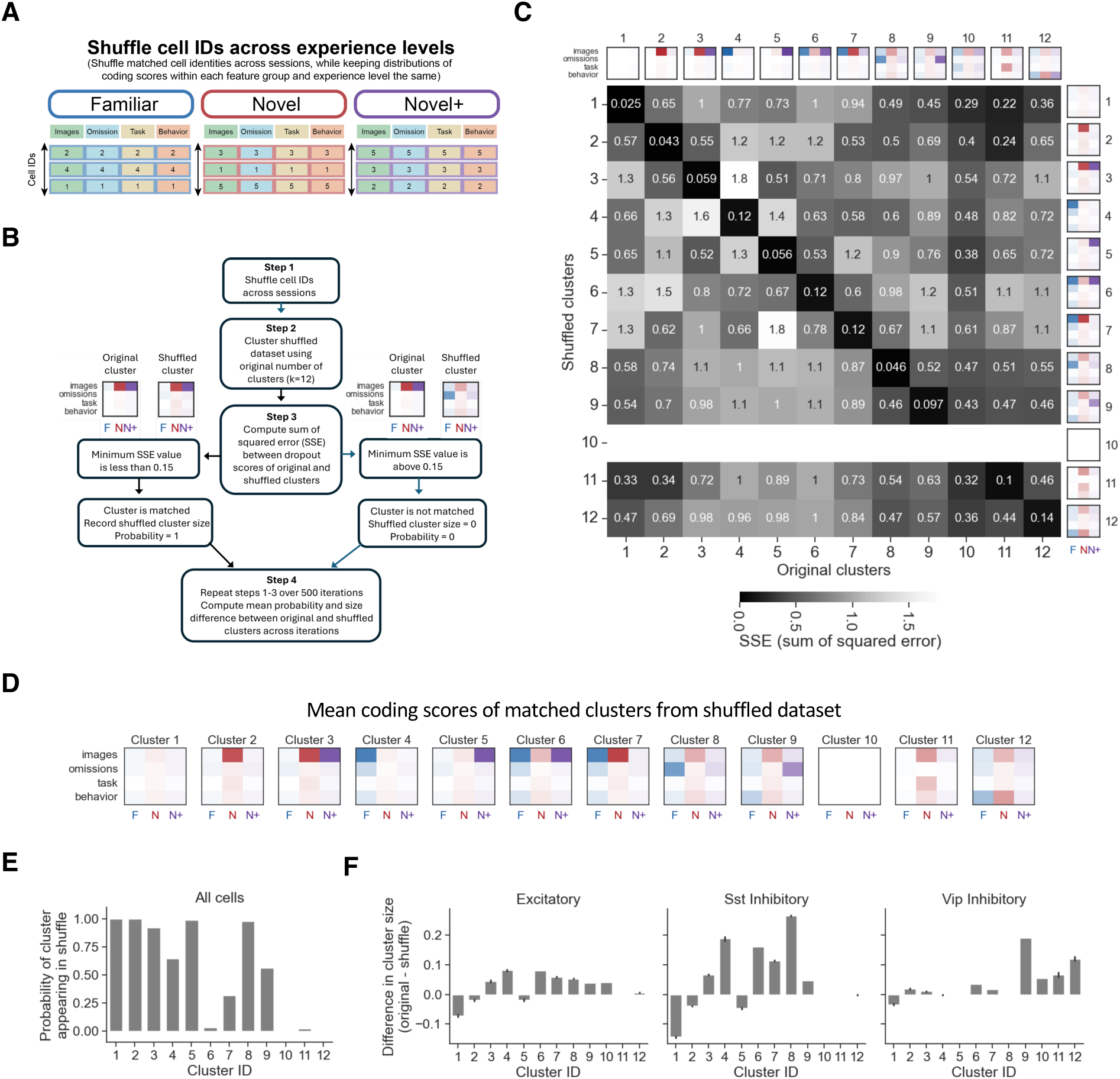
Shuffle control for experience dependent coding clusters. Shuffling cell IDs across experience levels results in smaller clusters for most experience dependent coding patterns. **A,** Schematic of shuffle procedure. Cell IDs were shuffled within sessions (experience levels) to disrupt single cell patterns of across day coding changes while maintaining the distribution of coding scores within each experience level and feature category. This allows comparison of original cluster patterns and sizes with the pattern of across day coding changes in the shuffled dataset. Numbers indicate hypothetical cell ids. **B,** Schematic of procedure for comparing original clusters with shuffled clusters. Clustering was run on shuffled dataset with the same value of k (number of clusters) over 500 iterations of the shuffle. In each iteration, shuffled clusters were matched to original cluster patterns based on minimizing the sum of squared error (SSE) between original and shuffled cluster coding scores. Shuffled clusters with < 0.1 SSE were considered as matched to the original cluster. The final step is to calculate the probability of a given cluster appearing in the 500 shuffle iterations, as well as the size (number of neurons) belonging to each shuffled cluster across iterations **C,** Mean SSE values comparing shuffled clusters (rows) to original clusters (columns). Rows with no values indicate that no shuffled clusters were found to match to the original cluster pattern. **D**, Mean coding scores of shuffled clusters matched to original clusters. Cluster 10 was absent in shuffled data. **E,** Probability of each cluster appearing in the set of shuffled clusters over 500 iterations. Clusters that did not appear in shuffled data have low or zero probability values. **F**, Difference in size of original clusters compared to shuffled clusters, computed as the difference in original cluster size and shuffled cluster size, averaged over 500 iterations (error bars show +/-95% confidence intervals). Positive values indicate more cells in original data clusters compared to shuffled clusters, negative values indicate fewer cells in original data clusters compared to shuffle.

**Figure S19.**
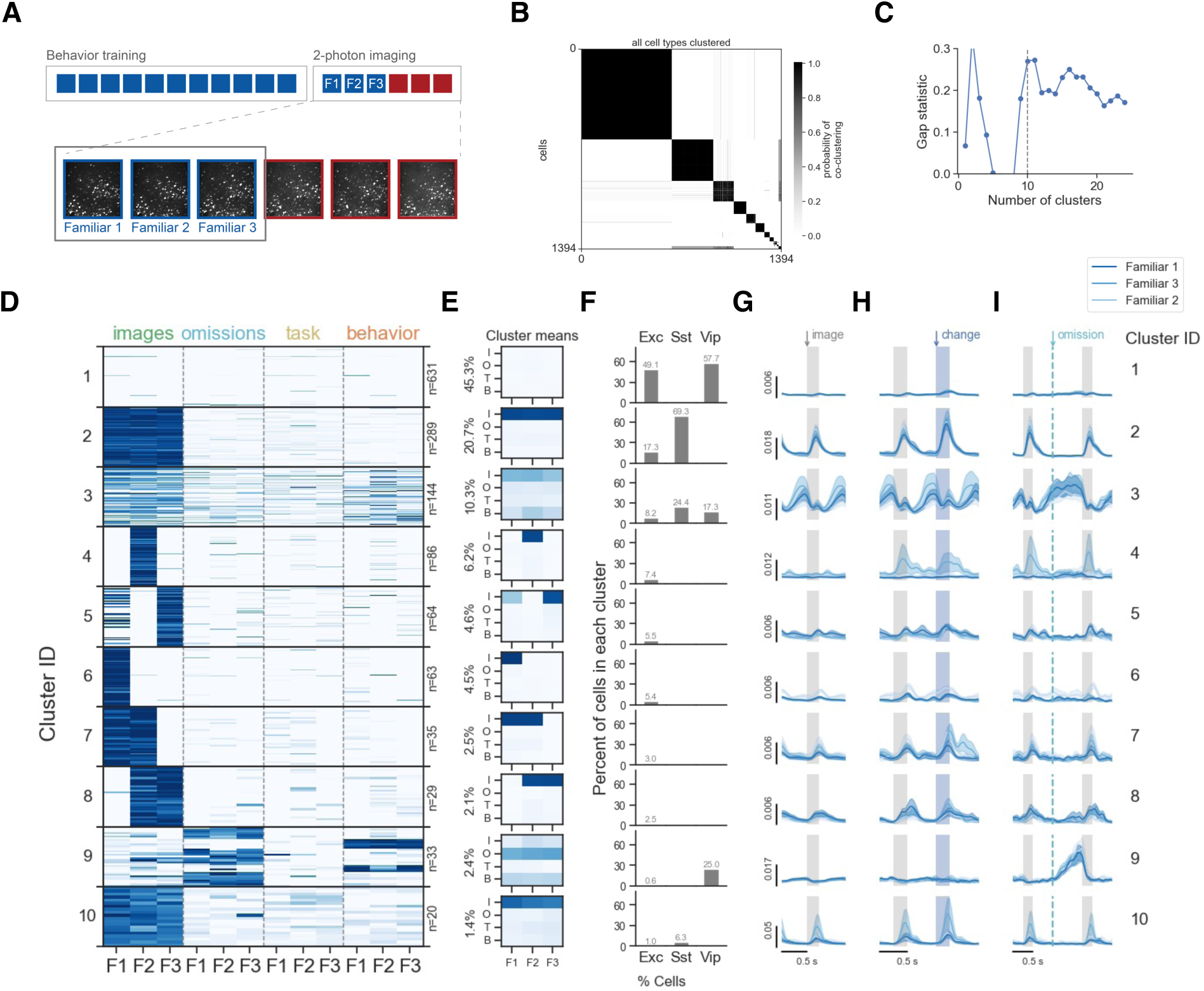
Clustering results using three sequential sessions with familiar images. Clustering on familiar sessions results in fewer clusters with more consistent coding across days compared to novelty-based clusters. (**A**) Schematic showing sessions were used for clustering. Matched cells from fields of view with 3 active behavior sessions with familiar stimuli were included in familiar-only clustering analysis (n = 1,163 excitatory cells, 127 Sst cells, 104 Vip cells; total of 1,394 cells from 52 fields of view from 23 mice). (**B**) Co-clustering probability matrix for clustering performed on cells matched across three Familiar sessions. (**C**) The optimal number of clusters was identified using eigengap values, yielding k=10 clusters (see Materials and Methods and Supplemental Text). (**D**) Clustered scores for cells matched in 3 Familiar sessions, sorted by cluster size., Y-axis shows individual neurons, X-axis shows feature categories (Images, Omissions, Behavioral, Task) across experience level (Familiar, Novel, Novel+). (**E**) Coding scores averaged across matched cells within each cluster. (**F**) Cluster sizes within each cell type (excitatory, Sst inhibitory, Vip inhibitory), shown as the percent of cells within each cell type belonging to each cluster. (**G**) Average response to non-change image presentations for each cluster. Shading represents the 3 Familiar sessions. (**H**) Average response to image changes across experience levels for each cluster. Gray bar is the pre-change stimulus, blue bar is the change stimulus. (**I**) Average response to image omissions across experience levels for each cluster. Dotted lines mark the time of omission. All cell types are included in the population average for each cluster.

**Figure S20.**
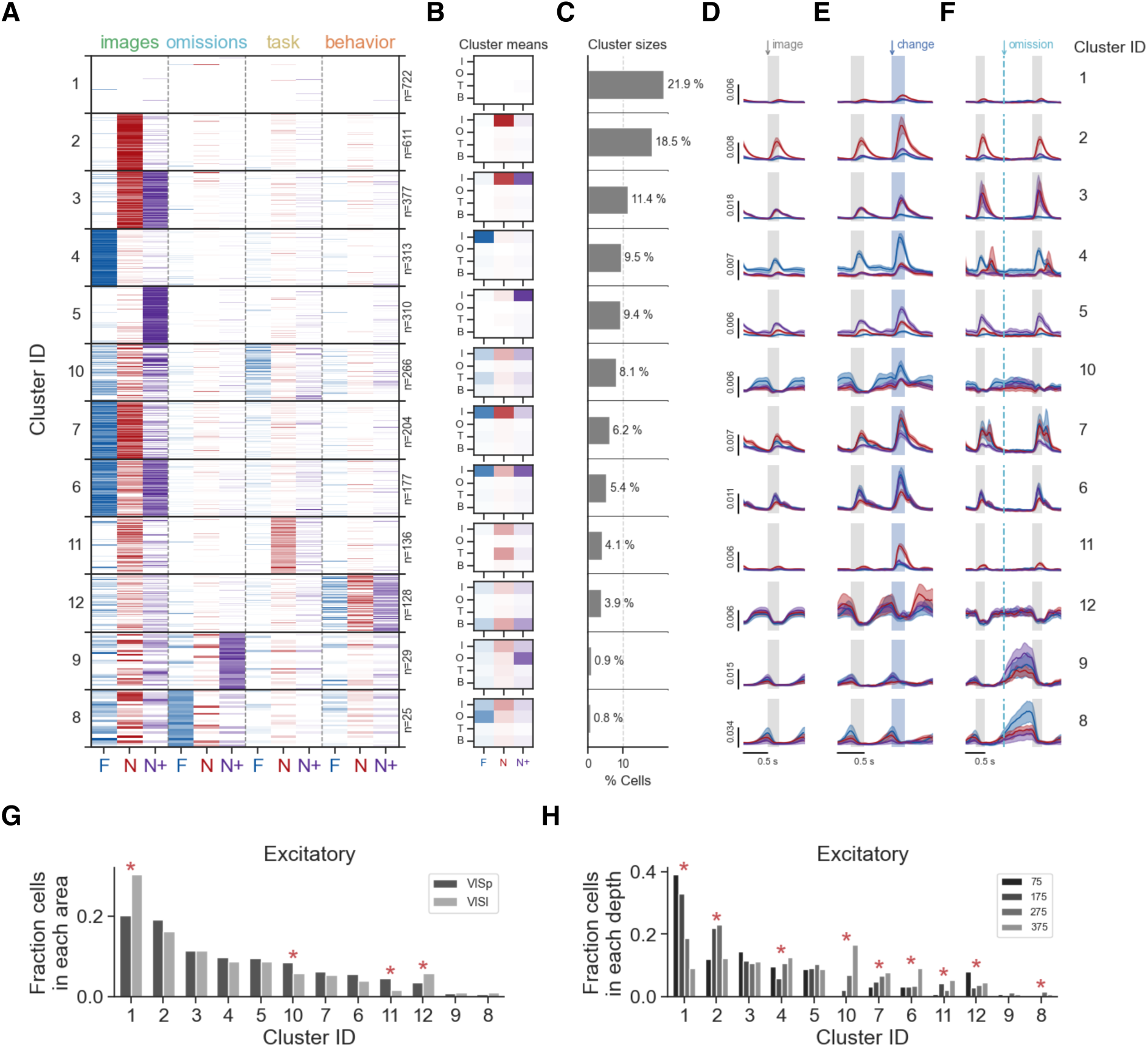
Cluster properties for excitatory cells. (**A**) Clustered coding scores for matched excitatory cells, organized by cluster size within the excitatory population. Color intensity represents strength of coding score, as in Fig. 5C. (**B**) Coding scores averaged across all excitatory cells in each cluster. Rows are feature groups (image, omission, task, behavior) and columns are experience levels (Familiar, Novel, Novel+). (**C**) Percent of excitatory cells belonging to each cluster. (**D**) Average response to non-change image presentations for each excitatory cluster, colored by experience level (red-Familiar, blue-Novel, purple-Novel+). (**E**) Average response to image changes across experience levels for each excitatory cluster. Gray bar is the pre-change stimulus, blue bar is the change stimulus. (**F**) Average response to image omissions across experience levels for each cluster. Dotted lines mark the time of omission. (**G**) Fraction of excitatory cells in each imaged visual area across clusters. Fraction is computed by normalizing the number of cells in each area for each cluster to the total number of excitatory cells in each area. Statistical comparison across areas was performed by chi-square test on the proportion of cells in each area for each cluster compared to the overall proportion of cells in that cluster. (**H**) Fraction of excitatory cells in each imaging depth across clusters. Statistics were performed as described in panel G (also see Materials and Methods).

**Figure S21.**
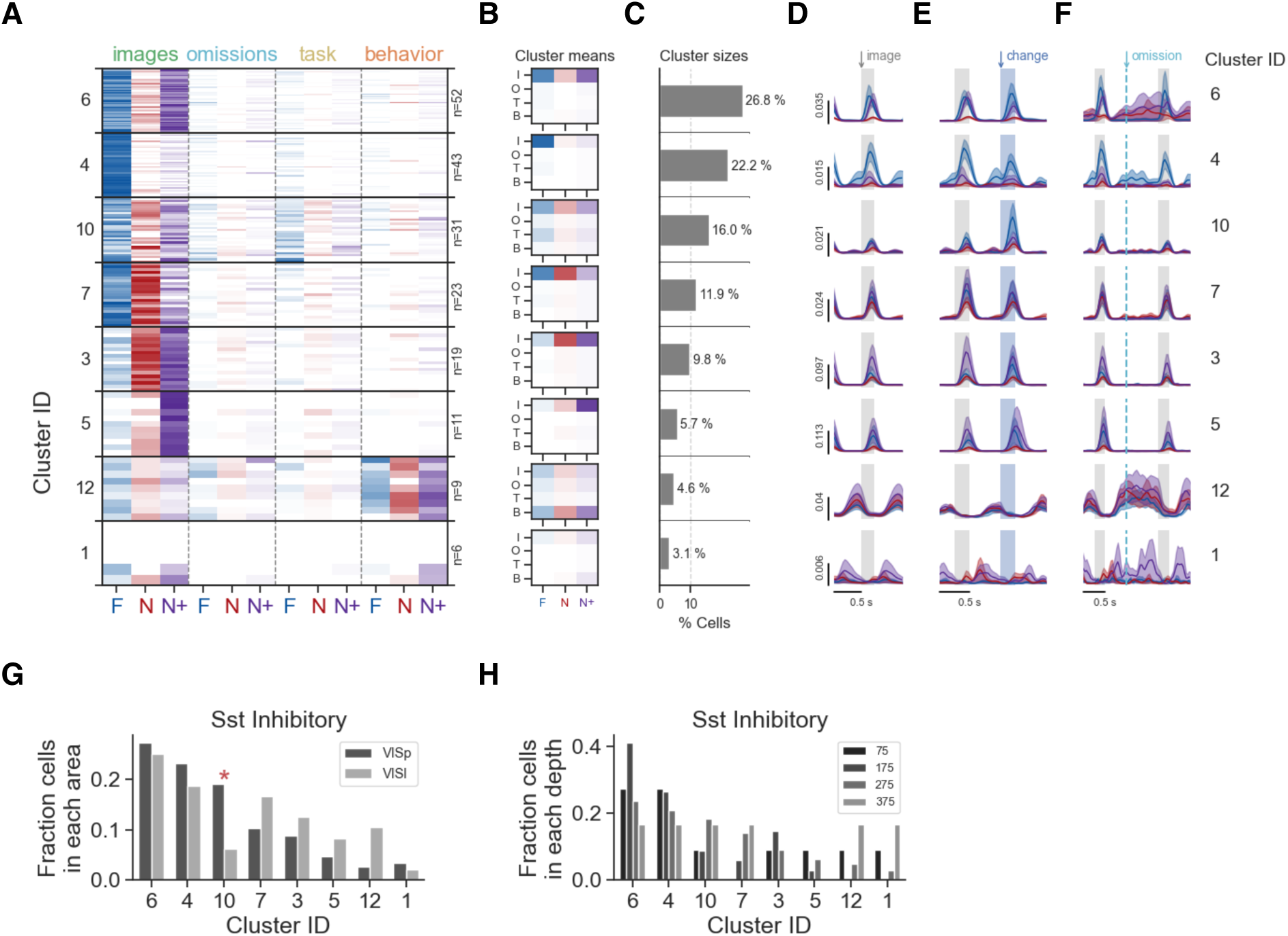
Cluster properties for Sst cells. (**A**) Clustered coding scores for matched Sst inhibitory cells, organized by cluster size within the Sst population. (**B**) Coding scores averaged across all Sst cells in each cluster. Rows are feature groups (image, omission, task, behavior) and columns are experience levels (Familiar, Novel, Novel+). (**C**) Percent of Sst cells belonging to each cluster. (**D**) Average response to non-change image presentations for each Sst cluster, colored by experience level (red-Familiar, blue-Novel, purple-Novel+). (**E**) Average response to image changes across experience levels for each Sst cluster. Gray bar is the pre-change stimulus, blue bar is the change stimulus. (**F**) Average response to image omissions across experience levels for each Sst cluster. Dotted lines mark the time of omission. (**G**) Fraction of Sst cells in each imaged visual area across clusters. Fraction is computed by normalizing the number of cells in each area for each cluster to the total number of Sst cells in each area. Statistical comparison across areas was performed by chi-square test on the proportion of cells in each area for each cluster compared to the overall proportion of cells in that cluster. (**H**) Fraction of Sst cells in each imaging depth across clusters. Statistics were performed as described in G (also see Materials and Methods).

**Figure S22.**
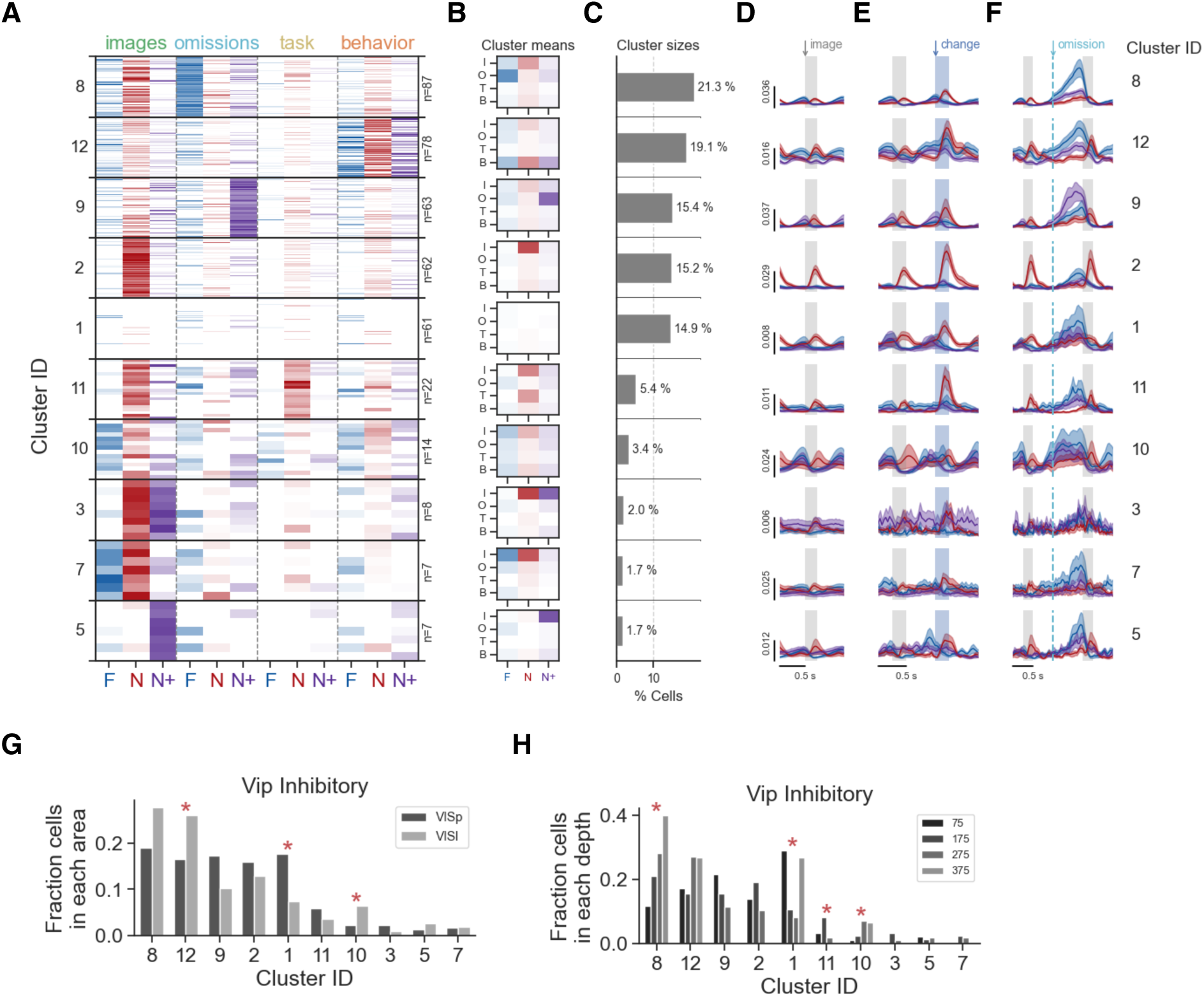
Cluster properties for Vip cells. (**A**) Clustered coding scores for matched Vip inhibitory cells, organized by cluster size within the Vip population. (**B**) Coding scores averaged across all Vip cells in each cluster. Rows are feature groups (image, omission, task, behavior) and columns are experience levels (Familiar, Novel, Novel+). (**C**) Percent of Vip cells belonging to each cluster. (**D**) Average response to non-change image presentations for each Vip cluster, colored by experience level (red-Familiar, blue-Novel, purple-Novel+). (**E**) Average response to image changes across experience levels for each Vip cluster. Gray bar is the pre-change stimulus, blue bar is the change stimulus. (**F**) Average response to image omissions across experience levels for each Vip cluster. Dotted lines mark the time of omission. (**G**) Fraction of Vip cells in each imaged visual area across clusters. Fraction is computed by normalizing the number of cells in each area for each cluster to the total number of Vip cells in each area. Statistical comparison across areas was performed by chi-square test on the proportion of cells in each area for each cluster compared to the overall proportion of cells in that cluster. (**H**) Fraction of Vip cells in each imaging depth across clusters. Statistics were performed as described in G (also see Materials and Methods).

**Table 1.**
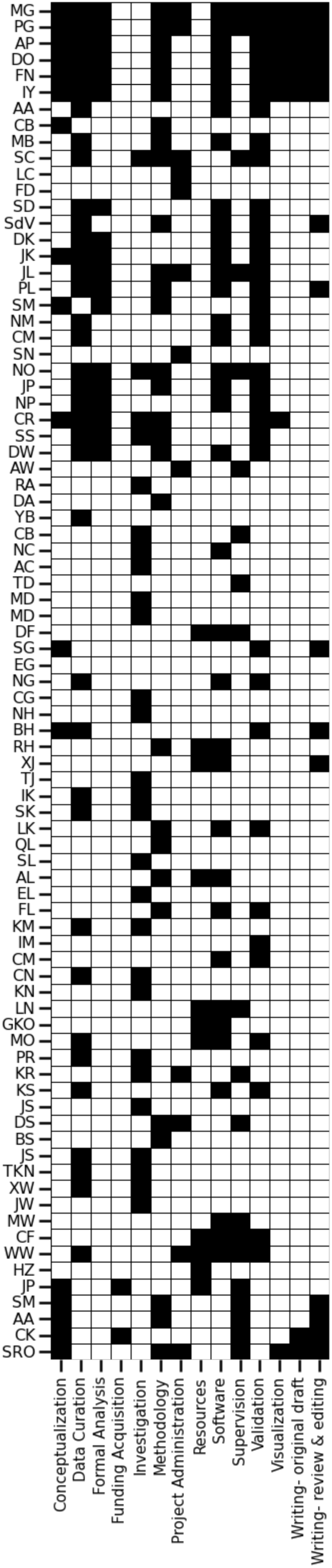
Author contributions table using CRediT taxonomy.

## Supplementary Materials

### Supplementary Text

### S1. Allen Brain Observatory Visual Behavior 2-Photon Dataset

This section of the supplementary text and associated figures are intended to provide a guide to the features of open access dataset used in this study. The selection criteria and rationale for inclusion in analysis are described in a subsequent section.

#### Features of the in vivo optical physiology dataset: recording conditions, cohorts, and QC

The Visual Behavior 2-Photon dataset was designed to address questions about the impact of experience, expectation, and task engagement on neural coding and dynamics during in the mouse visual cortex. The central experimental design involved tracking populations of neurons across multiple days with varying sensory and behavioral contexts, including active behavior sessions with familiar or novel images, and passive viewing sessions where the same stimuli were displayed in the absence of reward. The behavior task was a visual change detection task with repeated images, unexpected image changes, and unexpected stimulus omissions. All in vivo recordings were conducted in well trained mice. We surveyed the activity of thousands of excitatory and inhibitory neurons across multiple cortical depths and visual areas. Raw data were processed through a standardized pipeline, carefully curated, and packaged into NWB format. We have made the full dataset openly available for scientific inquiry, alongside a Python based Software Developer Kit (AllenSDK) for easy access of experimental data and metadata (https://allensdk.readthedocs.io/en/latest/visual_behavior_optical_physiology.html).

The primary dimensions of the dataset are defined by 1) the transgenic lines used to target specific genetically defined cell populations, 2) the cortical areas and depths that were imaged, and 3) the different session types and image sets that were shown (fig. S1). Depending on the question of interest, different subsets of the data can be selected using the metadata tables provided by the AllenSDK.

The dataset consists of four cohorts of mice, each with a unique recording configuration and set of stimuli shown during task performance (defined by their “project_code” in the AllenSDK metadata tables; fig. S2). In all cohorts, data was acquired from transgenic mice expressing GCaMP6 in excitatory, Sst inhibitory, or Vip inhibitory neurons. The majority of the dataset was collected using a GCaMP6f reporter line; a small subset of excitatory recordings used GCaMP6s. These can be distinguished by the “full_genotype” in the metadata tables (fig. S2D).

The first two cohorts were imaged using traditional single plane in vivo 2-photon calcium imaging in the primary visual cortex (fig. S2). The two single plane cohorts are distinguished by which image set was shown during task learning (the familiar image set), and which was shown for the first time during 2-photon imaging (the novel image set). In the *VisualBehavior* cohort of mice (cohort 1), image set A was showing during training, and image set B was the novel set shown only during imaging. The *VisualBehaviorTask1a* group (cohort 2) used the opposite configuration – image set B was shown during training and image set A was the novel set. The behavior and physiology measured in these two cohorts can be compared to dissociate differences due to stimulus novelty from differences in the specific features present in the image sets (fig. S4E-H; fig. S7; fig. S8).

We acquired data from additional groups of mice using a modified multi-plane Mesoscope 2-photon RAM system, which allowed near simultaneous recording across multiple visual areas and cortical depths (fig. S2). Up to 8 imaging planes could be recorded in each session. In the *VisualBehaviorMultiscope* group of mice (cohort 3), we targeted 4 cortical depths (75, 175, 275, 375 um) spanning upper layer 2 to upper layer 5, in two visual areas, the primary visual cortex (VISp, also called V1) and a secondary visual area (VISl, also called LM). The stimuli used in this cohort overlap with the *VisualBehavior* cohort; image set A became familiar during training and image set B was shown as the novel set during in vivo imaging. The data from group of mice can be used to study information flow and interactions across cortical areas and depths.

In the final cohort of mice (indicated by the “project_code” *VisualBehaviorMultiscope4areasx2d*, cohort 4) we made a key modification to the stimulus design – two of the images in the familiar set (image set G) were included in the novel image sessions during in vivo imaging. Thus image set H consisted of 6 novel images and two 2 familiar images that shared with image set G. This manipulation allows comparison of neurophysiology and behavior in response to familiar stimuli in a familiar context with familiar stimuli in a novel context. We also used a different recording configuration for this cohort, targeting 2 cortical depths (175 & 275 um) across 4 visual areas (VISp, VISl, VISal, VISam). This enables study of differences across hierarchically organized regions, with VISam at the top of visual hierarchy defined by connectivity, VISp at the bottom, and VISl and VISal at intermediate levels (*21*). Given the distinct goals and design of this portion of the dataset, this cohort was not included in the analysis for the current study.

The in vivo 2-photon imaging recordings were subject to stringent quality control (QC) criteria to ensure that only the highest quality data was included in the final dataset for analysis (fig. S1). QC was performed at the level of individual imaging planes within a session (defined by their unique “ophys_experiment_id”), entire recording sessions (which can contain multiple imaging planes, identified by a shared “ophys_session_id”), and the collection of multi-day recordings for a given field of view (indicated by a shared “ophys_container_id”). Critically, each session was assessed for drift in the z-plane of imaging over the session and removed from analysis if excessive drift occurred, which could result in loss of cells from the imaging plane. Accuracy of cell matching across sessions for a given field of view was another important criterion, and sessions that were not well matched (i.e. differences in the z-plane over days resulting in different sets of cells) were also removed from analysis. See Materials and Methods for detailed description of QC criteria and procedure.

Our aim was to acquire one passing session of each type (fig. S1F; two familiar active sessions with one interleaved passive day, and two novel active sessions with one interleaved passive day). If a session of a given type failed QC, an attempt was made to re-acquire that session. As a result, the dataset for a given field of view (united across sessions by its “ophys_container_id”) may not include all session types, and the dataset for an individual multi-plane recording session may not have all 8 imaging planes, as each imaging plane could fail QC independently. However, the behavior performance and stimulus display information is available for all sessions, both during training and during the ophys portion of the experiment, irrespective of whether the ophys data itself passed QC or not. This allows the full history of each mouse to be assessed and is provided in the “behavior_sessions” table accessible via the AllenSDK.

Accordingly, it is important to note that the “session_type” designation in metadata tables does not indicate the order in which the sessions were acquired. Instead it indicates the intended session type, and the session order must be determined from the full sequence of recording sessions provided in the “behavior_sessions” table. This is particularly important when addressing questions of novelty and experience dependence. The AllenSDK metadata tables provide several useful columns indicating the number of prior exposures to a given image set, session type, and prior exposures to omissions, which were only shown during imaging.

#### Relationship of imaging depth and cortical layer

We targeted our imaging to specific cortical depths based on known expression patterns for each cell type (*22*). In the single plane recordings, we targeted excitatory neurons at 175 and 375 um depths, Vip neurons at 175 um, and Sst neurons at 275 um. In the multi-plane recordings, we targeted 75, 175, 275, 375 um depths in each recording session. For Vip and Sst transgenic lines, few cells were observed at the 375um depth, likely due to a combination of lack of expression at this depth (particularly in the case of Vip, which is primarily expressed in superficial layers) and reduction in signal quality with deeper imaging. Fields of view at the 375um depth with fewer than 3 detected ROIs were excluded from analysis.

In the visual cortex of the mouse brain in the absence of any surgical implants, layer 1 spans ∼80um and layer 5 begins at ∼400um depth from the cortical surface. Thus, the 75um depth would correspond to layer 1 and the 375um depth would correspond to cortical layer 4. In our cranial window preparation, the cortex is compressed by ∼20% due to a flat glass coverslip being implanted over the surface of the naturally curved brain (*22*). This mild compression aids in stabilizing the brain under the window, minimizing brain motion. Accordingly, with a ∼20% compression of the cortex, layer 1 would span ∼65um and layer 5 would begin at ∼320um in depth. Thus, we estimate that our imaging depths span upper layer 2 (∼75um depth) to upper L5a (∼375um depth), including layer 3 (∼175um depth) and layer 4 (∼275um depth). Measurements of cell size (number of pixels per segmented ROI) corroborate this estimation, as excitatory neurons increase in size in imaging planes beginning around 350um, consistent with an increase in cell size in layer 5 compared to layer 4 (fig. S2F).

It is important to note that there can be variability in the amount of compression in each animal due to surgical and biological variability, thus these values are estimates and should not be taken as verified measures of cortical layer. Indeed, there is variability in measured cell sizes across animals within a given depth, indicating that some recording depths could contain a mixture of neurons from multiple layers. In particular, recordings at the 375 um depth show a bimodal distribution of small and larger neurons, indicating a mixture of cells from layers 4 and 5.

### S2. Details of change detection paradigm and training procedure

The full dataset includes the full training history of 107 mice, which can be analyzed independently from the physiology data to understand task learning. For example, a recent study using this dataset examined the behavioral strategies employed by mice learning the change detection task (*23*). In the current study, we selected mice with specific types of ophys sessions (see section on selection criteria below), resulting in 66 mice included in the study. The training history and performance across training stages for these 66 mice is included in fig. S3 and fig. S4. The task structure, training procedure, and controls are described below.

#### Task structure and trial types

The change detection task consists of a continuous stream of stimuli, presented over the course of 1 hour. There is no explicit trial structure provided to the mouse (i.e. no trial start cue), however there is an implicit, experimenter defined structure to the task (fig. S3A). Each “trial” based on this definition consists of a collection of individual stimulus presentations, and the transition between trials depends on the animal’s behavior, as described in detail below and in fig. S3.

At the start of the experimenter defined trial, an initial stimulus will be displayed, and a change time will be drawn from an exponential (for the first training stage) or geometric (for all other training stags) distribution between 3 and 9 seconds or 4 and 12 stimulus flashes, respectively (fig. S3C). This distribution was selected to provide a flat hazard rate, meaning the probability of a change occurring at a given time is constant regardless of how much time has elapsed during the trial. This is intended to prevent the mouse from predicting the change time. If the mouse emits a false alarm lick prior to the scheduled change time, the trial is considered “aborted” and will reset, using the same selected change time, for up to 5 trials in a row (or until the trial is not aborted). This is intended to discourage the mouse from licking early and require that they wait to respond until the change has occurred. If the mouse correctly withholds licking until the scheduled change time, they have a [150, 750] ms window after the change time to emit a lick to earn a water reward. This means that mice must lick before 750ms has elapsed; however, given the visual response latency of neurons in V1 (∼50ms, see Figure 2A,B of (*24*)) and the time required to initiate movement (∼100ms), the first 150ms after change stimulus onset are considered guesses (i.e. false alarms / aborts) and do not elicit a reward. After the 750ms reward window, mice are given a 3 second “grace period” to consume the water reward, during which no change can occur (fig. S3A, “consumption window”). The grace period occurs regardless of whether the mouse correctly licks (hit trial) or not (miss trial) after the change. The next trial then begins at the end of the grace period, and a new change time is drawn.

As a result of these task rules, the sequence of stimuli that are displayed, and specifically the time between image changes, depends on the licking behavior of the mouse. For instance, if a mouse is completely disengaged from the task and does not lick at all during a session (or if the mouse experiences a “passive viewing” session with the reward spout removed), image changes will be displayed purely based on the change time distribution, at intervals ranging from 3 to 10 seconds (for the first training stage), or 4 to 12 image presentations (in all other stages). This is equivalent to running the task in “open loop” mode, where rewards have no bearing on the task progression. In contrast, if a mouse reliably licks every 2 seconds, which is before the shortest possible change time, trials will be repeatedly aborted and the stimulus will never change. The ideal task performance would be if the mouse only licked in the 750ms window after image changes. Actual mouse behavior is a blend of these 3 scenarios. A detailed analysis of the behavioral strategies used in this task are described in (*23*, *25*). In short, some mice are prone to impulsively or predictably licking at regular, short intervals (approximately centered on the shortest possible change time), resulting in many aborted trials and long inter change intervals (yet still earning enough rewards by chance to make this “timing” strategy a viable option), while others successfully withhold licking and wait for image changes to occur (i.e. using the intended strategy of detecting image changes).

##### Image change selection

When a change time is drawn at the start of a trial, a change stimulus is also selected from an n x n transition probability matrix, where n is the number of stimuli being shown in that session (ex: 8 natural scene images, or 4 grating orientations) (fig. S3D). The initial stimulus will continue to be displayed until the selected change time, at which point the identity of the stimulus changes. Accordingly, if 8 images are shown during a session, the probability of image a changing to image b on a given trial is 1/64. Trials where the image identity changes are “go” trials (fig. S3D). On some trials, the change image that is drawn will be the same as the initial image, thus the image identity will not change. These trials can be considered as “catch” trials (fig. S3D) and used to compute the false alarm rate (i.e. the fraction of no-go trials where a mouse incorrectly licks when no change occurs). However it is important to note that the vast majority of stimulus presentations in a session are non-changes (fig. S3A), and could also be used to compute the false alarm rate. The definition of a “catch” trial described here is a conservative one, limited only to non-changes (no-go trials) that are drawn from the same time distribution as stimulus changes (go trials). A maximally liberal definition would be to compute false alarm rate across all non-changes in a session; however this would include repeated non-change image presentations that occur during reward consumption window, or during the time at the start of the trial when the stimulus can never change (prior to the minimum change time of 3 seconds or 4 image presentations). An intermediate definition would be to compute false alarm rate across all non-changes that *could* have been a change, based on the change time distribution and task rules. This is the definition we have used in the current study, and code is provided to identify this set of stimulus presentations based on the task rules <u>here</u>. Response rates for change and non-change stimuli across the 66 mice included in this study are shown for the final training stage in fig. S3B,E.

##### Image omissions

Image omissions occur randomly (∼5% of non-change image presentations are omitted) during ophys sessions (omissions never occur during training). The image presented before and after the omission is always the same image; thus image omissions are akin to an very long inter-stimulus interval (1250ms instead of the typical 500ms), and could be considered as a very challenging catch trial. Licks during omissions are never rewarded but licking during omissions can delay the onset of the next trial (as can licks occurring anywhere outside the behavioral response window and the post-change reward consumption period). Interestingly, mice are more likely to false alarm lick in response to non-change image presented just after an omission (fig. S3B; fig. S4D), rather than during the omission. This indicates that mice to do not perceive the omission as a “change” and instead treat it as a long catch trial.

#### Training procedure and session types

Mice learn the task through a series of automated training steps, beginning with a simple version of the change detection task and adding layers of complexity across successive stages based on each animal’s performance (i.e. mice progressed at their own pace). Below we describe the stimuli, reward volumes, and transition criteria for each stage of the task, along with their rationale. The training stages are also illustrated in fig. S3F.

##### Behavioral shaping

While the task itself is an operant task (mice are rewarded for licking a spout following an image change, i.e. rewards are contingent on mouse behavior), the very first day of training consists of a 15-minute associative pairing session where rewards were automatically delivered following stimulus changes. The goal of this stage was to form an initial stimulus-reward association under the simplest possible conditions, using high contrast, oriented grating stimuli with no gray screen interval. In each trial, a grating was continuously presented and changed in orientation from 0 to 90 degrees (or 180 or 270 degrees, which are the same orientation but opposite phase of the grating), resulting in highly salient changes paired with 10ul rewards. This session is referred to in the AllenSDK as “session_type” = “TRAINING_0_autorewards”.

In the next stage of the task (TRAINING_1_static_gratings), the stimuli and reward volume were the same (static grating orientation changes, 10ul rewards) but reward delivery was now contingent on the animal deciding to lick following a stimulus change. Mice remained in this training stage until their performance, computed using the d-prime discriminability metric across go and catch trials (z-score of distribution of hit and false alarm rates), was >2 for 2 out of 3 consecutive days.

Once performance reached criterion in the static gratings phase, a gray screen interval was added, making the task more challenging and incorporating a working memory component. In this stage (TRAINING_2_gratings_flashed), and all subsequent stages, stimuli were presented for 250ms followed by a 500ms gray screen interval. With the gray screen interval, the task of mice is to determine whether the image that is currently being presented is the same or different from the one presented 500ms ago.

After reaching criterion performance with flashed gratings (2/3 consecutive days with d-prime >2), mice transitioned to the performing the task with natural scene images. The initial image change detection training stage (TRAINING_3_images_10uL_reward) began with a 10ul reward volume, consistent with all prior stages. After 3 sessions with natural images were completed, the reward volume was reduced to 7uL. The initial 10uL reward was intentionally large to motivate mice to learn the initial task. Once mice learned the task rules, the reward volume was reduced to encourage mice to complete more trials per session.

Mice remained in the flashed natural image stage of the change detection task until they reached criterion of having 2/3 consecutive days with a d-prime > 1 and at least 100 engaged trials in all 3 sessions. “engaged” trials were defined based on the animals rolling reward rate, computed as the average over 10 image change trials across the session, with a threshold of 2 rewards per minute for inclusion as an “engaged trial”.

Once mice reached criterion performance for flashed natural images, mice were considered as “handoff ready” and could be transitioned to the *in vivo* 2-photon imaging portion of the experiment. Mice could stay in “handoff ready” state for a variable amount of time, depending on availability of the microscope.

Accordingly, if one wished to compute “days to criterion performance”, this should be calculated as time to reach “TRAINING_4_images_handoff_ready”, rather than time to reach the “OPHYS” stage of the experimental procedure. It is also important to note that mice could regress from “handoff_ready” to “handoff_lapsed” if their performance dipped below the criterion level over a sequence of 3 days, and return back to “handoff_ready” if it reached criterion again.

The full training sequence for all mice is shown in fig. S2H, along with quantification of the number of days in each stage and behavior performance in each stage across mice from each transgenic line in fig. S2I,J.

The full set of task parameters and transition requirements between training stages are described here: https://github.com/AllenInstitute/mtrain_regimens/blob/ophys_task1a_master/regimen.yml

The transition criteria between training stages are defined here: https://github.com/AllenInstitute/mtrain/blob/master/mtrain/criteria.py

##### Behavior during optical physiology

After transitioning to the OPHYS portion of the experiment, mice received up to 3 “habituation” sessions where they performed the image change detection task under the microscope, but no physiology data was recorded. This was intended to acclimate the animal to the experimental setup, which included new sounds, smells, a different enclosure and general surroundings, and often a new trainer.

In the OPHYS phase, mice were no longer progressed between stages based on their performance but instead based on whether the physiology data for each stage passed data integrity checks and quality control (QC) criteria, ensuring that all necessary data streams were present and the data was of sufficient quality for inclusion in analysis. Accordingly, specific session types could be repeated a variable number of times during the OPHYS stage, with the goal of acquiring at least one ophys session of each type for potential inclusion in the final dataset. Importantly, all behavior data for ophys sessions is available and provided in the AllenSDK, only ophys data streams were subject to QC criteria.

The OPHYS session types included “OPHYS_1_images_A”, “OPHYS_2_images_A_passive”, “OPHYS_3_images_A”, which were all sessions with the same natural scene images that were shown during training (image set A), along with “OPHYS_4_images_B”, “OPHYS_5_images_B_passive”, and “OPHYS_6_images_B”, which were sessions with a new image set that had not been seen prior to imaging (image set B). The intent was to collect sessions with a target sequence that interleaved active behavior and passive viewing sessions, in the order shown in fig. S1D. Passive viewing sessions were those in which the reward spout was retracted and mice viewed the stimulus in “open loop” mode (mice were also given their daily water allocation prior to the passive sessions to ensure that they were satiated and no longer motivated to do the task). These were included to assess the impact of task engagement on neurophysiology. The ideal sequence would have moved through OPHYS stages 1 through 6, in that order. However in practice, due to QC criteria and retakes, sessions often occurred in differing orders in different animals. For example, if a mouse underwent sessions OPHYS 1 through 3, and OPHYS_2_images_A_passive failed quality control for any reason, another OPHYS_2_images_A_passive session could be acquired.

The full ophys session sequence for mice included in the study, indicating which sessions were selected for inclusion in analysis, is shown in fig. S4A and described below.

### S3. Selection criteria and behavior for Familiar, Novel, and Novel + sessions

#### Selection criteria for this study

The present study evaluated the impact of novelty and familiarization at the session level during active behavior performance. Accordingly, we included the first 3 cohorts (“project_code” = [*VisualBehavior*, *VisualBehaviorTask1B*, *VisualBehaviorMultiscope]*) in our analysis, which had either familiar or novel images in each session and did not include the subset of data where familiar and novel images were interleaved within the same session (“project_code” = “VisualBehaviorMultiscope4areasx2d”). Passive viewing sessions were also excluded from the primary analysis. The specific sessions included in analysis for the current study across selected mice are shown in fig. S4A.

While the optical physiology dataset included multiple sessions with familiar or novel images, we focused on 3 specific sessions: the first session with novel images (Novel), a subsequent active behavior session with novel images (Novel +), and a familiar active behavior session (Familiar) (fig. S4A). It is important to note that the Novel + sessions was typically not the second session with novel images, due to the interleaving of active and passive behavior sessions (fig. S4B,C). Similarly, the Familiar session was not always directly before the Novel session, as there may have been an interleaved passive familiar session.

Our analysis required that the same field of view was recorded and passed QC for each of the 3 session types we evaluated, such that the activity of the same neurons could be tracked across days. The first exposure to novel images was particularly important, as it could only occur once. Accordingly, if the first session with novel images for a given field of view failed QC, all sessions from that field of view were excluded from analysis. The first novel session can be identified in the AllenSDK metadata tables (*ophys_sessions_table* and *ophys_experiments_table*) using the value of “prior_exposures_to_image_set” = 0. This sometimes resulted in all data from a given mouse being excluded (particularly in the case of single-plane imaging).

There were typically multiple options for sessions meeting the criteria for Familiar and Novel+, and we always selected the QC passing session that was closest in time to the first novel session (i.e. the last active behavior session with familiar images just before the first novel session, or the active behavior session with novel images just after the first novel session).

Applying these criteria resulted 201 imaging sessions from 134 fields of view in 66 mice. There were a total of 12,826 excitatory cells, 468 Sst cells, and 1,194 Vip cells (fig. S4I). The distribution of cells across transgenic lines, areas, and depths is shown in fig. S4J. A fraction of these were segmented and matched in all 3 sessions: 3306 Excitatory cells, 200 Sst cells, 415 Vip cells (fig. S4J, fig. S6).

The full set of ophys sessions acquired for each of these 66 mice is included in fig. S4A, illustrating the order in which sessions were acquired, which passed or failed QC, and which 3 sessions were selected according to the criteria for Familiar, Novel, and Novel+.

All analysis of animal behavior in this study was performed on the 66 animals selected based on the criteria described above; fig. S3G-J describes behavior training history, fig. S4D-H describes task performance during optical physiology recordings, and fig. S5 shows patterns of running, pupil diameter, and licking behavior across the 3 session types.

Metadata for the set of neurons and sessions included in this study is available in Table S2.

#### Generalization of behavior to novel images

To assess whether task performance and animal behavior was influenced by exposure to novel stimuli, we analyzed behavior of well-trained mice in the same session types that were selected for physiology analysis (Familiar, Novel, Novel+). No statistically significant differences were detected for any metric we examined, when comparing the distributions of metric values across sessions (Fig. 2F-H; fig. S4E-H; fig S5). Mice licked at similar rates during image changes, non-changes, omissions, and post-omission image presentations across experience levels (fig. S4D). Behavior performance, measured as the discriminability index d-prime, was consistent across experience levels on average (Fig. 2F), even when the identity of the familiar and novel image sets was reversed (fig. S4E-H), demonstrating robust generalization of behavior across image sets regardless of their image identity or novelty. While the differences were not significant, there was some variability in performance across individual mice (gray points in fig. S4E-H), which could be further evaluated in future analyses aimed at understanding individual differences in performance.

Measures of behavior including lick rate, running speed, and pupil width also did not significantly differ across experience levels. In fig. S5A-C, G-I, we aligned each data stream to the onset of image changes or image omissions, averaged across trials within each session, then averaged across mice. In figs. 5D-F, J-L, we took the average value of the lick rate, running speed, or pupil width in a 500ms window after images or omissions on each trial and averaged across trials within each session, then compared the distributions across experience levels.

While we did not find statistically significant differences in behavior across sessions (although note the slightly elevated pupil width in the first novel session), the pattern and dynamics of mouse behavior showed interesting features (fig. S5A-C, G-I). The change and omission aligned running trace and pupil traces indicate that both behavioral readouts are synchronized, on average, to the cadence of stimulus presentations (fig. S5B, H). The timing of these fluctuations indicates that mice slow their running in anticipation of stimulus onset, slowing to nearly a full stop after image changes, likely to enable reward consumption. Interestingly, the average running behavior during the image presentations just prior to image omissions differed, and did not slow in advance of stimulus onset, instead mice slowed after the stimulus presentation (fig. S5H). This may indicate that anticipatory slowing occurs at times when a change is more likely to be expected, given that omissions occur entirely randomly. During omissions, mice continued to run at a steady pace and did not slow until the next stimulus presentation, after which they slowed more than during a typical repeated image presentation (fig. S5H), consistent with the slight increase in lick rate during post-omission images (fig. S5G). These signatures indicate that mice are more likely to mistakenly lick for a non-change stimulus if the inter stimulus interval preceding it was longer than typical. Interestingly, this post-omission licking behavior is correlated with the animal’s behavioral strategy (see Figure 2E of reference (*23*)).

Pupil width also fluctuated relative to stimulus presentation times (fig. S5C), despite the images being luminance matched to the inter stimulus gray screen. The peak of this oscillation was slightly out of phase with the changes in running behavior; running peaked during the inter-stimulus interval and declined prior to image onset, while pupil width peaked during the stimulus presentations and declined during the inter-stimulus interval. This result may be surprising given the many studies that have described tight locking between running speed and pupil diameter (for a review, see reference (*26*)). However this may signify a similar cognitive process – arousal, attention, or anticipation of an upcoming stimulus, which may or may not be an image change and could thus lead to a reward. Mice may slow their running speed in anticipation of a stimulus, to make it easier to stop, lick, and consume reward, while at the same time showing an increase in pupil width, reflecting that same anticipation.

#### Reversed image set control

One of the primary goals of this dataset was to examine the impact of novelty and stimulus experience on neural representations and behavior, motivating the choice to train mice with a set of familiar images and evaluate performance and physiology with novel stimuli. An important consideration is whether any observed effects originated not in the novelty of the stimuli but in the particular features of the stimuli themselves. Neurons in the mouse visual cortex are selective for visual features and mouse behavior can depend on image features such as spatial frequency and image contrast. The images selected for inclusion in the familiar and novel image sets were intended to span a similar perceptual range, based on initial pilot studies testing mouse behavior with these stimuli (*25*, *27*). We also performed control experiments where a subset of mice were trained with the reversed image sets – i.e. the set that was novel for most mice was the familiar training set for the control cohort, and the set that was familiar for most mice was the novel set for the control cohort. This group of mice has the AllenSDK identifier (aka “project_code”), *VisualBehaviorTask1B*, and is referred to as “Cohort 2” in figs. S2, S4, S7, and S8.

Comparison of behavior performance across cohorts demonstrated consistent performance when comparing across experience levels within each cohort (fig. S4E,F) and when comparing the same experience level across cohorts (fig. S4G,H), indicating that the identity of the images within a given set was not a factor in driving the experimental results we observed.

Importantly, the consistency of behavioral performance, and metrics of behavior including running, pupil, and licking behavior, across Familiar, Novel, and Novel+ sessions are important to consider in contrast to the changes in physiology within the visual cortex, described in the section below; differences in behavior cannot trivially account for any observed differences in physiology.

### S4. Characterizing Neural Activity

The following section describes analysis of basic response properties across cell types (Excitatory, Sst Inhibitory, Vip Inhibitory) and experience levels (Familiar, Novel, Novel+). We focus on the two main task relevant stimulus events during each session – image changes and image omissions, both forms of contextual novelty or surprise. Image changes occur after a variable number of repetitions of the same image, and are thus visually salient and task relevant, as potential opportunities for reward. Image omissions occur randomly (∼5% of non-change image presentations are omitted) during ophys sessions, but never during behavioral training; thus they are an unexpected break in an otherwise highly predictable stimulus cadence.

Accordingly, the analyses described below examine neural responses aligned to image changes or image omissions. We examine event aligned neural response timeseries, and compute several metrics of activity, by averaging the cell response in a 500ms window after event onset for each event, then averaging across all events within a session. Thus our analysis focuses on overall activity levels or preference of neurons for specific task events, rather than trial to trial variability or dynamics within a session.

Here we describe caveats and considerations for how physiology data was processed, nuances of the specific metrics that were used, and control analyses subdividing the dataset across cohorts, cortical areas, and depths.

#### Optical physiology data processing considerations

To account for the slow decay times of GCaMP6 indicators, we employed a method for detecting the onset time and magnitude of calcium transients (*28*). The magnitude of detected events was proportional to the increase in calcium fluorescence at each time point. The L0 event detection method makes assumptions about the calcium transient decay time and the relationship between spiking and measured calcium, which can vary across cell types. We estimated the decay time for each transgenic line using the MLspike autocalibration procedure (*29*), which resulted in a decay tau of 0.347 for Slc17a7-IRES2-Cre;CaMK2-tTa; Ai93(GCaMP6f) mice, 0.464 for Sst-IRES-Cre; Ai148(GCaMP6f) mice, and 0.52 for Vip-IRES-Cre; Ai148(GCaMP6f) mice. Still, event detection algorithms are not perfect and there are likely to be some false positives and false negatives present in the processed results. We chose parameters that were more likely to lead to false negatives than false positives. In addition, we noticed that the Vip inhibitory cell fluorescence traces sometimes had slow changes in activity that were not effectively modeled by a method assuming transients with rapid rise times and may have been missed. Accordingly, the calcium events used in our analysis are likely to correspond to bursts of activity rather than single spikes or slow fluctuations in calcium or spiking.

Our across session cell matching procedure relied on image registration of all pairs of sessions for a given field of view, followed by alignment of segmented ROIs after applying registration transforms to the ROI masks. Results of image registration were visually evaluated for all datasets and registration failures were either further optimized by hand tuning or removed from the dataset if the features of the image appeared visibly distinct from other sessions (i.e. the same cells were not present) and thus could not be accurately matched (typically due to differences in the z-depth of the imaging plane, which cannot be recovered algorithmically).

Given that the alignment was performed on ROI masks segmented independently in each session, failures of segmentation could contribute to failures of cross session cell matching. Our segmentation algorithm depended on cell activity (changes in fluorescence over time) for ROIs to be segmented; thus if a cell was physically present but not active in a given session, it would not be detected. We also applied a classifier based on hand annotated ground truth data to further filter ROIs as being cell soma or dendritic processes. We visually inspected the final set of ROIs for all sessions to screen for problematic cases (ex: many false positives), which were reprocessed with optimized parameters until results were satisfactory, or the session removed from the dataset if the signal to noise was too low for activity-based segmentation to work effectively.

Thus our segmentation and cell matching procedures erred on the side of being conservative to provide confidence in detected and matched ROIs, which may result in some cells with weak activity or out of plane fluorescence being excluded. More specifically, cells that were active on some days and not others may not have been identified by the segmentation algorithm on inactive days and thus would not be found to match across sessions. Accordingly, our analyses limited to cells that were matched across days are likely to be an underestimate the full population of neurons, particularly for cells with very strong selectivity for one session over another.

#### Image change response

##### Area and depth dependence of image change response

We quantified the average response in the 500ms following image changes for each cell, then compared across experience levels within each cell type for all recorded neurons in Familiar, Novel, and Novel+ sessions (session types as defined in fig. S4A). Image change responses were significantly larger during the Novel session for Excitatory and Vip populations, and significantly lower in the Sst population (fig. S7A).

To evaluate whether these results were influenced by the identity of the image set used during training, we quantified population average change responses for each cohort of mice, which had different 2-photon imaging configurations and image sets (described in fig. S2). The central result of novelty enhancement for excitatory and Vip cells, with novelty suppression in Sst cells, was present in nearly all subsets of the data, with some exceptions. In particular, results were not significant in some cases for Sst neurons, likely due to a low number of neurons in some subsets of the data (such as in cohort 2, with n=61 neurons across the 175 and 275um depths, or in cohort 3 at the 375um recording depth, with n=32 neurons).

When the average change response magnitude was split by cortical depth within each cell type (fig. S7A-D), we found a few key differences. The largest differences across experience levels were found in the middle imaging depths (175 and 275um depths), regardless of whether the difference was an enhancement (in excitatory and Vip cells) or a suppression (in Sst cells) of image change evoked activity (fig. S7D). A large novelty enhancement was also observed at the 75um depth for Vip cells. While the results for the 75um and 375um depths among excitatory cells were not significant, some interesting trends were observed. Image change responses were similarly elevated in the Novel and Novel+ sessions at the 75um depth (rather than being reduced in Novel+ as with other depths), indicating a slower timescale of familiarization. At the 375um depth, the trend was the opposite; image change responses were largest in the Familiar session and declined in the Novel and Novel+ sessions. These results indicate that experience influences change evoked activity in different ways across cortical depths, particularly for excitatory cells.

We also compared the fraction of image change responsive cells across cortical depths (fig. S7E). A cell was defined as change responsive if it had a significantly larger response during image changes compared to a shuffled distribution of activity from the gray screen period at the start of the session, for at least 10% of image change presentations. Among excitatory cells, a larger fraction of neurons were responsive to image changes in the 375um depth, compared to 75, 175, and 275um depths, however there was no difference across experience levels at the 375um depth. In contrast, all other depths showed a larger fraction of responsive cells in the Novel session compared to Familiar and Novel+. Among Sst cells, a larger fraction of cells was image change responsive during Familiar sessions compared to sessions with novel images for the 175 and 275um depths. Vip cells were more responsive to image changes during Novel sessions for the 175 and 275um depths. Notably, these middle depths have a larger number of cells, as they were imaged in both the multi-plane recordings (cohort 3) and the single-plane recordings (cohorts 1 & 2) (see fig. S4J for numbers of cells by depth).

Similar results were obtained based on the fraction of non-change image responses across depths and experience levels (fig. S7F). For instance, while excitatory neurons were more responsive overall for image changes (fig. S7E) compared to non-changes (fig. S7F), there were significantly more responsive cells during Novel sessions compared to Familiar and Novel+. However there were a few exceptions. The 75um imaging depth for Vip cells showed a significantly larger fraction of responsive cells for non-changes, but not for image changes. A significant difference in the fraction of responsive Sst neurons was observed across all imaging depths for non-change image responses, whereas only 175 and 275um depths were significant when considering change images. This could indicate that either enhancement or suppression of activity during image changes compared to repeated non-change image presentations could obscure overall novelty effects.

To quantify enhancement or suppression of single cell activity by image changes compared to repeated, non-change image presentations, we computed a change modulation index for each cell (fig. S7J-L). This index took the difference in the mean response during image changes and the mean response during repeated pre-change image presentations over the sum, giving a positive value for larger responses to changes and a negative value for larger responses to non-changes. Across the full population (fig. S7J), we found that excitatory and Vip cells were positively modulated by image change on average, and were further enhanced by stimulus novelty. In contrast, the distribution of the change modulation index for Sst neurons was centered around zero, indicating no enhancement or suppression by image changes on average. However the distributions were wide, indicating potential heterogeneity in change modulation in the Sst population. To investigate possible sources of this heterogeneity in change modulation, we compared distributions across cortical areas and depths, finding that some imaging depths showed experience dependent differences in change modulation while others did not. In addition, this comparison revealed that change enhancement was larger overall in the deepest imaging depth (375um) for Excitatory neurons, indicating a larger magnitude of enhancement by image changes in deeper layers that was independent of experience level.

Overall, our results quantifying various measures of image change responses at the population level demonstrated that overall image change response magnitude, and enhancement of changes relative to non-changes, were largest in deeper layers for excitatory cells, but were not experience dependent, while change responses were selectively enhanced by novelty in superficial layers. The results for Vip inhibitory cells largely mirrored the excitatory responses, albeit with an even stronger enhancement by novelty. Sst activity during image changes was suppressed during Novel sessions, with the largest differences occurring at 175 and 275um depths.

##### Response diversity in cells matched across sessions

As a direct comparison of the impact of stimulus novelty on single cell activity, we compared image change responses across each pair of sessions for cells that were identified and matched across days (fig. S7M,N). While the distributions were generally wide, indicating a variety of strengths and directions of novelty related changes, a few key trends were observed. Excitatory and Sst single cell responses were generally larger in Novel sessions compared to Familiar sessions, and in Novel+ sessions compared to Familiar sessions, however these differences were more pronounced in Vip cells, particularly when comparing Novel and Novel+ sessions, indicating a faster timescale of familiarization among Vip cells.

Consistent with this result, Vip responses were comparable in Familiar and Novel+ sessions, indicating a rapid return of activity to the familiar level in Vip cells. The distribution for excitatory neurons indicated that many cells’ activity remained elevated in the Novel+ session compared to Familiar sessions. The distribution of experience modulation for Sst cells indicated a broad suppression by novelty in all comparisons, however results were more mixed when comparing Familiar and Novel+ sessions, indicating that some cells had comparable responses for short and long timescales of familiarization, while others preferred one session type over another, potentially indicative of image feature tuning combined with experience dependence.

#### Image omission response

##### Area and depth dependence of image change response

We quantified omission related activity similarly to image change responses, by computing the average response in a window after omission onset (750 ms in this case) for each cell and averaging across cells for several key conditions (fig. S8A-F), as well as computing a modulation index comparing single cell activity during image omissions with activity during the gray screen period just prior to omissions (fig. S8I-J).

The largest and most consistent effect was among Vip inhibitory cells, which showed elevated omission related activity in Familiar and Novel+ sessions across all cohorts of mice (fig. S8A-C). When splitting by imaging depth, we found that the differences across experience levels were largest at the 175 and 275um imaging depths across the Vip population (fig. S8D,F,J,K), with omission signals particularly elevated in the Familiar session at these depths. In addition, Vip omission responses were larger overall in VISl compared to VISp during the Familiar session.

While Sst population activity was low overall during omissions (fig. S8A-F), and did not show strong experience dependent effects on average, when we compared activity before and after omission onset in single cells (fig. S8I,J), we found a smaller difference in the Novel session across all imaging depths, indicating a more similar activity level during the gray screen period, regardless of whether it occurred after an unexpected event or not. This is consistent with the population average (Fig. 3J), where a slightly larger ramp in activity is observed during the Familiar and Novel+ sessions, while the Novel session activity remains similar throughout the omission period. Still, these effects were present in a very small subset of cells (fig. S8K).

Excitatory population responses were larger during Familiar image omissions, particularly in the deeper imaging depths (275 and 375um; fig. S8D, F). The Novel+ session had slightly elevated omission related activity in the 175um depth. Interestingly, omission related activity was significantly higher in VISl compared to VISp, across all experience levels (fig. S8E).

We found that activity during the omission period was more similar to the pre-omission gray screen for the 75 and 375um depths (more positive omission modulation index in fig. S8J), indicating a smaller change in activity during omissions.

##### Response diversity in cells matched across sessions

When comparing the strength of omission responses across experience levels for single cells matched across days, we again observed that Vip cells had elevated omission related signals in Familiar and Novel+ sessions compared to Novel sessions, and that excitatory cells had larger omission responses during Familiar sessions (fig. S8L,M). When comparing Familiar and Novel+ sessions for Vip cells, we found a wider distribution, suggesting that some Vip cells may prefer Familiar sessions while others are more omission responsive during Novel+ sessions; an effect not observable at the population level (Fig. 3K), but confirmed when examining individual neurons (Fig. 3M)

##### Omission response across active and passive sessions and prediction of image identity

Omission related activity could represent an expectation violation, a prediction of upcoming image identity or reward, or simply be a release from contrast suppression during image presentations. As a control to test some of these options, we evaluated omission related activity in passive sessions, where mice viewed the stimulus in open loop with no availability of reward (fig. S9C). Prior to passive viewing sessions, mice received their daily allotment of water and thus were not thirsty during the imaging session, nor could they engage in the task even if they were motivated as the lick spout was retracted. Surprisingly, omission ramping across the Vip and excitatory populations were nearly identical during active behavior and passive viewing sessions with familiar images (fig. S9C), indicating that expectation of reward and task engagement are not central features driving image omission responses.

We also evaluated whether omission related activity served as a prediction of upcoming image identity by decoding the identity of the upcoming stimulus from population activity during omissions (fig. S9A,B). In this analysis, we again observed that omission related activity was more prevalent during Familiar and Novel+ sessions for Vip cells, and during Familiar sessions for excitatory cells, however it was not possible to classify image identity from omission related signals. The difference in omission response magnitude for Familiar and Novel sessions also did not depend on which image set was used during training (comparison of cohorts 1 & 2; fig. S8B,C), indicating that the particular features of the images used as Familiar or Novel also could not account for omission related activity differences with experience.

Overall, we find that omission signals are related to experience rather than task engagement, reward expectation, or image features.

### S5. Kernel regression models

#### Methodological considerations

We used a kernel regression model to quantify neural encoding of sensory, behavior, and task features, based on the example set in (*30*). Some kernels were aligned to events of interest with an empirically selected window around the event (ex: image presentations, hits, misses; fig. S10A). Others represented continuous features, such as running speed and pupil area, and were computed as a +/-1 second window around each time point in the session. Accordingly, the features included in our model occurred at different frequencies throughout the session. Continuous features spanned the entire session. Some event aligned features, such individual image presentations, comprised a large fraction of the session (1/8 of the full session for each of the 8 images), while others were more rare, such as hit and miss trials (of the ∼4000 individual image presentations in a session, ∼400 of them were image changes) and image omissions (5% of non-change image presentations). To account for these differences in feature frequency, coding scores were only computed as a fraction of the variance explained during the time steps when each feature was present.

##### Model evaluation

Accuracy of model fits varied across transgenic lines and session types (fig. S10B). For instance, Sst cells, which had the most reliable image evoked responses, had the highest explained variance on average. For excitatory and Vip cells, which both displayed higher responsiveness during the Novel session, model explained variance was also higher during the Novel session. These results were confirmed by correlating the signal to noise ratio of each cell’s trace (the standard deviation over the mean) to the explained variance in the model, showing a clear correlation for all cell types (fig. S10C). Thus, cells that were more active were better fit by the model. While there remains neural activity variance that is unexplained by the features used in our model or other methodological considerations, the overall sparseness and strength of activity in individual cells is a major factor in determining the ability of the model to explain neural variance.

Coding scores were computed for individual features or groups of features as the relative change in variance explained for the full model compared to reduced models produced by removing those features from the model and refitting (fig. S10D). Thus, a cell with a low overall explained variance could have a high coding score for a specific feature if all of the explained variance was due to that feature. Importantly, these reduced models are describing the unique explained variance of the dropped feature that cannot be explained by any of the remaining features. We also computed single feature models to look at the total contribution of each feature (fig. S10E) and found that coding scores were higher overall, indicating possible interaction among regressors and overlap in the variance explained across multiple features.

As a result, when evaluating changes in feature coding across different sessions for matched cells, the overall explained variance within each session must be taken into account to ensure a fair comparison across features in different sessions. For instance, a cell may have high variance explained in one session, with most variance explained by a feature such as images, resulting in a high image coding score, and have low explained variance in a different session, with most variance explained by a different feature such as image omissions, resulting in a high omission coding score. A direct comparison of these coding scores would suggest a similar encoding strength for both images and omissions, despite the cell having higher overall variance explained in the image coding session. Thus, when comparing single cell activity directly across sessions (Fig. 5; figs. S16-22) we normalized the coding scores in each session to the maximum variance explained across all sessions for each cell to facilitate a direct comparison of coding properties across days when quantifying the effect of experience on neural coding. p

#### Interpreting the results

Because we used a linear model without interaction terms, a cell would only be found to encode one of the continuous behavior variable if the cell activity trace linearly tracked that variable. Indeed, the running and pupil kernels (Fig. S13B,C) showed dynamics that were consistent with the average image locked running and pupil traces (fig. S5). Influences such as increases or decreases in running speed producing gain modulation of sensory evoked responses, which are known to occur in the mouse visual cortex, would not be captured by this model. Indeed, many cells that did not show behavioral encoding were found to have enhanced stimulus responses during running compared to stationary conditions (fig. S17H). Accordingly, our quantification of behavioral encoding represents a direct modulation of continuous cell activity by behavior variables, rather than a gain modulation or multiplicative effect.

Another factor to consider when interpreting the results is the relationship between the kernels estimated by the model and the coding score computed by dropping out specific features or set of features (figs. S11-13). The kernels indicate the influence of a given feature on cell activity relative to the start of the kernel window and are signed, with positive weight indicating an increase relative to the first time point of the window and a negative weight indicating a decrease. Coding scores do not have a sign, and a high encoding value for a given feature could be associated with an excitatory or inhibitory effect of that feature on cell activity. One example is for cells with strong image encoding; most cells with strong image encoding show enhanced activity following image onset, however some cells (Vip cells in particular) are suppressed following stimulus onset and had negative kernel weights during the stimulus window yet were still characterized as image encoding because that feature had a strong influence on cell activity (fig. S11A). Accordingly, coding scores should not be assumed to indicate a positive relationship between a given feature and cell activity, and only indicate that there was a relationship, regardless of its direction.

To reduce complexity in analysis and interpretation of the results, we considered coding for groups of features. In most cases, the pattern of change in coding across sessions for feature groups was consistent with the results for the individual features in that group (fig. S10D; fig. S14). For instance, changes in coding for hits and misses individually was fully captured by the pattern of task coding when both features were removed together. The experience dependent changes in behavior coding among Vip cells was the one exception – while the overall coding of all behavior features together did not show a significant change with novelty, we observed that encoding of licks was strongest during the Familiar session, and encoding of pupil was strongest in the Novel+ session, while running coding was highest during the first Novel session (fig. S14A). This indicates that encoding of specific behavior information in the mouse visual cortex depends on experience, with licks requiring extensive stimulus-action-reward association, and running and pupil influencing activity more during initial stimulus exposure.

##### Area and depth dependent coding properties

We found that pattern of experience dependent changes for individual features was typically consistent across cortical areas and depths, however the strength of encoding often differed (fig. S15). Area or depth specific differences in coding strength could be due to differences in overall activity levels, or to area or layer specific coding properties. For instance, all features were more strongly encoded in lower depths by excitatory cells (fig. S15A). Given the strong correlation between model accuracy (and thus coding score range and max) and overall responsiveness, and the known elevated firing rates among excitatory neurons in deeper layers of the visual cortex (*31*, *32*), this non-specific increase in coding strength may be associated with overall increased activity in lower layers. In contrast, in the Sst population, coding of behavior and task information were selectively elevated in lower depths, while image and omission coding remained the same (fig. S15A). This suggests that there are depth dependent coding properties among Sst neurons, as others have observed in the somatosensory cortex (*33*).

Encoding of behavior and omissions among excitatory neurons was found to be elevated in the higher visual area VISl, while image encoding was higher in VISp (fig. S15B). Similarly, Task encoding was elevated in VISl among Vip cells (fig. S15B). Interestingly, coding for omissions by the Vip population was higher in VISl during Familiar sessions, but was higher in VISp during Novel+ sessions, suggesting that signals related to unexpected omission of stimuli arise in different regions depending on the degree of familiarity with the stimuli before and after the omission (fig. S15B).

### S6. Functional Subtypes

The following section describes the rationale and methods for our clustering analysis, including justification for algorithmic choices, relevant controls, and elaboration on key results.

#### Across session normalization of coding scores

The analysis of coding scores in Figure 4 examined the strength of encoding of various features within each session, computing the coding score as the fractional change in explained variance when a given feature was removed from the model, relative to the overall explained variance of that cell in that session. The aim of our clustering analysis in Figure 5 was to identify patterns of changes in neuronal encoding across sessions with different experience levels (Familiar, Novel, Novel+), which required an additional normalization step.

To put feature encoding across sessions on the same scale and enable cross session comparisons, we normalized each cell’s coding scores to the maximum overall explained variance across sessions for that cell. For example, if a cell had low overall explained variance in a given session (session 1), and all of the explained variance in that session was accounted for by a single feature (feature 1), the un-normalized coding score for that feature in that session would be high. If the same cell had high overall explained variance in another session (session 2), the across session normalized coding score for feature 1 in session 1 would be computed relative to session 2’s overall variance explained. In this example, this procedure would result in a lower cross-session normalized coding score for feature 1 in session 1, relative to the coding scores across features in session 2, which explained a larger amount of variance for that cell. This ensures that features that account for a small amount of variance in one session are not considered as having a similar coding strength as features that encoded for a large amount of variance in another session, thus facilitating a more direct comparison of feature coding across sessions.

#### Clustering methods

We used a spectral clustering algorithm in combination with consensus clustering to identify similar patterns of across session normalized coding scores (fig. S16A). Spectral clustering was selected as the initial algorithm due to its robustness to noise and ability to capture global structure in the data.

Qualitatively similar results could be found using linear clustering methods, such as K-means, but the results of spectral clustering were more robust across iterations, which was part of our consensus clustering approach. Specifically, clustering was performed iteratively 150 times for a given value of K (number of clusters), and the probability of each pair of neurons being assigned to the same cluster across iterations was computed. The final clusters were selected by hierarchical agglomerative clustering on the co-clustering probability matrix (fig. S16A).

The optimal number of clusters was by comparing the within cluster variability for the original data compared to a shuffled dataset for different values of K (number of clusters) and selecting the value of K with the largest difference between original and shuffled data (i.e. the gap statistic method; fig. S16A). This procedure results in a selected number of clusters that most effectively describes the structure of across session normalized coding scores relative to the inherent variance in the data distribution, by identifying the number of clusters that minimizes within cluster variability. This procedure resulted in K = 14 clusters. We applied an additional threshold to the clusters that required that there were at least 5 cells of a given cell type within the cluster for it to be valid, and 2 small clusters that did not meet this criterion were excluded from further analysis.

Thus, our method resulted in 12 robust clusters that were consistent across clustering iterations and minimized within cluster variance. Computing the correlation across all pairs of cells within each cluster confirmed these results, demonstrating high correlation in the coding properties within each cluster (fig. S16C), with the exception of the non-coding cluster (cluster 1) which would be expected to be more variable given that the coding scores were near the noise level. We further verified that the clusters contained neurons from multiple mice, and that each mouse contributed to multiple clusters (fig. S16F) to ensure that coding properties represented by the clusters were general and consistently identified across subjects, and not a result of idiosyncratic properties of a given animal’s neural population or recording conditions.

Importantly, the clusters we identified varied in size (fig. S16B), indicating that some patterns of across-day feature encoding were more common than others.

#### Breakdown of clusters by cell type

Clustering was performed on all matched cells across the 3 cell types (fig. S17). Given that the number of excitatory cells was much larger than the number of inhibitory cells (3306 excitatory cells, 200 Sst cells, and 411 Vip cells), excitatory cells formed the largest fraction of each cluster. Accordingly, we computed the distribution of cells across clusters within each cell type (Fig. 5F) to assess the frequency of occurrence of particular coding properties within each cell population. Importantly, while excitatory and inhibitory cells sometimes co-clustered with each other (ex: clusters 2, 3, 4), in other cases they did not (ex: clusters 8 & 9), demonstrating that our methods were able to identify smaller clusters composed of just one inhibitory type, despite their smaller number overall. In addition, our clusters showed high within cluster correlation (fig. S16C), despite consisting of multiple cell types, indicating that patterns of coding properties could be very similar across cell types.

#### Cell ID shuffle control

Given the variability of neural activity from day to day and other potential confounds arising from recording stability or reliability of image segmentation, it is possible that some across session coding patterns could arise by chance. To further validate our clusters and evaluate how numerous cells with a particular coding pattern were relative to chance, we performed an additional shuffle analysis (fig. S18). While the gap statistic method used to select the optimal number of clusters broke all relationships between cell identity and coding features across days, in this analysis we shuffled cell identities within each cell type, while preserving the relationship of coding scores within each session (fig. S18A). We then ran our clustering procedure on the set of shuffled coding scores and evaluated the likelihood of identifying the same cluster properties as in the original, unshuffled dataset (fig. S18B). This allowed us to determine whether a particular pattern of across day changes in coding could have arisen randomly based on the distribution of feature coding within each session, or whether cell-specific relationships across experience levels were critical to obtaining the clusters in the original data, as well as to quantify how prevalent each coding pattern was in the original data compared to the shuffle.

On each iteration of clustering with the cell ID shuffled data, we computed the similarity (sum of squared error) between the shuffled clusters and the original data clusters to determine whether a given pattern of across day coding properties was present (fig. S18B,C). This was repeated over 500 iterations of clustering on the shuffled data to compute the probability of a given cluster appearing in the shuffled results (fig. S18D), and the size of each matched cluster relative to the original clusters (fig. S18E). Most of the image encoding clusters were routinely identified in the set of matched shuffled clusters, consistent with a large fraction of cells encoding images in the overall dataset (fig. S18D). However the shuffled clusters often differed in size from the original clusters, indicating that the original clusters could be more or less prevalent than expected based on the overall distribution of coding properties (fig. S18E). For example, selective encoding of familiar images (cluster 3) was more common in the original dataset compared to the shuffled clusters, indicating that this coding property is selectively enriched in the neural data, specifically for Excitatory and Sst cells. In contrast, selective encoding of images in the Novel+ session was less prevalent in the original data compared to shuffled clusters (fig. S18E), indicating that this pattern of experience dependent coding could have arisen randomly from the overall distribution of coding properties and session to session variability. That is not to say that cells within this cluster do not meaningfully encode images in this way, only to highlight that the exact identity of the neurons involved was not essential to finding that particular cluster.

#### Familiar-only sessions clustering control

As yet another means to evaluate the likelihood and prevalence of particular patterns of experience dependent encoding, we also performed clustering on familiar only sessions (fig. S19). This was enabled by our experimental design incorporating longitudinal imaging of the same populations of neurons over multiple repeated days with familiar or novel images (fig. S1F; fig. S4A). Specifically, we identified cells that were matched across 3 active behavior sessions with familiar images, and ran the same clustering procedure as was run on the data from Familiar, Novel, and Novel+ sessions. We again determined the optimal clusters using the gap statistic, to identify the number of clusters that minimized within cluster variance relative to a shuffled distribution (fig. S19C) and performed consensus clustering by identifying pairs of neurons with high probability of co-clustering across clustering iterations (fig. S19B).

Clustering on familiar-only sessions produced a slightly smaller number of clusters (10, fig. S19C) compared to clustering on sessions with novel stimuli (14 identified by gap statistic, 12 after removing very small clusters). However, the distribution of cells across the familiar-only clusters was very different from the results of clustering with novelty. Over 75% of the cells in the familiar-only dataset were contained within the first 3 clusters (clusters 1-3, 76.3% of all cells), whereas the 3 largest clusters in the novelty-containing dataset made up only 47.9% of the total population (fig. S19E,F). To get to >75% for the novelty-containing dataset required adding together 7 distinct clusters (clusters 1-7, 79.4% of all cells). This demonstrates that there is more heterogeneity in functional properties when novelty in incorporated in the dataset.

There were some similarities in the pattern of encoding in the familiar-only clusters and some of the clusters in the novelty dataset. For instance, the largest cluster in both cases was a non-coding cluster (43.5% of cells in familiar-only dataset, fig. S19D-G; 20.2% of cells in novelty dataset, Fig. 5; fig. S17). There was also a cluster encoding behavioral features across all sessions in both datasets (cluster 3 in familiar only dataset, 10.3% of cells, fig. S19; cluster 12 in novelty dataset, 5.5% of cells, Fig. 5; fig. S17), consistent with the finding that behavior encoding is not novelty dependent. The second largest cluster in the familiar-only dataset consistently encoded images across all 3 sessions (cluster 2, 20.7% of cells; fig. S19), a property not observed when clustering on sessions including novelty (Fig. 5; fig. S17), likely resulting in part from differences in cell tuning for image features in the familiar and novel image sets. In addition, there were several small clusters that encoded familiar images on one or two of the 3 familiar-only sessions (clusters 4-8, ranging from 2.1% to 6.2% of cells belonging to each; fig. S19). This is in contrast with the much larger fraction of cells belonging to clusters encoding familiar or novel images in specific session types in the novelty containing dataset; for example, the cluster encoding novelty on the first day of exposure (cluster 2, Fig. 5) contained 17.3% of cells, compared to clusters encoding familiar images only on a single day (clusters 4 and 6, fig. S19) which only contained 5 or 6% of cells. This suggests that the fraction of cells that could appear to have session specific coding due to drift, neural variability, or other experimental confounds, is around 5%.

While the overall sizes of the clusters in the familiar-only dataset differed from the novelty-containing dataset, the distribution of cells across clusters *within* each cell type (fig. S19F) was consistent with what could be expected based on familiar session coding properties in the novelty containing clusters (Fig. 5). For instance, most Sst cells encoded familiar images in the familiar-only clusters (69.3% of Sst cells, fig. S19E), which was also the predominant coding pattern among Sst cells in the novelty-based clustering (60.9% of cells between clusters 3, 8, and 7 which all encoded familiar images, Fig. 5; fig. S21). Similarly, Vip cells in the familiar-only dataset either encoded omission (cluster 9, 25% of Vip cells, Fig. 5; fig. S22), behavior (cluster 3, 17.3% of Vip cells), or none of the features (cluster 1, 57.7% of Vip cells) - the fraction of Vip cells encoding omissions and behavior during familiar sessions in the novelty-containing dataset were very similar (cluster 8, familiar omission coding, 21.3% of Vip cells; cluster 12, behavior coding, 19.1% of Vip cells; Fig. 5; fig. S22).

Overall, the clustering analysis performed on matched cells from 3 familiar-only sessions showed remarkable consistency in the fraction of cells within each cell type that encoded particular features in familiar sessions, but showed marked differences in the overall diversity and distribution of the clusters across the full population of neurons. This demonstrates that novelty is an important feature driving functional diversity within excitatory and inhibitory cell populations.

#### Experience dependent coding properties and activity dynamics within and across cell types

There are a few notable characteristics of specific clusters, and specific clusters within each cell type, that should be considered. While we show the average coding properties of each cluster across cell types in Figure 5, and note the high within cluster correlation of coding scores within each cluster, examining the change and omission aligned traces for each cluster split by cell type reveals some intriguing differences. First, it is important to note that the average image and omission evoked responses shown in Figure 5 will inevitably reflect the cell type with the largest number of cells within each cluster, which is invariably the excitatory cells, except in the cases of the omission clusters (clusters 9 & 10) which are dominated by Vip cells. Accordingly, the image, change, and omission aligned responses for each cluster within excitatory (fig. S20), Sst (fig. S21), and Vip (fig. S22) cells should be considered, and we will compare them directly below.

First, a few notable characteristics of image change evoked responses across cell types (fig. S20-22E): 1) within excitatory cells, all image coding clusters show some degree of enhancement by image changes, 2) image change modulation is variable across Sst clusters, with some showing change enhancement (cluster 10), change suppression (cluster 4), or experience dependent change modulation (cluster 6, suppressed by image changes in Novel+, no difference between change and pre-change in Familiar session), 3) similar to the excitatory cells, Vip clusters that encode images show enhancement by image changes (although, notably, Vip cells only respond to images during the first novel session).

Next we consider the behavior encoding cluster (cluster 12), which is shared by all 3 cell types (4-5% of excitatory and Sst cells, 19% of Vip cells) – the image aligned response profile of this cluster shows elevated activity during the inter-stimulus interval, and suppression by image onset (Fig. 5D; fig. S20-22D,E). For excitatory and Sst cells, this elevated interstimulus response remains elevated at a consistent level during image omissions and is consistent across all experience levels, whereas Vip behavior coding cells have a ramping profile that continues to increase during omissions (similar to omission coding cells) and is specific to Familiar and Novel+ sessions (instead responding to novel images in the Novel session). These observations are consistent with a linear relationship between calcium fluorescence in Sst and Excitatory cells and running and/or pupil diameter, which fluctuate at the stimulus frequency and decrease just before or just after stimulus onset (on average; fig. S5). The Vip response profile for the behavior coding cluster appears to be a mixture of linkage to running and pupil, along with some degree of omission response in Familiar sessions and image response in Novel sessions (i.e. multiplexed coding of multiple features).

The two task encoding clusters (clusters 10 and 11) had rather distinct coding characteristics and response profiles. Enhanced task coding during the Novel session (cluster 11) by excitatory and Vip cells was highly specific to image and task features, with a response profile that was essentially silent except during the image change (Fig. 5D,E; fig. S20D-F; fig. S22D-F). Enhanced task coding during Familiar sessions (cluster 10) was accompanied by broad encoding of images across all sessions and cells belonging to this cluster had distinct response dynamics across cell types. Sst cells with Familiar task encoding (cluster 10; fig. S21) responded similarly to the Novel task coding cluster – with an enhanced response to image changes relative to repeated stimuli in an experience dependent manner (specific to the Familiar session; note that this is the only Sst cluster with change enhancement). Vip cells with Familiar task encoding (cluster 10; fig. S22) showed dynamics more similar to the behavior coding cluster (cluster 12) with elevated activity during the inter-stimulus interval and suppression by stimulus onset. Excitatory cells with Familiar task encoding showed a blend of the two – elevated activity during the interstimulus interval that was suppressed by onset of repeated stimuli but enhanced by image changes (cluster 10; fig. S20). This could be suggestive of a shared source of input to Sst and excitatory cells, and a distinct source to Vip and the same excitatory cells.

The omission coding clusters primarily consisted of Vip cells, but a small fraction (<2%) of excitatory neurons also encoded omissions. Both cell types responded to omissions with a ramp-like increase in activity following stimulus omissions, and during the inter-stimulus interval (clusters 8 & 9; fig. S20, fig. S22). Excitatory omission coding cells showed ramping behavior during novel sessions, while Vip omission coding cells switched to encoding images during novel sessions. This could arise from a shared source of input, or local excitatory drive onto Vip cells, from both omission and novel image coding excitatory neurons.

It is also important to note that, while both omission and behavior coding clusters show inter-stimulus and omission related activity, the dynamics are distinct – with the omission coding response being more consistent with an anticipatory ramping pattern (or a very delayed release from suppression that has not reached its saturation point) and the behavior coding response profile being more consistent with alignment to behavioral dynamics (fig. S5) or potentially a high baseline firing rate that is suppressed by stimuli.

Among image coding clusters, the familiar image coding clusters stand out as having unique dynamics (Fig. 5, figs. S20-22). For instance, the cluster selective for Familiar sessions (cluster 4), which is comprised of excitatory and Sst neurons, shows elevated baseline activity specific to the Familiar session. This effect is more pronounced in excitatory than Sst cells, and is consistently elevated throughout the inter-stimulus interval; whereas the Sst response appears to increase just prior to stimulus onset (cluster 4, fig. S20D-E, fig. S21D-E). It is possible that this elevated baseline activity among excitatory neurons coding for Familiar stimuli could arise from Vip mediated disinhibition, provided by Vip cells with ramping activity during omissions and the inter-stimulus interval (clusters 8 or 12). These Vip cells could suppress Sst cells that do not encode Familiar images, forming a specialized subnetwork that could serve to prime Familiar coding excitatory cells to respond to upcoming image changes. Alternatively, a cell type not imaged in this study, such as parvalbumin (PV) expressing inhibitory neurons could be involved. In any case, this intriguing observation warrants further inquiry.

Finally, the Sst image coding cluster with a novelty suppressed response profile (cluster 8, fig. S21) stands out by displaying an experience dependent shift in response latency, with the response to highly familiar images in the Familiar session occurring earlier than the response to recently familiar images in the Novel+ session. In addition, this cluster shows a reduced change response in the Novel+ session, but similar response to repeated and change images in the Familiar session. In both cases, these shifts in dynamics are indicative of longer timescale plastic changes, such as changes in excitability, arising from distinct neuromodulatory inputs, experience dependent changes in expression of ion channels, or circuit connectivity and synaptic dynamics.

#### Distribution of experience dependent coding clusters across cortical areas and depths

As our recordings spanned multiple cortical depths (from 75 – 375 um from the cortical surface, approximately upper layer 2 through upper layer 5; see section “S1 - Relationship of imaging depth and cortical layer”) and visual areas (VISp, also called V1, and VISl, also called LM), we evaluated for biases in cluster distributions across areas and depths.

We quantified the proportion of cells belonging to each area or imaging depth for each cluster, relative to the overall proportion of cells in that cluster, then tested for significant differences in the proportion of cells across areas or depths within each cluster (figs. S20-22G,H). For instance, if 20% of all Vip cells belong to cluster 12, and 30% of all cells in VISl fall into cluster 12, but only 10% of VISp cells are in cluster 12, that would indicate that the coding properties associated with cluster 12 are more prevalent in VISl compared to VISp While most of the Sst and Vip inhibitory clusters were relatively small (typically fewer than 50 cells per cluster), splitting further by depth or area often did not yield sufficient numbers to draw meaningful conclusions about area and depth biases (figs. S21, S22). However, there were a few exceptions. For instance, we found that the Vip clusters encoding behavioral (cluster 12) and task (cluster 10) features were biased to VISl compared to VISp. Sst cells showed a similar, but nonsignificant, trend for behavioral coding biased to VISl, however the task coding cluster (cluster 10, selective for Familiar sessions) showed the opposite bias, and was more prevalent in VISp. There were no significant depth differences among Sst clusters. Interestingly, while few Vip cells were found in the deeper layers (275 and 375 um imaging depths), there was an over-representation of cells encoding omission of Familiar stimuli (cluster 8), along with behavior (cluster 12) and non-coding cells (cluster 1). Vip cells were more numerous in the superficial layers (75 and 175 um imaging depths) and were also more diverse, being split between multiple clusters without any significant bias (with the exception of the non-coding response profile, which was prevalent in the 75um depth).

The distribution of excitatory neurons across areas and depths benefitted from a larger sample size (fig. S20G,H). We found that task encoding (clusters 10 and 11) by excitatory cells was more prevalent in VISp compared to VISl, similar to Sst cells. Similarly, behavior coding (cluster 12) was biased to VISl, similar to Vip cells. This could suggest that inhibitory neurons within a given area reflect the coding properties of local excitatory neurons, or vice versa. Consistent with this notion, task encoding by excitatory cells (cluster 10) was heavily biased to deeper layers (375 um depth), similar to both Vip and Sst cells. We also found an interesting tradeoff across cortical depth among the image encoding excitatory clusters – the excitatory cluster encoding novel images (cluster 2) was biased to the middle depths (175 and 275 um imaging depths), while excitatory clusters encoding familiar images (clusters 4 and 6) were significantly biased to the lowest imaging depth (375 um).

## Materials and Methods

### Mice

All experiments and procedures were performed in accordance with protocols approved by the Allen Institute Animal Care and Use Committee. Male and female transgenic mice expressing GCaMP6 in various Cre-defined cell populations were used in these experiments (*1*). The three genotypes used in this study were *Slc17a7*: Slc17a7-IRES2-Cre;Camk2a-tTA;Ai93(TITL-GCaMP6f), n=41; *Sst*: Sst-IRES-Cre;Ai148(TIT2L-GC6f-ICL-tTA2), n=19; *Vip*: Vip-IRES-Cre;Ai148(TIT2L-GC6f-ICL-tTA2), n=22. Prior to surgery mice were singly-housed and maintained on a reverse 12-hour light cycle (off at 9am, on at 9pm); all experiments were performed during the dark cycle.

### Surgery

All mice received a headpost and cranial window surgery as previously described (*2*, *3*). Briefly, surgery was performed on healthy mice that ranged in age from 5-12 weeks. Mice were deeply anesthetized with isoflurane prior to removing skin and exposing the skull. A custom titanium headframe was cemented to the skull and a circular piece of skull 5 mm in diameter was removed, durotomy performed, and a glass coverslip stack was cemented in place. Mice were given 2 weeks to recover from surgery before intrinsic signal imaging.

### Intrinsic Signal Imaging

Intrinsic signal imaging (ISI) was used to measure the hemodynamic response of the cortex to visual stimulation across the entire field of view. As previously described, ISI was used to delineate functionally defined visual area boundaries for targeting of 2-photon imaging experiments (*3*).

#### Data Acquisition

Mice were lightly anesthetized before imaging sessions began with a vasculature image acquired under green light. Next the imaging plane was defocused and the hemodynamic response to a visual stimulus was imaged under red light. The stimulus consisted of an alternating checkerboard pattern (20° wide bar, 25° square size) moving across a mean luminance gray background. On each trial, the stimulus bar was swept across the four cardinal axes 10 times in each direction at a rate of 0.1 Hz(*4*).

#### Data Processing

A minimum of three trials were averaged to produce altitude and azimuth phase maps, calculated from the discrete Fourier transform of each pixel. A “sign map” was produced from the phase maps by taking the sine of the angle between the altitude and azimuth map gradients. In the sign maps, each cortical visual area appears as a contiguous red or blue region (*5*).

To provide a reliable map for subsequent targeting of 2-photon calcium imaging experiments, a consistent anatomical coordinate corresponding to the center of VISp (which maps to center of the retina) was used to realign the maps. A map of eccentricity from the VISp centroid was produced by shifting the origin of the map of visual eccentricity to the coordinates at the VISp centroid, thereby representing the retinotopic gradients relative to this point. A representation of the corresponding retinotopic location is present in nearly all higher visual areas (HVAs). Using these modified VISp and HVA targets for optical physiology experiments ensured that recorded neurons represent a consistent region on the retina, approximately at the center of the right visual hemifield.

### Behavior Training

#### Water restriction and habituation

Throughout training mice were water-restricted to motivate learning and performance of the behavioral task (*6*). Mice had access to water only during behavioral training sessions or when provided by a technician on non-training days. During the first week of water restriction mice were habituated to daily handling and increasing durations of head fixation in the behavior enclosure over a five-day period. The first day of behavior training began after 10 days of water restriction. Mice were trained 5 days per week (Monday-Friday) and were allowed to earn unlimited water during the daily 1-hour sessions; supplements were provided in a home cage water dish if the earned volume fell below 1.0mL and/or body weight fell under 80-85% of initial baseline weight. On non-training days mice were weighed and received water provision to reach their target weight, but never less than 1.0 mL per day.

#### Apparatus

Mice were trained in custom-designed, sound-attenuating behavior enclosures equipped with a 24” gamma-corrected LCD monitor (ASUS, #PA248Q). Mice were head-fixed on a behavior stage with 6.5” running wheel tilted upwards by 10-15 degrees. The center of the visual monitor was placed 15 cm from the eye and visual stimuli were spherically warped to account for the variable distance from the eye toward the periphery of the monitor. Water rewards were delivered using a solenoid (NI Research, #161K011) to deliver a calibrated volume of fluid through a blunted, 17g hypodermic needle (Hamilton) positioned approximately 2-3 mm away from the animal’s mouth using a custom-made 3-axis motorized stage.

### Change detection task

#### Overview

The change detection task and automated procedure for training this task have previously been described in detail (*7*, *8*). Briefly, mice were trained using a behavioral program implementing a go/no-go change detection task (Fig. 2A; fig. S3A,C,D). Mice were presented with a continuous stream of flashed visual stimuli (250ms stimuli interleaved with 500ms gray screen) and were trained to lick a reward spout when the identity of the stimulus changed. If mice responded correctly within a short, post-change response window (150-750ms after stimulus change) a water reward was delivered. A ‘grace period’ of 3 seconds occurred after each change, during which no additional image changes could occur, thereby providing time for reward consumption before the next trial was initiated. At the start of each trial, trial type was determined (87.5% “GO” or 12.5 % “CATCH”) and a change time was drawn from a geometric distribution ranging from 4 to 12 flashes (3 to 9 seconds; fig. S3C,D). If the mouse licked prior to the stimulus change the trial was reset (“aborted”) up to 5 times before a new trial and change time was re-drawn. Thus, continuous licking would result in no opportunity to obtain reward. No other punishment was delivered for false alarms or aberrant licking.

#### Automated Training

Mice were trained using an automated training procedure that consisted of 4 stages of increasing complexity (fig. S3F). On Day 1 of the automated training protocol mice received a short, 15-min “open loop” session during which non-contingent water rewards were delivered coincident with 90° changes in orientation of a full-field, static square-wave grating (Stage 0). This session was intended to 1) introduce the mouse to the fluid delivery system and, 2) provide the technician an opportunity to identify the optimal lick spout position for each mouse. Each session thereafter was run in “closed loop”, and progressed through 3 phases of the operant task: Stage 1: static, full-field square wave gratings (oriented at 0° and 90°, with the black/white transition always centered on the screen and the phase chosen randomly on every trial), Stage 2: flashed, full-field square-wave gratings (0° and 90°, with phase as described in 1), and Stage 3: flashed full-field natural scenes (8 “Familiar” natural images). In a subset of mice the image sets labeled “Familiar” and “Novel” were switched in order to balance the experimental design.

#### Progression through training stages

Starting with Stage 1, the advancement criteria required mice to achieve a session maximum performance of at least d-prime=2 (calculated over a rolling 100 trial window without trial count correction) during two of the last 3 sessions. The fastest progression from Stage 1 to Stage 3 was 4 training days. Once mice exhibited consistent performance (d-prime >/= 1 over 3 consecutive sessions) they became eligible to transition to the 2-photon calcium imaging stage of the experiment.

### 2-photon Calcium Imaging

Calcium imaging data was acquired using two microscope platforms, each of which was built around our custom-designed Allen Brain Observatory behavior platform and common mouse-to-screen geometry as previously described (*3*, *9*, *10*).

#### Single Plane Imaging Apparatus

Single-plane calcium imaging was performed using a 2-photon microscope (Scientifica Vivoscope), as used by de Vries et al., 2020 and Garrett et al., 2019. Scientifica microscope design is based on 8 kHz resonant scanning mirror and employs conventional hardware (photomultiplier tubes, Hamamatsu; transimpedance amplifier, Femto; DAQ hardware, National Instruments; 16x imaging objective, Nikon) to collect emitted fluorescence and form an image on the acquisition computer. The microscope is controlled by the company’s proprietary LABView software SciScan. Laser excitation was provided by a Ti:Sapphire laser (Chameleon Vision, Coherent) at 910 nm. Pulse dispersion compensation was set at ∼10,000 fs^2^. Movies were recorded at 30Hz using resonant scanners over a 400 µm field of view.

#### Multi-Plane Imaging Apparatus

Multi-plane calcium imaging was performed using a Dual-Beam Mesoscope (Multiscope) that allowed us to double imaging throughput (*11*). The second laser beam was introduced to the original 2P-RAM system (*12*), packaged into a compact opto-mechanical add-on unit, and optimized for ease of alignment. The dual-beam modification consisted of a 1) delay line used to split the original laser beam into 2 and delay one of the beams by half a period of the excitation laser; 2) a secondary z-scanner and 3) custom-built demultiplexing unit. The delay line allowed for temporal encoding of the excitation beam and further demultiplexing of the detected fluorescence based on the arrival time at the photodetector. A secondary z-scanner allowed us to send two beams to the two focal planes located along Z axis. Laser excitation was provided by a Ti:Sapphire ultrafast laser (Chameleon Ultra II, Coherent). Pulse dispersion compensation was optimized for GCaMP6 using a custom-built external pulse compensation module based on a single-prism four-path design (*12*, *13*). The Multiscope was controlled with customized ScanImage software (VidrioTech) as well as an in-house developed Workflow Sequencing Engine. Like conventional 2-photon microscopes, the emitted fluorescence was detected using a single photomultiplier tube, and a custom analog demultiplexing circuit was used to separate fluorescence from two planes. This was achieved by multiplying the PMT signal with two complimentary square waveforms, where each waveform corresponded to the temporal window during which fluorescence received by the PMT consisted of the signal from one of the focal planes. The duration of the integration window was 6.25 ns (half a period of the excitation laser’s pulse train), which is not enough to fully capture the decay of fluorescence, which resulted in the tail of fluorescence leaking to the opposite integration window and causing inter-plane crosstalk (∼10% remaining crosstalk on average). We used an ICA-based demixing algorithm to further clean up the data acquired in simultaneously imaged focal planes. National Instruments data acquisition hardware (PXI chassis, PXIe6363 DAQ boards) was used to control the microscope, form, and record the image.

#### Data Acquisition

Daily preparations for the 2-photon imaging experiments were conducted under ambient red light to maintain the reversed day-night cycle, and imaging itself was performed in the dark. Mice were head-fixed in a behavior stage identical to that used during behavior training. A water immersion objective was used for single-plane experiments on the Scientifica microscope whereas a water-based ultrasonic gel was used as immersion medium for the multi-plane experiments on the Multiscope. Two-photon movies (512x512 pixels, 31 Hz for single plane and 512x512 pixels, 11 Hz for each plane in multi-plane experiments), eye tracking (30 Hz), and behavior (30 Hz) were recorded and continuously monitored.

Recording sessions were ∼1 hour long, but could be interrupted if any of the following was observed: 1) mouse stress as shown by excessive secretion around the eye, nose bulge, and/or abnormal posture; 2) excessive pixel saturation (>1000 pixels) as reported in a continuously updated histogram; 3) loss of baseline intensity caused by bleaching and/or loss of immersion water in excess of 20%; 4) hardware failures causing a loss of data integrity. At the end of each experimental session, a z-stack of images (+/- 30 µm around imaging site, 0.75 µm step) was collected to evaluate cortical anatomy as well as z-motion during acquisition. In addition, a full-depth cortical z stack (∼800 µm total depth, 5 µm step) was collected to document the imaging site location.

#### Quality Control

Each experimental session was analyzed for data integrity based on a broad range of operational parameters. A comprehensive report was automatically generated to track data trends, animal behavior, experimental failures and errors. To minimize bias, QC reports were reviewed by rig operators other than the operator who performed the session.

Assessment of the following quality metrics was performed after each imaging session. Failure to meet any of these criteria resulted in the session being retaken on a subsequent day.

1. Image Saturation: Initial movie frames were assessed to confirm that photonic saturation did not exceed 1000 pixels and the full dynamic range of the recording system was adequately covered.
2. Photobleaching: Epochs of fluorescence at the beginning and end of a session were compared to ensure that baseline fluorescence did not drop greater than 20%.
3. Field of View Targeting Validation: Targeted imaging locations were checked against the intrinsic signal imaging data, using a registered coordinate system, to confirm that data was collected from the correct visual area.
4. Z-Axis Stability: Stability of image recording was assessed by comparing a windowed average image from the first and last 5 minutes of the experiment to a z-stack of images (+/- 30 μm around imaging site, 0.75 μm step) collected at the end of each experimental session to calculate the amount of drift that occurred over the session. Experiments with z-drift above 10μm over the course of the entire session were excluded.
5. Animal Stress: Behavior videos were viewed to confirm that animals did not show excessive signs of stress. Any animal that showed eye secretion covering the pupil or excessive orbital tightening was returned to its home cage to recover. The presence of nose bulge, flailing and abnormal postures was also monitored.
6. Temporal Sync: Temporal alignment of data streams was confirmed.
7. Hardware/Software Failure: Multiple datastreams and metrics were assessed to ensure that incoming data integrity was not compromised by hardware and/or software related errors
8. Excessive Motion: Imaging frames were checked for residual motion (after motion correction algorithms had been applied)
9. Interictal Events: Presence of interictal events was assessed by measuring the full field fluorescence and calculating the intensity spike prominence and width for the first 10,000 frames of the 2-photon imaging movie. Experiments with a non-zero probability of interictal events were then checked manually in order to exclude any potentially epileptic mice (*14*).

A final, container-level QC assessment was performed once all data collection for a mouse was completed. This secondary assessment included assessing the following metrics:

1. Full Container Status: Containers (the set of imaging sessions for a given field of view) were confirmed to contain all required datasets.
2. Brain Health: 2-photon serial tomography sections were examined to assess general brain health. Health assessment includes checking for excessive bruising, brain abnormalities, deformities, necrotic tissue damage, and checking for any signs of laser damage.
3. Cell Matching: Imaging fields of view were checked across all experiments in a container to confirm successful targeting of the same field of view. This assessment was made visually, based on similarity in patterns of vasculature and the presence of clearly identifiable matched cells across sessions. A final visual check was conducted after data processing using red-green overlay images after cross-session image registration had been performed, along with a computed value of the <u>structural similiarity index</u> as a confirmation of image similarity.

### 2-Photon Data Processing

An overview of the data processing steps for both single- and multi-plane imaging is shown below.

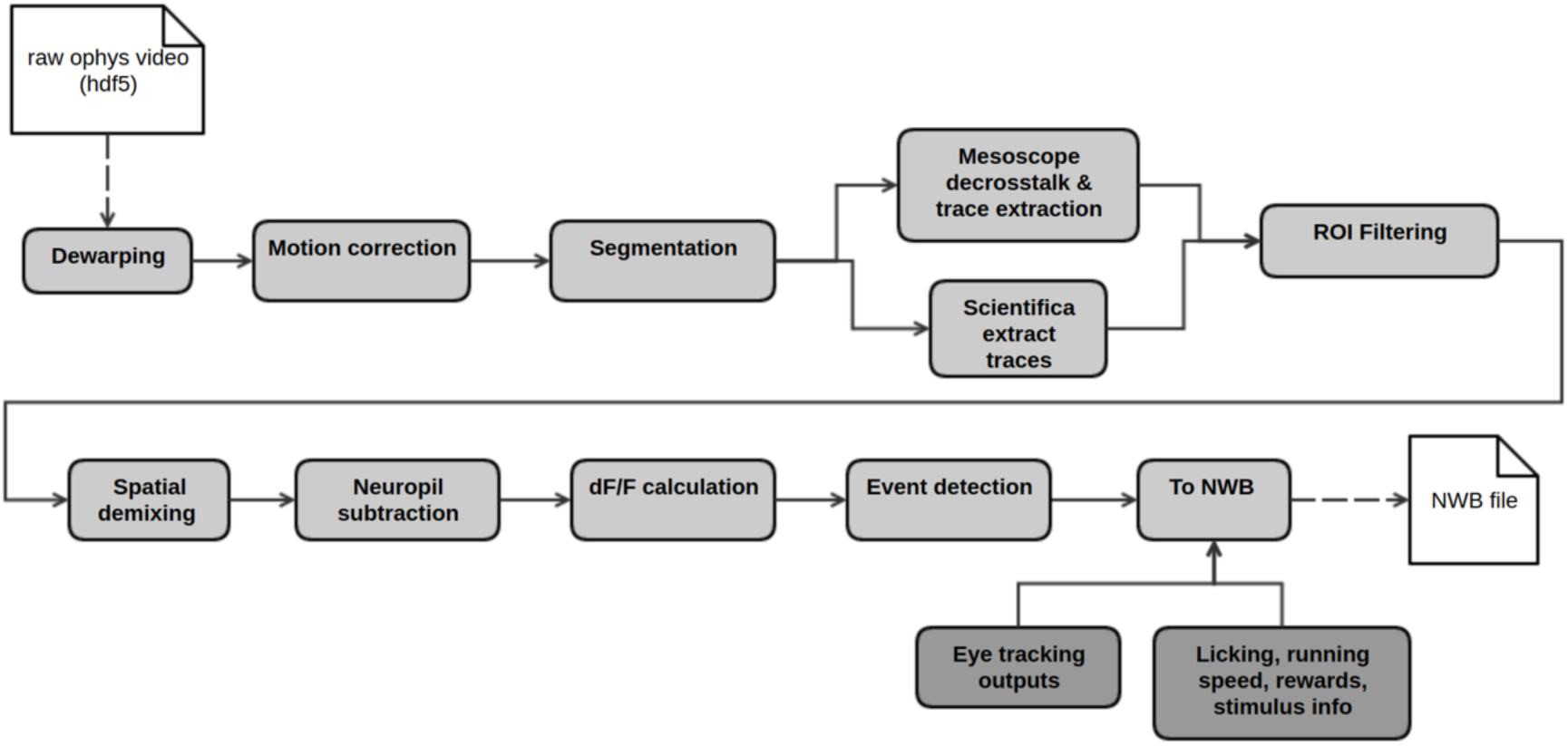

#### Dewarping

To account for variation in scanning rate across each line in the 2-photon imaging frame, due to the non-constant speed of the resonant scanner, data acquired with the Scientifica microscopes required a dewarping step (Multiscope instruments have built-in dewarping correction).

Correcting for the warping in the image involves taking a subset of the warped image’s columns, towards the edges of the image where the scanning speed is lower, and combining them using parameters derived from calibration data acquired with a standard grid image. To determine correct dewarping parameters, the grid image is adjusted until the dewarped grid is uniform. Each side of the image is dewarped independently.

The following equations are used to determine which columns are chosen from the warped image and how they are combined. The *j*^th^ column of the dewarped image will be given by the formula:

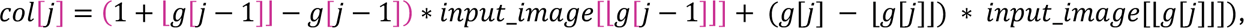

where

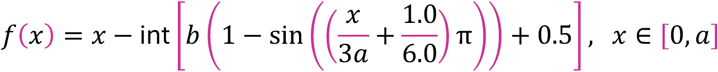

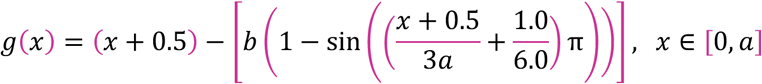

Notice that this is taking a linear combination of the two (generally, adjacent) columns of the original image, using the decimal parts of the g function as coefficients. If these ever end up being non-adjacent columns from the original image, then we also add the value of the skipped column to col[j].

In the event that both f[j] and g[j - 1] are not nonnegative, then we fill that column with the average from the original image. And if f is nonnegative but g is not, then we simply replace that column with the exact same column from the original image.

For each experiment, there are four parameters that tell us how much the image has been warped. These are called *aL*, *aR*, *bL*, and *bR*. The L and R refer to the side of the image which is being dewarped. In the equations above, a and b are replaced with their corresponding parameter, depending on whether you are dewarping the left side or the right side. The parameters *aL* and *aR* specify how far (in pixels) into the image the warping occurs. The parameters *bL* and *bR* are more a measure of how warped the image is in those areas.

#### Motion Correction

We used <u>Suite2P</u> v0.9.3 <u>rigid registration</u> for motion correction of 2-photon movies. Suite2P performs an iterative phase-correlation-based registration on a small subset of frames to generate a <u>reference image</u> from the average projection of those frames. The registration of the entire movie then proceeds with registration by phase correlation of each frame to this reference image. We saved Suite2P’s registered output tiff stacks and concatenated them into a single hdf5 format compatible with the rest of our processing pipeline.

Suite2P’s parameter *maxregshift* has a default value of 0.1, which clips lateral motions of more than 10% of the FOV dimension. We found a few examples in our release data where our experiments had real, long-timescale lateral shifts of greater than 10%. We increased the *maxregshift* parameter to 0.2, to allow for registration of these experiments. A consequence of this change was that up to 20% lateral shifts were allowed by Suite2P over short timescales as well. We observed that these short timescale shifts, often single frames, were likely not physical shifts, but struggles of the registration algorithm to register low signal-to-noise frames. Across an entire experiment, we monitor the worst-case shifts and establish a motion border which invalidates any ROI which touches it. To prevent too much inflation of the motion border exclusion area by these unphysical short timescale shifts, we detrended the x and y lateral corrections with a median filter over a 3 second window. We clip frame displacements to a +/-5% window around any outlier more than 5% above or below the detrended corrections was truncated to 5%. This allowed us to handle physical long timescale drifts and limit the impact of artifactual short timescale shifts. Any movie frames where the lateral shift was truncated were translated to the new truncated translation value.

For each movie, we produce the following outputs: the registered hdf5 movie for downstream processing, the motion correction shifts both from Suite2P directly and after our detrended clipping step, and Suite2P’s output corrXY, the value of phase-correlation at the best alignment for each frame. For QC inspection, we generate plots showing the x and y lateral shifts and corrXY resulting from motion correction, as well as a video preview showing the movie before and after motion correction, side-by-side, averaged to 0.5 frames-per-second and a 10x playback speed.

#### Cell Segmentation & ROI Filtering

The Visual Behavior 2P project used the same segmentation procedure that was developed for the Visual Coding 2P dataset, published in de Vries et al., 2020. The active cell segmentation module was designed to locate active cells within a field-of-view (FOV) by isolating cellular objects using the spatial and temporal information from the entire movie. The goal of the active cell segmentation module is to achieve robust performance across experimental conditions with no or little adjustment, such as different mouse cell lines, fluorescent proteins (e.g., GCaMP6f or GCaMP6s), and FOV locations of visual areas and depths. The process begins with the full image sequence as input to apply both the spatial as well as temporal information to isolate an individual active cell of interest without data reduction, such as by PCA, and does not make assumptions about the number of independent components existing in the active cell movie. Also, in contrast to other methods, this approach separates the individual steps, including identifying and isolating each cellular object, computing confidence of each identified object (by object classification) and the step of resolving objects overlapping in x-y space (which lead to cross talk in traces), so that each can be improved upon if necessary.

#### Pre-segmentation

The motion corrected image sequence was spatially median filtered (using 3x3 pixel kernel) to reduce white noise. The sequence was then low pass filtered and downsampled by 1/8 temporally to enhance the signal-to-noise ratio (SNR). The processed image sequence was then divided into periods of fixed temporal length p, where p = 50 frames (∼13.3 sec.). The maximum projection image from each period and the mean image (mu_image) of the whole sequence were computed. The maximum projection image from all temporal periods, called Periodical Projection frames PP(t), were further normalized to become Normalized Periodical Projection (NPP) frames:

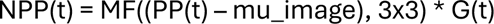

Where G(t) is the frame intensity normalization gain computed based on the intensity histogram of each PP(t). This is to normalize any change in overall intensity across the experiment, and to reduce experiment-to-experiment variability. MF(3x3) is median filtering with a 3x3 pixel kernel.

Note in each NPP(t), a subset of cells can be found with changes in fluorescence during that time period. With sufficient experiment length, and with many sweeps of different stimuli, various repetitive cell firing patterns can be found. Cells with overlapping spatial positions in x and y can be observed as firing at different time frames, allowing the following detection process to identify them individually despite having spatial overlap.

#### ROI Detection

Adaptive and mathematic morphological image processing techniques were applied to process each NPP(t). After band-pass filtering, an initial binary object map was generated by thresholding the resulting image minus a low pass version of itself to capture spatially varying background intensity.

Conditional dilation, erosion and connected component analysis were applied to filter the candidate binary objects and fill holes. The final set of regions of interest (ROIs) in each NPP were identified using another connected component labelling and a simple rule-based classifier. This classification was based on comparing measured morphometric attributes (object area, shape, intensity, uniformity, etc.) of the objects to the statistics derived from the targeted active cell components in the sample data sets. After each frame was processed, a set of candidate ROIs from each NPP were then grouped with candidate ROIs from all other NPPs. ROIs within a similar spatial location (defined by the distance between centroids < 5 µm) and with similar morphometric attributes (e.g., delta (area or shape) < 20%) across NPP frames were grouped as the same cell object. The ROI with the highest contrast and with shape and area within range statistically derived from sample data was selected to represent that cell in the mask image and in a composite image for visual QC. ROIs with different spatial locations (centroids > 5 µm apart) and or dissimilar morphometric attributes are recorded as different cells.

Occasionally two or more spatially overlapping ROIs could be found that were active in the same timeframe and were therefore detected as a single ROI for that NPP. Additional steps to classify them as multiple-cell objects were taken using their attributes of combined area and shape (eccentricity).

After ROI detection in each NPP frame and grouping of all ROIs across frames was completed, a set of “unique” active cell objects was identified. Cells near the FOV boundaries (3 µm) were eliminated from further consideration due to the fact that motion shifts can create boundary effects to the traces computed from these cells. To generate the segmentation mask image, all non-overlapping cells were placed in a single mask plane and overlapping cells were placed in subsequent planes, to ensure unique identification of all cells.

#### Crosstalk Removal in Multiscope Data

Crosstalk between focal planes is a fundamental limitation in multiplexed microscopy systems, such as the Multiscope. Details about system hardware and methods used to optimize crosstalk in the data are described above and can be found in Orlova et al., 2020 (***11***). Crosstalk removal was performed on fluorescence traces using an ICA-based approach (implementation scikit-learn.FastICA), where independent components are estimated by minimizing Gaussianity of the data (***15***). The assumption is that the two planes are a mixed observation of two clean sources and are mixed linearly using a mixing matrix. We assume a mixing matrix of the form [[1-a, a], [b, 1-b]], where a and b are in [0,1), and probably around 0.15. After FastICA, we transform the resulting mixing matrix to be of this form to recover the proper scaling of the mixed signals. Prior to FastICA, data undergoes whitening; we do not use the built-in whitening of the FastICA module (as it appears to contain a bug that affects scaling of the outputs).

#### Algorithm description

The plane whose traces we are correcting is referred to as the signal plane. The plane coupled to it (simultaneously acquired with temporal multiplexing) is referred to as the crosstalk plane. The algorithm is run on both permutations of signal plane and crosstalk plane, i.e., for each pair, it is run once with plane A as the signal plane and plane B as the crosstalk plane and once with plane B as the signal plane and plane A as the crosstalk plane.

Cell segmentation is performed on each plane independently to generate cell ROIs. We define the set of ROIs as those detected in the signal plane. We construct raw signal traces by measuring the average per-pixel flux in each ROI in the motion-corrected movie taken from the signal plane (trace extraction). We construct raw crosstalk traces by measuring the average per-pixel flux in the same ROIs in the motion-corrected movie taken from the crosstalk plane (these ROIs are those detected in the signal plane). For both the raw signal traces and the raw crosstalk traces, raw neuropil traces are constructed using the pixels bordering the ROIs. We flag any ROI whose footprint or neuropil intrudes on the motion correction border as invalid and remove it from further processing.

Signal and crosstalk planes are coupled and leak into to each other due the imperfections of the temporal multiplexing approach. We assume that, for each ROI, the measured, or raw signal and crosstalk traces are linear combinations of clean, or unmixed signal and crosstalk traces, i.e. R = M U where R is a 2xN matrix (N is the number of timesteps in the traces) such that R[0,:] is the raw signal trace and R[1,:] is the raw crosstalk trace. U is a 2xN matrix such that U[0,:] is the unmixed signal trace and U[1,:] is the unmixed crosstalk trace. M is the 2x2 mixing matrix relating the two. For each ROI, U[0,:] represents the trace coming from that ROI uncontaminated by crosstalk. That unmixed signal is the trace that we want to use in all future processing steps. We use Independent Component Analysis (ICA) [1] to solve for U and M. Specifically, we:

1. Subtract the mean from each trace and find the 2x2 matrix W which will transform (R-mean) into a “whitened” dataset whose correlation matrix is approximately the 2x2 identity matrix.
2. Use <u>scikit learn’s FastICA</u> algorithm to solve for U and M
3. The scale of the traces is corrected by computing the scaling that will restore the mixing matrix to the assumed form of [[1-alpha, alpha], [beta, 1-beta]]. This scaling can be computed by multiplying the inverse of the mixing matrix by the vector [1,1]. This transformation is applied to the traces to restore the original scale.
4. If the off-diagonal elements of M are positive and less than 0.3, ICA has converged to a valid result. Identify the unmixed trace that is most closely correlated with the raw signal trace as the unmixed signal trace (ICA is agnostic regarding the ordering of unmixed signals).
5. If ICA failed to converge to a valid result, the ROI is marked as having failed ICA.

We use all the ROIs that pass the above algorithm to construct an average mixing matrix M for the signal plane. The inverse of this matrix is used to unmix any ROIs marked as “failed” in step (4). We find unmixed neuropil traces by taking the mixing matrix used for the associated ROI and multiplying its inverse by the matrix of raw neuropil traces. If no ROIs in the plane converged to a valid result, it is impossible to construct an average mixing matrix for the plane and we mark the plane as having failed processing.

We run the unmixed signal and unmixed crosstalk traces through an event detection algorithm described in (*16*). We define events as a sequence of five timestamps whose sum of log probabilities exceeds a threshold (14 in our implementation). We calculate the probability relative to a Gaussian distribution centered on the mode trace value whose width is calculated considering the distribution of all trace values less than the mode. We define each timestamp that passes this test as an event in the trace.

For each ROI, we assess whether any events occur in the unmixed signal trace that are independent from the events in the unmixed crosstalk trace. We define independent events as those events in the unmixed signal trace which do not occur within two timestamps of an event in the unmixed crosstalk trace. If no such independent events occur in the unmixed signal trace, we mark the ROI as a ghost (i.e. an ROI that was only detected because of crosstalk contamination from the coupled plane) and discard it from future processing steps.

#### ROI Filtering

Not all ROIs generated by segmentation are complete individual cell bodies. To exclude ROIs that are not actually cell bodies from further analysis, the ROIs are labeled with a multi-label classifier that distinguishes ROIs that are considered well defined cell bodies from other ROIs. The set of reasons to exclude an ROI are: the ROI is a union of two or more cells; the ROI is a duplicate of another; the ROI is close to the edge of the FOV and is impacted by motion such that parts of the ROI are missing from the video; the ROI is likely an apical dendrite and not a cell body; or that the ROI is too small, too narrow, or too dim to confidently be considered a cell body.

The initial ROI filtering was generated by a set of heuristics based on depth, shape, area, intensity, signal-to-noise, and the ratio of mean to max intensity of the max projection. The initial filtering was used to generate a set of training labels, on which a multi-label classifier was trained. The multi-label classifier is implemented using a linear Support Vector Classifier trainer for each label (binary relevance) using metrics generated in segmentation combined with depth, driver, reporter, and targeted structure as features. The final ROI filtering workflow is to 1) label ROIs that fall within the motion cutoff regions at the border, 2) label ROIs using the binary relevance classifier, 3) label significantly overlapping ROIs as duplicates, and 4) label ROIs that significantly overlap two or more ROIs as unions.

#### Demixing Traces From Overlapping ROIS

The simplest way to extract fluorescence traces, given a set of ROI masks, is to average the fluorescence within each ROI. If two ROIs overlap, this procedure will artificially correlate their traces. Therefore, a model is used where every ROI has a trace which is distributed across its ROI in some spatially heterogeneous, time-dependent fashion:

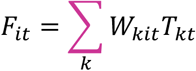

where *W* is a tensor containing time-dependent weighted masks: *W*_*kit*_ measures how much of neuron *k*’s fluorescence is contained in pixel *i* at time *t*. *T*_*kt*_ is the fluorescence trace of neuron *k* at time *t* - this is the desired value to estimate. *F*_*it*_ is the recorded fluorescence in pixel *i* at time *t*.

Importantly, this model applies to all ROIs, including those too small to be a neuron or otherwise filtered out.

Duplicate ROIs (defined as two ROIs with >70% overlap) and ROIs that are the union of two other ROIs (any ROI where the union of any other two ROIs accounts for 70% of its area) are filtered out before demixing, and the remaining filtering criteria are applied after demixing. Projecting the movie (*F*) onto the binary masks (*A*) reduces the dimensionality of the problem from 512x512 pixels to the number of ROIs:

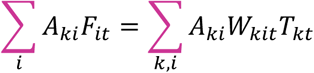

where *A*_*ki*_ is one if pixel *i* is in ROI *k* and zero otherwise–these are the ROI masks from segmentation, after filtering out duplicate and union ROIs. At a particular time point *t*, this yields the simple linear regression:

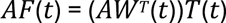

where the weighted masks *W* are estimated by the projection of the recorded fluorescence *F* onto the binary ROI masks *A*. On every imaging frame *t*, the linear least squares solution *T*^ are computed in to extract each ROI’s trace at that time point.

It was possible for ROIs to have negative or zero demixed traces *T*^. This occurred if there were union ROIs (one ROI composed of two neurons) or duplicate ROIs (two ROIs in the same location with approximately the same shape) that the initial detection missed. If this occurred, those ROIs and any that overlapped with them were removed from the experiment. This led to the loss of ∼1% of ROIs.

#### Neuropil Subtraction

The recorded fluorescence from an ROI was contaminated by the fluorescence of the neuropil immediately above and below the cell due to the point-spread function of the microscope. In order to correct for this contamination, the amount of contamination was estimated for each ROI. The estimated *F*_*N*_ was done by taking an annulus of 10 mm around the cellular ROI, excluding pixels from any other ROIs. In order to remove this contamination, the extent to which ROI was affected by its local neuropil signal was evaluated.

The recorded traces were modeled as *F*_*M*_ as *F*_*M*_ = *F*_*C*_ + *rF*_*N*_, where *F*_*C*_ is the unknown true ROI fluorescence trace and *F*_*N*_ is the fluorescence of the surrounding neuropil. In order to estimate the contamination ratio *r* for each ROI, the error was minimized *E* = 〈(*F*_*C*_ − (*F*_*M*_ − *rF*_*N*_))^2^ + *λ*Λ(*F*_*C*_)〉_*t*_ by jointly optimizing for *r* and *F*_*C*_. Λ(*F*_*C*_) is the first temporal derivate of the cellular trace, weighted by *λ* = 0.05; this smoothness constraint on the cellular trace allows per-ROI optimization for *r*. 〈·〉_*t*_ denotes an average over time. Gradient descent was used on *r*. At each step of the gradient descent, *F*_*C*_ was solved at the zero gradient of *E*. Gradient descent was performed on the first half of the traces and computed *E* on the second, so that it is a cross-validation error. After computing *r* and *F*_*C*_ for an ROI, the neuropil-subtracted trace *F*_*M*_ − *rF*_*N*_ was used as the basis for all subsequent analysis in order to avoid any residual effects of the smoothness constraint.

To standardize the learning rate and initial conditions of the gradient descent, each ROI’s neuropil trace was normalized to (0,1). The measured ROI trace was normalized by the same amount, used a learning rate of 10 and initial condition of *r* = 0.001. The gradient descent was stopped at the first local minimum of *E*. If the resulting *r* was greater than 1 or less than 0, or final cross-validation error *E* greater than 2 |〈*F*_*M*_〉_*t*_|, the gradient descent was attempted again with a 10x slower learning rate. If those convergence criteria were still not met, an initial condition of *r* = 0.5 was used. If those convergence criteria were still not met, that ROI was flagged and after computing *r* for all other ROIs in the experiment, set *r* for un-converged ROIs to the mean.

To validate the performance of our algorithm, it was tested on a publicly-available benchmark dataset (Chen et al., 2013). A distribution of contamination ratios was obtained, centered nearly on the author’s choice of 0.7 (mean r of 0.68 vs their choice of 0.7), but with significant heterogeneity. For this benchmark dataset and using the same optimization parameters as for the Allen Brain Observatory – Visual Coding data, 6 of 36 cells failed the initial neuropil subtraction with r>1, and would have gone through the additional steps outlined above.

#### dF/F Calculation

We generate normalized, detrended traces of neuronal activity by performing the following algorithm on each trace:

First, we estimate the standard deviation of the noise in the trace. We do this by centering the trace on the curve resulting from applying a median filter with kernel size 3.33 seconds to the trace. To avoid including signal events in our estimate of noise, we discard any values that exceed 1.5 times the absolute value of the minimum of this centered trace. We make a first estimate of the standard deviation of noise in the trace as 1.4826 times the median absolute deviation of this truncated, centered trace. We further discard any values that exceed 2.5 times this first estimate of the standard deviation and finally return 1.4826 times the median absolute deviation of the remaining centered trace values as the standard deviation of noise in the trace.

Next, we calculate the dF/F trace. We define the baseline activity in the trace as the result of applying a median filter with kernel size 600s to the trace. We subtract this baseline from the raw trace and normalize the difference by the baseline. At timesteps where the baseline trace is less than the standard deviation of noise calculated above, we normalize by the standard deviation of noise, instead. This gives the dF/F trace.

Finally, we detrend the dF/F trace. We estimate the trend in the dF/F trace by applying a median filter with kernel length 3.33 seconds. To prevent anomalously large trends from arising, we use the same algorithm we used to estimate the standard deviation of noise in the raw trace to estimate the standard deviation of noise in the dF/F trace. We constrain the trend to never exceed 2.5 times this estimated standard deviation of noise in the dF/F trace. We subtract the trend from the dF/F trace and report this as the detrended dF/F trace.

#### Calcium Event Detection

We used the <u>FastLZeroSpikeInference</u> (“FastLZero”) to identify events in traces derived from the 2-photon imaging movies (*17*). FastLZero fits a trace to a sum of exponentially decaying spikes, with the timescale of decay determined by a single parameter (gamma), while imposing an L0 regularization to minimize the number of events used to fit. Gamma has been empirically determined for each genetically defined cell type by running the MLSpike autocalibration routine on all available recordings and determining the maximum likelihood decay constant across the distribution of fitted decay constants for available neurons of any given cell type (*18*).

The relative weight in the optimization penalty between number of events used and best residuals is controlled by a regularization factor (lambda) that controls the tradeoff between missing events (false negatives) and overfitting of noise with extraneous events (false positives). Lambda needs to be determined empirically as a function of the noise level present in each recording. We iteratively searched the regularization factor space to find the regularization factor where the smallest event provided by FastLZero was at least two times the estimated noise level of the fluorescence trace. The noise estimator was initially optimized for experiments recorded at 31Hz (single-plane imaging with Scientifica 2-photon microscopes) and validated against ground truth data manually annotated by a human expert.

Multiscope experiments were recorded at 11Hz, and we observed that the same multiplicative factor (2.0) resulted in many more small amplitude events than in the 31Hz data when probed with synthetic calcium data. One can rationalize this difference intuitively by noting that the characteristic timescales of fluorescence decay do not change based on sampling rate, but the noise estimation does. Thus, our noise estimator, which was only validated for 31Hz data, systematically underestimated the noise for 11Hz data. To correct for the effects of sampling rate on the noise estimation, for Multiscope, we empirically determined the equivalent multiplicative factor to be 2.6 by identifying the value that minimized the discrepancy between event magnitude traces extracted (using factor of 2.0) from Scientifica data sampled at 31 Hz, and event magnitude traces extracted from the same data but after downsampling by a factor of 3.

### Session to Session Cell Matching

Multiple 2-photon calcium imaging movies were acquired for each imaging plane across multiple imaging sessions. To map cells between sessions, we used an automated matching algorithm. The module has 4 steps:

1. Determining the spatial transform between each pair of sessions using image registration techniques on the average projection images of the sessions.
2. Applying the derived spatial transform to segmented cell masks.
3. For each pair of sessions, solving the linear assignment problem (i.e. bipartite graph matching) to determine which pairs of masks are most likely to be related, based on the inter-centroid distance of the segmented masks and the intersection over union (IOU) of the masks. A 10 pixel distance threshold is placed on the inter-centroid distance to ensure only closely overlapping masks were matched.
4. A graph combination method was used to join all the bipartite graphs to determine the most likely label sets that match across all sessions.

Code for pairwise session cell matching is available on GitHub.

#### Determining the spatial transformation

The module first used an intensity-based method to register the average intensity projection images of each pair of sessions, producing a Euclidean transformation that registered each image pair. We found that a single image registration strategy did not work for all cases. In some cases, the true lateral shifts between two experiments were too large for the ECC registration algorithm. In other cases, the border in the projection images biased the registration towards matching borders and not cell soma. In still other experiments, a histogram equalization step that helped most experiments resulted in a degraded registration.

We implemented a “meta-registration” where 4 different registration sequences were attempted. Each sequence employed an initial cropping step to eliminate the motion border problem. The sequences employed varying combinations of ECC, PhaseCorrelation, and contrast adjustment. After each of the 4 sequences were attempted, we chose the best candidate based on the structural similarity metric (SSIM) between the two registered images. If none of the candidate sequences improved SSIM beyond that achieved with unregistered images, we flagged the registration as a failure for QC inspection.

In a few cases, there were still some unexplained failures to register. In these cases, we found that the average projection intensity images had low contrast, but the maximum projection intensity images had more visible features. By substituting in the maximum projections for the registration step, we achieved good registration and good cell matching for these containers.

#### Applying the spatial transformation to cell masks

The cell masks were materialized into a set of images, such that no ROIs overlapped in any one image. We then applied the relative transformation found from the intensity projections to these materialized images, to get the cell masks in an aligned space.

#### Solving the linear assignment problem between 2 sessions

To map cells, a bipartite graph matching algorithm (the Blossom method) was used to find correspondence of cells between sessions. The algorithm used cell labels in the pair-wise experiments as nodes, and the edges of the graph were weighted with weights, *w*:

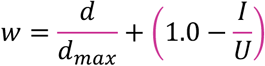

where *d* is the distance between the 2 mask centroids (in pixels), *d_max_*_’_= 10 pixels, *I* is the number of pixels that are shared between the 2 masks (the intersection), and *U* is the total number of pixels covered by the two masks combined (the union). The graph edge for any pair of masks with *d* > 10 pixels was assigned a very large weight, eliminating that pair as a candidate match.

The original version of this module used the “Hungarian” method to minimize the total weight.

SciPy has an <u>equivalent implementation</u> of that method. We found that the solution from the “Blossom” method was more robust to order permutations of the inputs. We used the <u>networkx implementation</u> of the “Blossom” method. As this implementation seeks to maximize total weight, we inverted the weights, taking care to prevent division-by-zero:

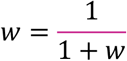

By maximizing the summed weights of edges, the bipartite matching algorithm found the “best match” between cells in each pair of experiments.

#### Joining the pairwise bipartite graphs

All the pairwise bipartite graphs were naively combined. This combined graph is not constrained to avoid labeling conflicts. Matching was performed on the full set of ROIs identified by the segmentation algorithm, but ROIs that were deemed invalid by the ROI filtering step were not included in the final set of matched cells.

For quality control checks, we produced standard plots of pairwise registrations across sessions, as well as quantification of the fraction of ROIs matched across sessions. Session pairs with visible failures to register (clear features in the image were not aligned), or with very low fraction of matched cells (<10% matching between session pairs), were manually evaluated and an attempt was made to optimize the registration process to produce accurate registration. Session pairs that were not possible to register were excluded from the dataset.

The code for performing field of view registration and cell matching is tested in a CI/CD system and openly available at: https://github.com/AllenInstitute/ophys_nway_matching

### Eye Tracking Data Processing

A standardized pipeline was built for fitting ellipses to the pupil, eye (visible perimeter of the eyeball), and corneal reflection of the right eye, based on points tracked using the open source software DeepLabCut (https://github.com/DeepLabCut/DeepLabCut). We used DeepLabCut, initialized with a pre-trained ResNet 50 deep residual network, to track (up to) 12 points along the perimeters of the eye, pupil, and corneal reflection.

Ellipses were then fit to the tracking points and the ellipse fit parameters were saved to disk.

Validation against hand-annotated ‘ground truth’ frames confirmed that a single ‘universal’ model, trained on a broad selection of data samples, robustly generalized on held-out data across different physiology rigs and individual animals.

All code for processing of the eye tracking data is visible in the <u>brain_observatory.behavior.eye_tracking_processing</u> module of the AllenSDK. The eye tracking data is available as the ‘eye_tracking’ dataframe in the session object. The dataframe has the following columns:

- Timestamps: the timestamp of every frame
- {pupil, eye or cr}_center_x: the x-position of the center of the pupil, eye or corneal reflection ellipse fit on that frame, in pixel space of the eye tracking movie frame.
- {pupil, eye or cr}_center_y: the y-position of the center of the pupil, eye or corneal reflection ellipse fit on that frame, in pixel space of the eye tracking movie frame.
- {pupil, eye or cr}_width: the (more) horizontal half-axis of the ellipse fit of the pupil, eye or corneal reflection
- {pupil, eye or cr}_height: the (more) vertical half-axis of the ellipse fit of the pupil, eye or corneal reflection
- {pupil, eye or cr}_phi: the angle to the (more)horizontal axis of the ellipse fit of the pupil, eye or corneal reflection, measured CCW from the x-axis of the image frame.
- likely_blink: a Boolean defining frames that have been identified as likely outlier fits, which is often caused by blinking/squinting of the eye.
- pupil_area: the area of the pupil, assuming that the pupil is a circle with a diameter defined by the major axis of the ellipse fit. Set to NaN where likely_blink == True.
- pupil_area_raw: the area of the pupil, assuming that the pupil is a circle with a diameter defined by the major axis of the ellipse fit. No outliers/likely blinks are removed.
- {eye or cr}_area: the area of the eye or corneal reflection ellipse fit. Set to NaN where likely_blink == True.
- {eye or cr}_area_raw: the area of the eye or corneal reflection ellipse fit. No outliers/likely blinks are removed.

Briefly, the ellipse fit parameters produced by the processing pipeline were loaded for each session. It was assumed that the pupil is actually round, but when viewed obliquely, it appears as an ellipse, the major axis of which reflects the pupil diameter. Thus, the area of the pupil is calculated on every frame as the area of a circle with a diameter defined by the longest axis of the ellipse on that frame. The area of the ellipses for both the corneal reflection and eye ellipse fits are calculated using the standard formula for the area of an ellipse.

On frames where the animal is blinking, tracked points may be missing or the confidence of tracked points can be low, and ellipse fits either fail completely or may be fit erroneously. To avoid including these erroneous fits in analysis, an algorithm attempts to identify these blink frames, adding a column to the eye_tracking dataframe called ‘likely_blink’. Likely blinks are identified as frames where either the eye or pupil fit is missing, or where the z-scored value of the eye or pupil areas exceeds 3. In addition, two frames before and after every likely_blink identified by the above methods are also labeled as likely blinks. This is to avoid the possibility of analyzing erroneous fits caused by a partially opened eye.

It is important to note that the outlier frames identified by the likely_blink algorithm are not always caused by blinks. Instead, they may be frames where the DeepLabCut algorithm simply failed to identify a reasonable fit, triggering the outlier detection portion of the algorithm.

### Behavior data analysis

#### Behavior performance

Reward rate was computed as the average rewards per second in a rolling 320 second window, over each 60-minute behavior session. Engaged periods of the session were defined as having >2/3 rewards per minute.

Response probability was computed as the fraction of image presentations for a given condition where the mouse emitted a licking response in a 750 ms window after event onset. Conditions included stimulus changes, non-change image presentations that could have been a change, image omissions, or image presentations following omissions (fig. S3B, E; fig. S4B). Response probability for non-changes (i.e. false alarm rate) was computed as the fraction of stimulus presentations that could have been a change (i.e. not during reward window, reward consumption “grace period”, or 2 second period prior to drawing a change time) when an incorrect response was emitted.

The value of <u>d-prime</u>, also known as the discriminability or sensitivity index, was calculated as the relative difference between the inverse cumulative normal distribution function of the hit rates and false alarm rates. Hit and false alarm rates were computed as described above (hit rate as response probability for image changes and false alarm rate as response probability for stimulus presentations that could have been a change but did not change). The mean d-prime was computed for all image transitions for each behavior session, regardless of engagement state. The max d-prime was used for some analyses to assess peak performance, independent of task engagement. The distribution of average d-prime values is shown for all trainings stages in fig., S3J, for individual cohorts in fig. S4E-H, and as the max d-prime value across each session for Fig. 2F.

#### Normalized pupil width

Pupil width, defined as the horizontal half-axis of the ellipse fit of the pupil (described above in “Eye Tracking Data Processing”), was <u>pre-processed</u> by filtering out “likely blinks” and linearly interpolating values. Pupil width was then normalized to the 5-minute gray screen period at the beginning of the session to account for any differences in pupil area resulting from slightly different placement of the dichroic mirror reflecting the eye to the eye tracking camera. This allowed a direct comparison of normalized pupil width across Familiar, Novel, and Novel+ sessions in Fig. 2H and fig. S5B,E,H,K. Normalized pupil width was aligned to the time of image changes or image omissions within each session and averaged, then averaged across all sessions of a given type.

#### Lick rate

For each session, the lick rate was <u>computed</u> as a rolling average of the number of licks in a 100ms window (6 frames at 60Hz acquisition rate) to give a continuous timeseries in units of licks / 0.1 second, then multiplied by 10 to give licks / 1 second. The rolling lick rate was aligned to the time of image changes or image omissions within each session, then averaged across sessions and mice, to produce the average lick rate at each timepoint relative to changes or omissions, as shown in fig. S5C,F,I,L.

#### Temporal alignment of 2-photon imaging and behavior

Temporal alignment is needed to link and analyze data streams collected at different frame rates. In our experiments, 2-photon movie timeseries collected on Scientifica microscopes for single-plane imaging were collected at 30Hz. 2-photon movie timeseries collected on the Multiplane Mesoscope were collected at 11Hz. Running speed, licking, and reward delivery were collected at 30Hz, at the frequency of the stimulus display. Eye tracking movies were collected at 60Hz.

To generate stimulus locked traces of neural activity, we extracted a subset of each cell’s dF/F trace (or detected events) in a window around each stimulus onset time. To prevent shifting of timestamps to align 2-photon data to stimulus times, we interpolated neural activity traces to a 30Hz sampling rate relative to the stimulus onset time for each trial. This allowed more accurate averaging of neural signals across trials, as each trial’s timestamps were interpolated relative to the stimulus start time. A <u>demonstration of the time alignment procedure can be found on GitHub</u> along with the <u>functions to</u> <u>perform the alignment</u>.

### Response characterization

All response analyses were performed using detected calcium events. To remove aliasing between the neural activity timestamps and stimulus timestamps all continuous signals were interpolated onto stimulus-aligned timestamps, as described above.

#### Cell response metrics

For each 2-photon session, event triggered responses were extracted for each cell, in a window around the time of stimulus changes, repeated stimulus presentations, and stimulus omissions.

To produce a population average timeseries across all neurons within a given cell type, image change, repeated image, or image omission aligned traces were aggregated over a given condition (ex: change aligned or omission aligned) and averaged for each neuron (as shown in Fig. 3F-H,M), then averaged across all sessions of a given type (Familiar, Novel, Novel+) and displayed +/-SEM in Fig. 3B-D, I-K. The population average omission response is shown for active and passive behavior sessions with familiar stimuli in fig. S9C.

In addition to a segment of each neuron’s trace in a window around the stimulus (or omission) onset time, we computed the mean response in a 0.5 second window after stimulus (or omission) onset. For each stimulus condition (changes, repeated stimuli, and omissions), we created a distribution of mean response values across neurons for each experience level (Familiar, Novel, Novel+) and report the mean +/-95% CI in fig. S7A-D and fig. S7A-F.

The significance of the response to each stimulus presentation (or omission) was computed by comparing the mean response in the 0.5 second window after stimulus (or omission) onset with the mean response in a 0.5 second window taken from a shuffled distribution of that neuron’s activity during the gray screen spontaneous activity period where no stimulus was present. A p-value was calculated by comparing these values across 10,000 shuffles and computing the fraction of those shuffles where the mean stimulus response was larger than the mean response in the shuffled data. A neuron was considered “responsive” for a specific condition if at least 25% of trials for that condition (ex: changes, omissions) had a p-value < 0.05 (p-value determined by comparing each trial’s mean response to a shuffled distribution as described above in “trial level metrics” section). The fraction of responsive neurons for each session was then calculated as the number of “responsive” neurons divided by the total number of neurons, and is shown in Fig. 3E,L and fig. S7E-F, and fig. S8K.

A modulation index was computed to quantify single cell modulation by image changes and image omissions. For each cell, we computed the mean response in a 500ms window following image changes, the mean response in a 500ms window following onset of the repeated, pre-change stimulus, then took the difference over the sum of these values, producing a metric with positive values indicating stronger responses to changes compared to pre-change images, and negative values indicating stronger responses to pre-change images compared to changes. Results are shown in fig. S7I-L. Similarly, an omission modulation index was computed by comparing the mean response following image changes compared with the 500ms gray screen period prior to the time of the image omission, with results shown in fig. S8I-K.

An experience modulation index was computed for cells matched across pairs of sessions, as the difference over the sum of each cell’s average change response in one session relative to another (ex: mean response to Novel image changes versus Familiar image changes), or the difference over the sum of each cell’s average omission response across sessions. Distributions of this metric over matched cells for each pair of sessions are shown in fig. S7M,N and fig. S8L,M.

Code for all cell response metrics can be found on GitHub.

#### Statistics for response metric comparisons

For all metrics, to test whether response distributions were significantly different across experience levels, areas, or depths, we used one-way ANOVA, followed by Tukey HSD to compare population means across the conditions being compared. A p-value of 0.05 was used as the significance threshold.

#### Stimulus locked behavior timeseries

For behavioral timeseries, the stimulus aligned traces for all trials of a given condition were averaged for each session, then averaged across sessions, and displayed +/- SEM in fig. S5.

### Population decoding

#### General methodology for decoding population activity

A linear SVM (Python scikit-learn package) was trained on each bin of the population activity in each session. Population activity was aligned on either the image changes or omissions (fig. S7G,H; fig. S8G,H; fig. S9A,B). Time bins within the window [-0.5 0.75] sec surrounding the image change (or omission) were used for classification. One classifier was trained per time bin (non-overlapping 93ms time bins for Mesoscope experiments, and 33ms time bins for Scientifica experiments). Decoding analysis was performed on the full population of neurons recorded in each session, and also on the “cell matched” population (fig. S6). To obtain the “cell matched” population, neurons were matched across 3 sessions: the last Familiar session, the 1st Novel session, and the 2nd Novel session. Similar results were found from both populations (full and cell matched).

Details of the decoding method are explained in (*11*). In brief, to break any dependencies on the sequence of trials, we shuffled the order of trials for the entire population. L2 regularization was used to avoid over-fitting. 5-fold cross validation was performed by leaving out a random 1/5 subset of trials to test the classifier performance, and using the remaining trials for training the classifier. This procedure was repeated 50 times. A range of regularization values was tested, and the one that gave the smallest error on the validation dataset was chosen as the optimal regularization parameter. Classifier accuracy was computed as the percentage of testing trials in which the class was accurately predicted by the classifier and summarized as the average across the 50 repetitions of trial subsampling. A minimum of 10 trials and 3 neurons was required to run the SVM on a session. The inferred spiking activity of each neuron was z-scored before running the SVM.

#### Decoding image change occurrence

To address whether population activity in single trials carries a signal about image changes, population activity following an image change was classified against the activity following a non-change image (fig. S7G,H). For the non-change image, the image immediately preceding the image change was used. One SVM was trained per time bin in the window [-0.5 0.75] sec surrounding the pre-change image and image change. The input to each SVM was formed in the following way: population activity at time t relative to each image change (class 1; size: n x m, where n: number of image changes; m: number of neurons) was vertically concatenated with the population activity at time t relative to each pre-change image (class 0). This was done for all image changes and pre-change images in the session (SVM input size: 2*n x m, where n: number of image changes; m: number of neurons), allowing to study how the population activity at time t after an image change was distinct from the activity at time t after a non-change image. The output of the SVM constituted of two classes, representing image changes and no changes.

#### Decoding omission occurrence

To address whether population activity in single trials carries a signal about omissions, population activity following an omission was classified against the activity during the gray screen (fig. S8G,H). Gray screen activity was taken from the time bin immediately preceding the omission. One SVM was trained per time bin in the window [-0.5 0.75] sec surrounding the omission. The input to each SVM was formed in the following way: population activity at time t relative to each omission (class 1) was concatenated with the population activity at time t-1 relative to each omission (class 0, gray-screen activity). This was done for all omissions in the session (SVM input size: 2*n x m, where n: number of omissions; m: number of neurons). The output of the SVM constituted of two classes, representing omissions and no omissions, i.e., gray screens.

#### Decoding image identity from omission-evoked activity

To address whether population activity following omissions carries information about the identity of the subsequent image, population activity following an omission was classified according to the post-omission image identity (fig. S9A,B). One SVM was trained per time bin in the window [-0.5 0.75] sec surrounding the omission. The input to each SVM was formed in the following way: population activity at time t relative to omission was concatenated for all omissions (SVM input size: n x m, where n: number of omissions; m: number of neurons). The output of the SVM constituted of 8 classes, representing the 8 different images that followed the omission in each session.

#### Summary quantification of decoding results

To quantify the magnitude of population decoding accuracy, the decoding accuracy trace of each experiment was averaged over 400 ms after image (or omission) onset for decoding image changes, image identity, and post-omission image identity, and over 750ms after omission onset for omission decoding analysis. This quantity was averaged across all experiments collected using both single and multi-plane two-photon imaging (fig. S7G; fig. S8G; fig S9A).

#### Statistical tests

We used two-way ANOVA, followed by Tukey HSD to compare population decoding accuracies across experience levels (Familiar, Novel 1, Novel+). Two-sided t-test was used to compare correlations between the real and shuffled data, for each experience level. A p-value of 0.05 was used as the significance threshold.

### Regression model

Our regression model is a linear model with time-dependent kernels with a gaussian noise model.

Each model feature is a vector *f_i_*(*t*), and was convolved with a learned kernel *k_i_*(*t*) to produce the predicted model component of that feature *f_i_*(*t*) ∗ *k_i_*(*t*). Model components were summed together to produce the full model response *r*(*t*) = ∑*_i_f_i_*(*t*) ∗ *k_i_*(*t*). The learned kernels were instantiated as vectors of weights. The length of each feature kernel was determined using domain knowledge and trial and error. Image feature vectors were 0.75 seconds in length following the time of each stimulus presentation, and were aligned to all image presentations including image changes. The omission feature was 3 seconds in length following the time of the expected stimulus presentation. Hit and miss features were 2.25 seconds in length after the time of image changes, separately for trials where the mouse licked correctly and was rewarded (hits) and trials where no response was emitted (misses). Lick feature vectors were 2 seconds in length starting 1 second before the start of each lick. For continuous features (running speed and pupil diameter) *f_i_*(*t*) was a time-series containing the data series at each time point, the kernel was 2 seconds in length aligned to a +/- 1 second window around each time point. For discrete features *f_i_*(*t*) was 1 on timesteps where that feature occurred, and 0 elsewhere.

#### Stimulus pre-processing

For each cell, we fit the model to detected calcium events smoothed with a causal half-gaussian filter with scale 65ms. This smoothing step was performed in order to fit the model to a continuous signal. To remove aliasing between the neural activity timestamps and feature timestamps all continuous signals were interpolated onto stimulus-aligned timestamps. Discrete features were binned onto the nearest stimulus-aligned timestamp. For a given experiment, if a feature had less than 5 discrete events, that feature was not included in the model. Continuous features (pupil diameter and running speed) were standardized to have mean zero and unit variance.

#### Toeplitz Matrix Implementation

Convolutions are time-invariant linear shift operators, which allowed us to implement the convolution of kernels with feature timeseries as a banded toeplitz matrix operation *y* = *Wx*. Here *y* is the 1d time-series for a cell, *x* is a vector of all kernel weights concatenated together, and W is a toeplitz matrix with diagonal bands that map kernel weights onto time-shifted features.

#### Closed form solution and ridge regularization

The toeplitz matrix implementation with a gaussian noise model yields the standard solution for ordinary least squares regression: *x* = (*W^T^W*)^-1^*W^T^y*. We added an L2 ridge regression penalty to the cost function, resulting in the following solution: *x* = (*W^T^W* + *λI*)^-1^*W^T^y* where *λ* is the L2 penalty. We fit the model using 5-fold cross validation. We split each session into 50 intervals, and randomly assigned 10 intervals to each cross-validation fold. Therefore, each of the 5 folds were intermingled in time. To determine the hyper-parameter *λ* we evaluated the model on a grid of potential *λ* values from 0 to 500, and for each experiment we selected the *λ* that resulted in the best test-set performance across cells in that experiment. The training/test splits were different for hyper-parameter selection than fitting the model for analysis.

#### Model evaluation

We evaluate our model by computing the explained variance: 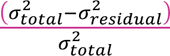. If the smoothed calcium event trace had no activity (no events) on a cross-validation fold, then then cell has no variance on that fold. Therefore, for numerical stability, when a cell had no activity on a cross-validation fold, the model explained variance was set to 0.

#### Unique contribution of model features

To evaluate the contribution of each kernel in explaining each cell’s activity we fit a series of reduced models where individual kernels or groups of kernels (components) were removed from the model and the model was refit. We compute a coding score which measures the fraction of the full model’s explained variance that is lost when using the reduced model. This coding score therefore captures the unique explained variance from that kernel or set of kernels. The raw coding score for feature *i* is given by: 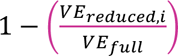. Some model features are only present during a portion of the session, we therefore adjust the raw coding score to only evaluate the explained variance on the time points when that feature was present. The adjusted coding score for feature *i* is given by: 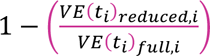. Here *t_i_* indicates timepoints where feature *i* was in the model, and *VE*(*t*_*i*_) is the explained variance only on the timepoints where feature *i* was in the model. For numerical stability, the coding scores were bounded between (0,1). Further, if the full model explained less than 0.5% of the variance on the relevant time points, the coding score was set to 0. In fig. S10E we also compute coding scores for reduced models where all features were removed except for the feature of interest. In this case the coding score is computed by: 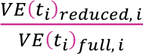.

#### Statistical tests on coding scores

We used one-way ANOVA, followed by Tukey HSD to compare population coding scores across experience levels (Familiar, Novel, Novel +). A p-value of 0.05 was used as the significance threshold.

Code for regression model can be found on GitHub. Functional clustering

#### Data Selection

For the functional clustering analysis in Figure 5 we selected neurons that were successfully matched across sessions with each of the three experience levels (Familiar, Novel, Novel+; fig. S4, fig. S6), regardless of their overall activity level or encoding strength. Feature coding of each of these neurons was represented by a 12-element vector of coding scores describing the four main model components (images, behavioral, omissions, and task features) across three experience levels (Familiar, Novel, Novel+).

#### Across Session Normalization of Coding Scores

Our functional clustering analysis is restricted to cells recorded across all three experience levels and specifically seeks to assess coding changes across sessions. We therefore normalized the coding scores across sessions to account for changes in the overall full model explained variance for a single cell across sessions. Specifically, the fractional change in variance explained (i.e. the coding score) of each feature in each session was quantified relative to the maximum overall variance explained across all 3 sessions for a given matched cell.

Since model features could be present for a variable number of timepoints across sessions, we normalize the explained variance into explained variance per relevant timepoint. Here, *VE*(*t_i_*)*_full, i, k_* is the explained variance on session k, on timepoints when model feature *i* was present. Likewise *VE*(*t_i_*)*_reduced, i, k_* is the explained variance on session k, on the same timepoints when model feature *i* was removed. *n_i,k_* is the number of timepoints when model feature *i* was present in session *k*. The full across session coding score is then given by:

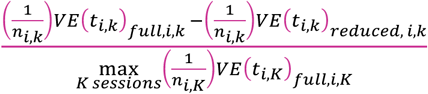

#### Spectral Clustering

We performed a clustering workflow that used spectral clustering in combination with a consensus clustering method (*19*) to isolate robust clusters that are not influenced by random initial conditions. In spectral clustering, the clusters are determined by their ‘connectivity matrix’, where nearby points (nodes in the graph) are assigned to the same cluster. We computed the connectivity matrix using a radial basis function. Next, the data was projected into a low-dimensional space using Laplacian embedding, and then each datapoint was assigned a label using the K-means strategy.

This spectral clustering algorithm was run 150 times with optimal k clusters determined as described below (see ‘Selecting Optimal Number of Clusters’). To perform consensus clustering on these 150 iterations, we computed a co-clustering association matrix and then calculated the probability of any given neuron being assigned to the same cluster with every other neuron. Lastly, we applied hierarchical agglomerative clustering to the association matrix to obtain final cluster labels. We used the <u>scikit-learn</u> library for clustering analysis.

#### Selecting Optimal Number of Clusters

We estimated the optimal number of clusters for all matched imaged neurons using the gap statistic method. To compute the gap statistic, we shuffled the coding features to create a null distribution. In this shuffle, the coding scores for each regressor and experience level are shuffled within their own category across cells, which preserves the original distributions of coding scores but breaks structure across days and regressors. Gap statistics were computed by clustering both shuffled and original data 20 times for k clusters range from k = 2 to k = 25. For each k number of clusters, we computed the mean of within cluster variability as measured by pairwise Euclidean distance for cells in each cluster: 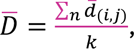, where 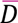 is the grand mean of means of pairwise distances, 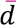, between points *i* and *j* in each *n* cluster with total k clusters. The peak difference between reference (null hypothesis based on shuffled data) and measured (alternative hypothesis based on original data) values indicates the largest reduction in data variability due to clustering (fig. S18a). To obtain eigengap values, we computed the eigenvalues of the Laplacian graph(*20*) (fig. S16A). We used the last non-zero peak of the sorted eigenvalue differences as an optimal number of clusters. Based on inspection of the gap statistic results we determined an optimal number of clusters (k=14). Two clusters contained were than five cells of each cell type, thus they were dropped due to small sample size, resulting in 12 clusters total.

#### Cell ID Shuffle

To investigate whether the functional clusters that we obtained from our dataset are cell specific or are a property of coding score distributions across days, we performed a cell id shuffle analysis (fig. S18). By shuffling cell ID matching across days while preserving coding scores within experience level, we perturbed within cell coding changes across days while maintaining overall effects of novelty at a population level. To maintain cell type specific information, we shuffled cell ids only within each cell type, such that coding scores remained cell type specific. We then followed the same clustering steps as described above using predetermined number of clusters (N=12, fig. S18), iterating the shuffling procedure 500 times. On each iteration, post-shuffle clusters and original clusters were matched using lowest sum of squared error value (SSE, fig. S18C). If all SSE values exceeded maximum threshold of 0.1 for a match, we concluded that a given original cluster was not found among post-shuffled clusters, and the matched cluster size was set to zero. We then compared mean size of matched post-shuffled clusters with the size of the original clusters to determine the pattern of coding across days in each cluster is equally likely to obtain by chance. Cluster size differences were marked as significant post chi-squared-test with Benjamini-Hochberg correction.

#### Distribution of Clusters Across Cortical Depth and Areas

To determine whether each neural cluster was evenly distributed across cortical area (VISp or VISl) or cortical depths (75, 175, 275, 375 um imaging depths, fig. S20-22G,H), we performed a chi-squared test to compare the difference between observed and expected proportions of cells in each cluster. We used the Benjamini-Hochberg procedure to correct for multiple comparisons. To compute proportions, we first normalized number of cells in each cluster and location to the total number of cells in each location across clusters. Expected proportion of cells per cluster was computed as an average of normalized observed proportions for each cluster.

Code for clustering analyses can be found on GitHub.

